# Left-right-alternating theta sweeps in the entorhinal-hippocampal spatial map

**DOI:** 10.1101/2024.05.16.594473

**Authors:** Abraham Z. Vollan, Richard J. Gardner, May-Britt Moser, Edvard I. Moser

**Author notes:** Equal contribution (Abraham Z. Vollan, Richard J. Gardner). Corresponding authors. contact: Abraham Z. Vollan,; Edvard I. Moser, Editorial correspondence.

## Abstract

Place cells in the hippocampus and grid cells in the entorhinal cortex are elements of a neural map of self-position^1–5^. To benefit navigation, this representation must be dynamically related to surrounding locations^2^. A candidate mechanism for linking places along an animal’s path has been described in place cells, where the sequence of spikes within each cycle of the hippocampal theta oscillation encodes a trajectory from the animal’s current location towards upcoming locations^6–8^. In mazes that bifurcate, such trajectories alternately traverse the two upcoming arms as the animal approaches the choice point^9,10^, raising the possibility that the trajectories express available forward paths encoded on previous trials^10^. However, to bridge the animal’s path with the wider environment, beyond places previously or subsequently visited, an experience-independent spatial sampling mechanism might be required. Here we show in freely moving rats, that within individual theta cycles, ensembles of grid cells and place cells encode a position signal that sweeps linearly outwards from the animal’s location into the ambient environment, with sweep direction alternating stereotypically between left and right across successive theta cycles. These sweeps were accompanied by, and aligned with, a similarly alternating directional signal in a discrete population of parasubiculum cells with putative connections to grid cells via conjunctive grid×direction cells. Sweeps extended into never-visited locations that were inaccessible to the animal and persisted during REM sleep. Sweep directions could be explained by an algorithm that maximizes cumulative coverage of surrounding space. The sustained and unconditional expression of theta-patterned left-right-alternating sweeps in the entorhinal-hippocampal positioning system provides an efficient ‘look-around’ mechanism for sampling locations beyond the travelled path.

## Main text

Grid cells are position-tuned cells whose firing locations rigidly tile the environment in a hexagonal lattice pattern^4,11,12^. Two decades of investigation have pointed to a mechanism for grid cells in which their firing emerges as activity is translated across an internally generated toroidal continuous attractor manifold, based on external speed and direction inputs^5,11–19^. However, the role of grid cells in navigation remains poorly understood. One clue is that the spatial coordinate system defined by grid cells allows position offsets to be expressed as vectors^11,13–15^. Not only does this make it possible to continuously update the neural position representation by path integration; it also allows the network to relate the current position estimate to target locations such as goals^20–24^. Grid cells have been suggested in computational models to dynamically probe the surrounding environment by way of linear “look-ahead” trajectories^21–25^. These proposed trajectories are reminiscent of the theta-paced forward sweeps recorded in hippocampal place-cell ensembles when animals run on linear tracks or mazes^6–10^. The sweeps observed on tracks and in mazes have limited navigational utility, however, since they are constrained to the travelled path. Here, recording from medial entorhinal cortex (MEC) and hippocampus with high-site-count Neuropixels probes^26,27^, we searched for a more general and experience-independent mechanism for rapid sampling of the ambient environment, including places never visited, in the population activity of many hundreds of grid, place and direction cells.

### Grid cells and place cells sample ambient space with alternating “sweeps”

Neural activity was recorded in 16 rats with Neuropixels probes targeting MEC and parasubiculum (384 to 1,559 cells per session; Extended Data Fig. 1) while the rats foraged for scattered food in an open field arena of 1.5m × 1.5m. As expected, the activity was patterned by the theta rhythm (Fig. 1a, top), which discretizes population activity in MEC and hippocampus into successive packets of ∼125 ms^28–30^. To examine the dynamics of spatial coding within individual cycles of the rhythm, we decoded position in 10-ms bins by correlating the instantaneous firing-rate population vectors from all MEC-parasubiculum cells with the session-averaged population vector at each position of the environment (i.e. the stack of firing-rate maps). Over the course of each theta cycle, the decoded position generally swept in a straight trajectory outwards from the location of the animal into the nearby environment (Fig. 1a, Supplementary Video 1). In general, such trajectories, referred to as “sweeps”, moved forwards at an angle from the animal’s head axis, with direction alternating between left and right on successive theta cycles. This pattern of left-right alternating sweeps was particularly prominent during periods of fast, straight running (Fig. 1a).

**Figure 1.**
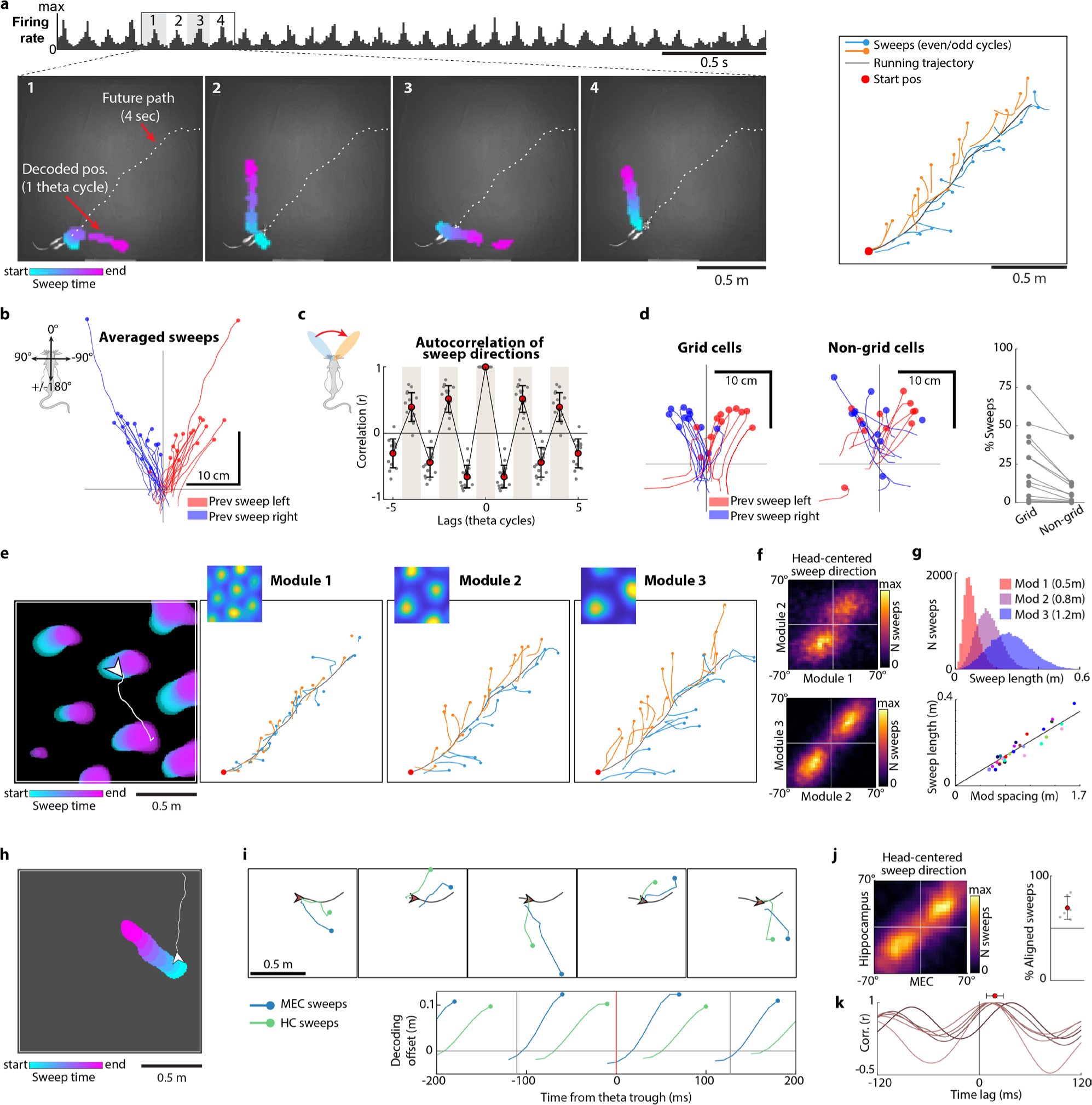
Grid cells and place cells sample ambient space with alternating “sweeps”. **a.** Theta-paced sweeps in ensembles of MEC and parasubiculum cells. Top: Summed spike counts from co-recorded MEC and PaS cells during a 3.5s epoch, showing 8-10 Hz theta-rhythmic population activity (Rat 25843, session 1). Bottom: Sweeps decoded from joint activity of all MEC-parasubiculum cells during the four successive theta cycles highlighted above. Panels show snapshots of the recording arena at the beginning of each theta cycle. White dashed line shows animal’s future trajectory. Decoded position throughout each theta cycle is plotted as colored blobs (position bins where correlation values exceeded the 99th percentile) with color indicating time within sweep. Note that sweeps progress outwards from the animal and are directed to the animal’s left and right side in alternation. Right: Decoded sweeps during the whole 3.5s period shown in top panel (one sweep per theta cycle). Each sweep is plotted as a line with colors corresponding to even and odd theta cycles. **b.** Sweeps rotated to head-centered coordinates (head orientation is vertical) and averaged across theta cycles where the preceding sweep was directed to the left (red) or to the right (blue). Data from 16 animals is shown (one pair of red/blue lines per animal). Note that the decoded position sweeps forward to alternating sides, in the opposite direction of the previous sweep (no red sweeps on left side, no blue sweeps on right side). **c.** Sweeps systematically alternate between left and right. Temporal autocorrelograms of angles between successive sweep directions, discretized in theta cycles, across all recording sessions. Grey dots show values for individual animals (n=16). Mean and s.d. are indicated by red dots with whiskers. Theta cycles are indicated by alternation of grey and white background shading. Note peaks at every other theta cycle, indicating rhythmic alternation from side to side. **d.** Sweeps are strongly expressed by grid cells. Left and center: Position was decoded separately from grid cells or a size-matched random selection of non-grid cells, repeated 100 times, within the recording region from the same session in the same animal. Plots show averaged decoded position for grid cells (left) and non-grid cells (right) during theta cycles that followed left (red) or right (blue) sweeps detected in the whole-population decoder (1 pair of lines per animal), plotted in head-centered coordinates as in **b**. Note that the decoded position sweeps outwards more uniformly when analysis is restricted to grid cells. Right: Fraction of theta cycles with detected sweeps when decoding position from grid cells or non-grid cells. Dots connected by lines show data from the same session in the same animal (n=16). **e.** Grid modules express sweeps at different scales. Left: Colored blobs show correlation between instantaneous population vectors and reference maps for a single grid module during a single sweep (plotted as in **a**; color corresponding to temporal order of decoding frames). Right: Running trajectory of the rat (grey) during a 3.5s period (same as in **a**), with trajectories of alternating sweeps decoded from single-module population activity (orange/blue for even/odd theta cycles). Note coordinated left-right alternation and the relationship between the grid spacing and sweep trajectory length. **f.** Sweeps of grid cells in different modules are aligned in direction. Heat maps show joint distribution of head-centered sweep directions (left/right) for pairs of simultaneously recorded modules during one recording session (same as **e**). Note strong correspondence between left/right sweeps across the modules. **g.** Sweep length is proportional to grid spacing. Top: Histograms showing distribution of sweep lengths (measured from start-to end-position of each sweep) in the same 3 simultaneously recorded grid-cell modules shown in **e** (module spacing in parenthesis). Bottom: Mean sweep lengths for all modules. Each dot corresponds to one grid module; different animals are plotted with different colors (1 session per animal). **h.** Example sweep decoded from hippocampal ensemble activity (all cells) during foraging in an open field arena (plotted as in **a** and **e**). **i.** Hippocampal sweeps follow MEC-parasubiculum sweeps. Top: Each panel shows decoded sweeps from co-recorded hippocampal cells (green) and MEC-parasubiculum cells (blue) over 5 successive theta cycles. The rat’s current location and its future path are indicated by arrowhead and grey line. Note alignment of hippocampal and MEC-parasubiculum sweeps. Bottom: Plot shows the forward progression of decoded sweeps as a function of time from the beginning of each theta cycle during an example recording session. The offset between decoded position and lowpass-filtered decoded trajectory was projected onto the rat’s head axis and averaged across all theta cycles in the session. Note that hippocampal sweeps are delayed relative to MEC sweeps. **j.** Left: Heat map showing density of sweep directions (head-centered) decoded from co-recorded MEC and hippocampal cells. Note directional alignment of hippocampal and entorhinal sweeps. Right: Fraction of theta cycles where sweeps from both regions pointed to same side of the animal’s head axis for 6 animals with paired hippocampus/MEC-parasubiculum recordings. Red dot and whiskers indicate mean and s.d. **k.** Place-cell sweeps lag behind sweeps in grid cells. Temporal cross-correlation of decoded positions in MEC and hippocampus for 6 animals (1 session per animal). Decoded position was referenced and rotated as in **i** (bottom panel) before cross-correlation. Cross-correlations from individual animals are plotted as individual lines. Note peak correlation values at positive lags, indicating that sweeps in the hippocampus are delayed relative to MEC. Mean and s.d. are indicated by red dot and whiskers.

Sweeps were identified by a sequence detection procedure that looked for near-linear trajectories of decoded positions spanning at least 4 successive time bins of the theta cycle, with analysis restricted to locomotion periods (running speed > 15cm/s). Sweeps were detected in all 16 animals with recording sites in MEC or parasubiculum (Fig. 1b). The number of identified sweeps increased linearly with the number of recorded cells and the number of spikes recorded per theta cycle (Extended Data Fig. 2a). Sweeps were detected in 72.9 ± 3.4% (mean ± s.e.m.) of the theta cycles in rats with more than a thousand cells (3 rats). In the full sample (16 rats, mean of 769 cells), sweeps were detected in 48.0 ± 1.3% of theta cycles. Sweep directions alternated from side to side across theta cycles in all rats (Fig. 1b-c, Extended Data Fig. 2c-d), with left-right-left or right-left-right alternation occurring in 79.8 ± 0.6% of successive sweep triplets (mean ± s.e.m.), significantly more often than when sweep directions were shuffled (61.1 ± 0.2%, >99.9^th^ percentile in all animals). The sweeps were directed forwards at an angle of 23.9 ± 0.7 degrees to either side of the animal’s head direction (mean ± s.e.m. across 16 animals; Fig. 1b; Extended Data Fig. 2c-d), approaching ∼30 degrees to either side in the animals with most cells (Extended Data Fig. 2d). The average length of a sweep was 22.3 ± 0.4 cm (mean ± s.e.m.). Similar left-right alternating sweeps were observed when other decoding methods were used (Extended Data Fig. 3a, Supplementary Video 1).

Having observed sweeps in combined activity of all MEC-parasubiculum cells, analyzed as a whole, we next asked if they were preferentially expressed in spatially modulated neurons such as grid cells (24.8% of the recorded cells, range 8-42% across animals). Restricting the analysis to grid cells revealed a stronger presence of sweeps relative to non-grid cells (although in both cases the absolute numbers detected were smaller than when position was decoded from the full population, with many more cells). Using a criterion of at least 100 cells for the decoding, we identified sweeps in 28.5 ± 2.0% of the theta cycles in grid cells compared to only 13.4 ± 1.4% in a similar number of randomly sampled non-grid cells from the same session (p=0.001, Wilcoxon signed-rank test, 11 rats; Fig. 1d). In two rats with more than 300 grid cells, sweeps were detected in 64.0 ± 9.1% of the cycles in grid cells and 42.7 ± 8.4% in non-grid cells. The contribution of subtypes of grid cells^5,31,32^ was assessed by dropping out either burst-firing grid cells or non-bursting grid cells from the decoder. Burst-firing grid cells were located primarily in MEC layer II and parasubiculum and non-bursting grid cells mainly in MEC layer III (Extended Data Fig. 4b). When burst-firing grid cells were omitted, the number of detected sweeps decreased in comparison to matched control analyses where the same numbers of non-grid cells were excluded at random (Extended Data Fig. 4c). No reduction was observed after dropping non-bursty grid cells. In analyses of individual cells, bursting grid cells were maximally active and displayed out-of-field firing at the end of the theta cycle when the sweep signal was far away from the animal (Extended Data Fig. 4d-e). Firing in non-bursty grid cells, in contrast, was not locked to the time of outgoing sweeps (Extended Data Fig. 4d). In bursty grid cells, the firing fields were sharpened substantially when the activity was plotted as a function of decoded position instead of tracked position (Extended Data Fig. 4f-g). Non-bursty grid cells were not affected by the choice of reference position. Taken together, these comparisons point to burst-firing grid cells as the strongest carriers of the entorhinal sweep signal.

Grid cells are organized in modules, with each module consisting of cells with common grid spacing and field size^16,33^. The fact that grid modules can operate as independent networks^16^, each residing on their own toroidal manifold^5^, raises the possibility that each module has its own sweep dynamics. To determine if sweeps are coordinated across grid modules, we began by sorting the recorded grid cells based on clustering of their spatial rate-map auto-correlograms^5^. We identified 36 modules, with 1-4 modules per animal and 13-215 cells per module. Position was decoded separately for each module with more than 40 grid cells (31 modules in 15 animals). As in the full MEC-parasubiculum population, individual grid modules expressed sweeps that were directed forwards at an angle from the animal’s head direction (19.3 ± 0.9 deg to either side; mean ± s.e.m. across 31 modules), alternating from side to side across theta cycles (alternation in 74.4 ± 0.3% of theta cycle triplets, >99.9^th^ percentile of shuffled values in 28 out of 31 modules; Fig. 1e, Extended Data Fig. 2e, 3b). Sweep directions of co-recorded grid modules were tightly aligned to each other (circular correlation between sweep directions r=0.43 ± 0.009, p<0.001 for all 23 module pairs), pointing to the same side of the head-axis in 70.3 ± 0.4% of theta cycles with detectable sweeps (significantly above a chance level of 50% for all 23 module pairs, p<0.001, binomial test, Fig. 1f). Sweep lengths scaled with the spacing of the grid modules (Fig. 1e; Supplementary Video 2; Pearson correlation between grid spacing and sweep length r=0.95, p=2.9e-16, n=31 modules from 15 animals, Fig 1g). Individual sweeps spanned 19.7 ± 0.09% of the module spacing on average and only rarely exceeded half of the spacing (8.8 ± 0.3% of all sweeps; mean ± s.e.m.; mean sweep lengths from 0.07 to 0.38 m, module spacing from 0.46 to 1.62 m). The proportional relationship between sweep length and grid spacing means that sweeps lengths are approximately equal across modules when mapped onto the modules’ toroidal unit tile.

Because MEC provides a major input to the hippocampus, sweeps may be propagated from grid cells to place cells. To investigate this, we decoded position from place cells in 8 animals with hippocampal implants (6 of which also had MEC-parasubiculum implants). Place cells mirrored grid cells in that they expressed sweeps forward to either side of the animal in an alternating pattern (Fig. 1h-i, Extended Data Fig. 2f). Sweeps were detected in 43.7 ± 2.1% (range: 23.9-69.4%) of identified theta cycles when position was decoded from the joint activity of all hippocampal cells (157-747 cells). Identified sweeps had average lengths of 0.27 ± 0.01 m (mean ± s.e.m.) and offsets of 20.8 ± 1.8 deg relative to the animal’s head direction. The direction of hippocampal sweeps (left or right with reference to the head axis) was aligned to sweeps in simultaneously recorded grid cells in 70.6 ± 2.1% of the theta cycles where sweeps were detected in both brain regions (range: 60.7-84.3%, p<0.001 compared to chance in all animals, one-tailed binomial test). The absolute mean angle between sweep directions in the two cell populations was only 5.5 ± 0.8 deg (correlation: r=0.46 ± 0.036, n=6 rats; Fig 1i-j). Place cell sweeps were delayed compared to simultaneously recorded MEC sweeps (temporal crosscorrelation of position decoded from cells in hippocampus vs. MEC: lag of 19.4 ± 1.6 ms, mean ± s.e.m., 6 rats, all cells in each region; Fig 1k, Extended Data Fig. 2g), raising the possibility that hippocampal place cell sweeps are inherited from entorhinal grid cells^24,34^.

### Sweeps are aligned with an alternating direction signal in a separate cell population

The shared direction of sweeps across grid modules implies an overarching coordination mechanism, such as a common directional input signal^11,13^. To search for a direction signal that matches the direction of sweeps, we examined cells in MEC-parasubiculum with time-averaged tuning to the animal’s allocentric head direction, generally referred to as head direction cells^35–37^. Among all cells recorded in the region, 29.5% (3,632/12,300 cells) displayed significant and stable head direction tuning (16 animals, 1 session per animal; Fig. 2a and Extended Data Fig. 5a). As in previous studies^36,37^, these cells were often more broadly tuned than the “classical” HD cells recorded in anterior thalamus^38^ or presubiculum^35^, with directional firing fields of the former spanning angles as wide as 120 degrees (tuning width: 111.7 ± 23.9 deg, mean ± s.d.; Fig. 2a). Most of the direction-tuned cells in our sample, including conjunctive grid×direction cells (referred to as ‘conjunctive grid cells’ hereafter), were strongly modulated by the local theta rhythm (Fig. 2a and Extended Data Fig. 5a). These cells were anatomically segregated from non-conjunctive (‘pure’) grid cells, with most of the theta-rhythmic directional cells located in parasubiculum (85.6% or 1,699/1,984, 14 rats; Fig. 2b, Extended Data Fig. 1a-b).

**Figure 2.**
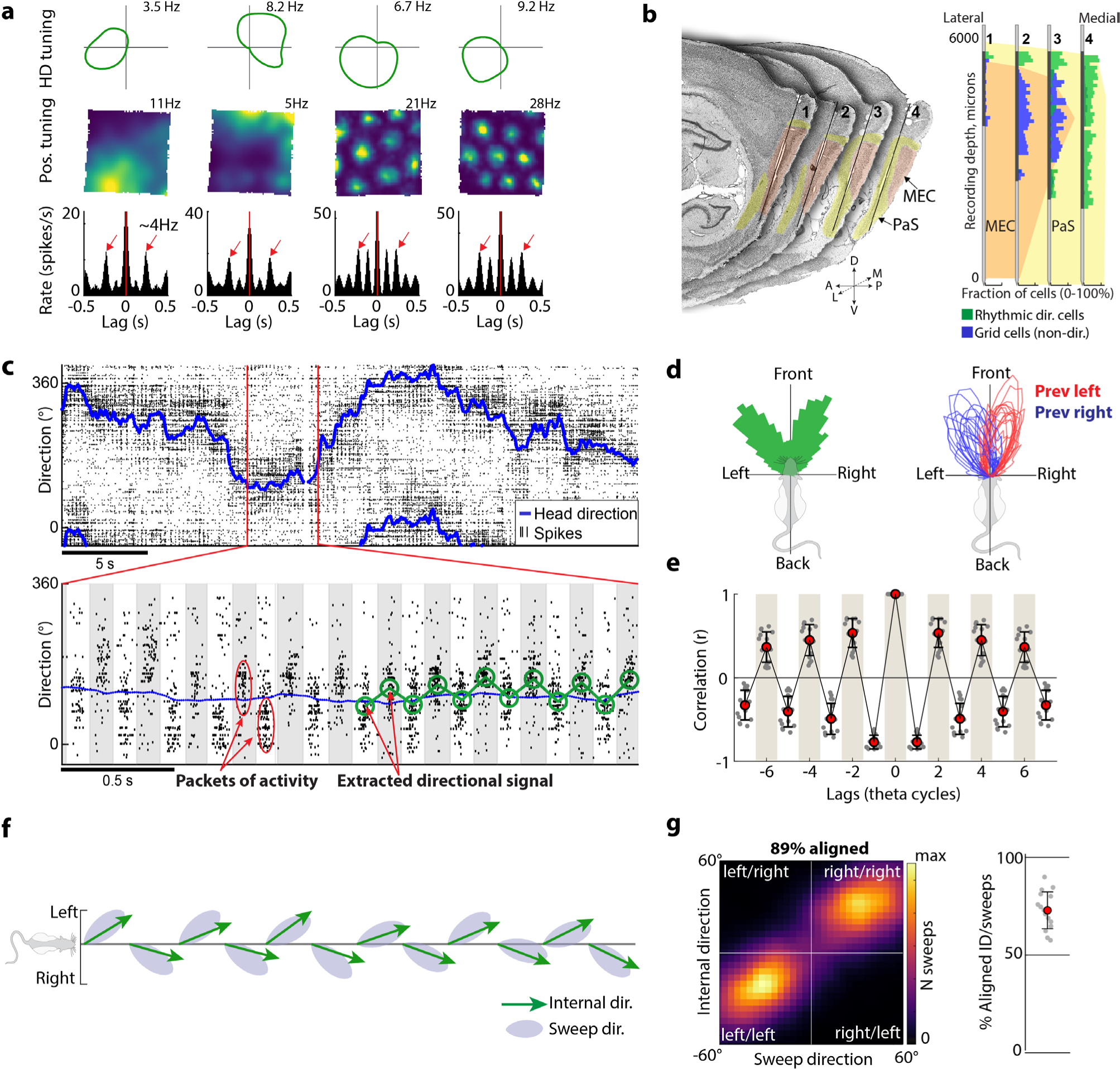
Sweeps are aligned with an alternating direction signal in parasubiculum. **a.** Many theta-modulated cells in parasubiculum are broadly tuned to the animal’s head direction. Top row: circular firing rate plots showing tuning to head direction (HD) for four example cells. Middle: 2D position firing rate maps for the same cells. Bottom row: temporal auto-correlograms of the cells’ firing rates. Arrows indicate temporal lags corresponding to the second theta peak (∼250 ms). Note prominent theta rhythmicity, varying degrees of theta cycle skipping (stronger in the first two cells), and conjunctive grid tuning in some cells. **b.** Theta-rhythmic directional cells are localized primarily in parasubiculum. **Left:** Serial sagittal sections from rat 25953 showing tracks from a 4-shank Neuropixels 2.0 probe. Sections are organized from lateral to medial and each section contains the track from one of four recording shanks (∼250 µm apart). Parasubiculum and MEC are highlighted in yellow and orange, respectively. **Right:** Density of theta-rhythmic ‘internal direction’ cells (green) and pure grid cells (blue) along each of the probe shanks (quantified as proportion of all cells at corresponding depths). Anatomical location of recording sites is indicated by background color. Black portions of probe shanks show active recording sites (combining 7 sessions with different site configurations). **c.** Raster plot showing flickering of direction-tuned activity around the animal’s head direction on successive theta cycles. **Top:** Solid line shows head direction of a rat (rat 25843, session 1) during 30 s of running in an open arena. Black rasters indicate spikes fired by 533 co-recorded internal direction cells, mostly from parasubiculum. Cells (rows) are sorted according to preferred firing direction. **Bottom:** Directional activity in parasubicular direction cells is dominated by theta-timescale population dynamics. Population activity is shown for a 4 second extract from top panel. Theta cycles are indicated by alternating grey and white background shading. Discrete, theta-paced “packets” of population activity alternate between the left and right of the animal’s head direction (solid blue line). Green circles (in right half of plot) show the decoded direction in each theta cycle. **d.** The parasubicular direction signal is correlated with, but distinct from, tracked head direction (circular correlation r=0.79 ± 0.006; absolute mean offset 4.7 ± 0.2 deg; 16 animals). **Left:** Plotting the offset between head direction and decoded direction in a polar histogram shows a systematic relationship between the two variables (rat 25953, session 4). Decoded direction is distributed at two principal angles on either side of head direction. **Right:** Distributions of decoded direction (relative to head direction) for all 16 animals. Each red or blue line shows the distribution of decoded directions when the previous decoded direction was oriented to the left (red) or right (blue). **e.** Decoded direction systematically alternates between left and right on successive theta cycles. Shown are temporal autocorrelograms of angles between decoded direction in successive theta cycles, for all animals. Grey dots correspond to individual animals. Mean and s.d. are indicated by red dots with whiskers. Theta cycles are indicated by grey and white background shading. Note peaks at every other theta cycle, indicating rhythmic alternation of decoded direction from side to side. **f.** Sweeps are aligned to the direction signal. **Top:** Decoded direction (green arrows) and sweep direction (grey patches) over 12 successive theta cycles (∼1.5 seconds, same session as c) plotted with reference to the animal’s head direction (horizontal line). Theta cycles are evenly spaced along the horizontal axis. Note that internal direction and sweeps are aligned to each other. **g.** Left: Heat map showing joint distribution of decoded direction and sweep direction (both in head-centered coordinates) across theta cycles throughout the recording session shown in **c** and **f**. Right: Alignment of decoded direction and sweeps across animals was quantified by computing the fraction of theta cycles where they pointed to same side of the animal’s head axis. Each dot corresponds to one animal. Means (± s.d.) are indicated with colored dots (whiskers).

At the population level, there was strong correspondence between sweep direction in grid cells and the direction encoded by theta-modulated directional cells. While the population activity of the directional cells roughly tracked the animal’s head direction (Fig. 2c, top panel), sub-second analyses revealed discrete, theta-paced packets of coordinated activity that flickered from left to right on successive theta cycles (Fig 2c, bottom panel). To quantify these switches, we turned to the same population-vector correlation methodology that we previously used to decode position. Direction was decoded by correlating firing rate population vectors at the center of each theta cycle with the session-averaged population vectors for each head direction. The decoded signal, referred to as ‘internal direction’, alternated from side to side of the head axis in 86.1 ± 0.5% of theta cycle triplets (>99.9^th^ percentile of shuffled values in all 16 animals; Fig 2c-e, Extended Data Fig. 3c-d), with peak offsets at 19.9 ± 0.6 deg on either side of the head axis (27.9 ± 0.9 deg in the three animals with most cells). Sweeps in grid cells were aligned with the decoded internal direction (circular correlation between sweep direction and internal direction: r=0.66 ± 0.01, p<0.01 in all animals; absolute mean angle 4.4 ± 0.2 deg, mean ± s.e.m.; Fig 2f-g, Supplementary Video 3). The two signals pointed to the same side of the head axis in 72.5 ± 0.6% of theta cycles (p<0.01 compared to chance in all animals, one-tailed binomial test). The rigid left-right alternation of the population direction signal provides an explanation for the phenomenon of theta cycle skipping in individual cells, which was present in most directional cells and grid cells (Fig. 2a; Extended Data Fig. 2b and 5b). Theta cycle skipping is a pattern of activity where direction or position-tuned cells predominantly fire on every other theta cycle, resulting in prominent peaks in the autocorrelogram at even multiples of the theta period (∼250 and 500 ms)^10,39–41^. The presence or absence of firing on alternating cycles likely reflects the left-right dynamics of grid and direction cells.

To determine whether cells that participate in the alternating direction signal are distinct from classical head direction cells, we compared each cell’s relative tuning strength to tracked head direction versus decoded (‘internal’) direction (Extended Data Fig. 5c). Theta-rhythmic directional cells, including conjunctive grid cells, were more strongly tuned to internal direction (‘internal direction cells’), while the smaller sample of non-rhythmic directional cells faithfully followed tracked head direction (‘head direction cells’; Extended Data Fig. 5c). The latter cells were generally located outside the area with internal direction cells, in deep layers of MEC, presubiculum or postrhinal cortex (Extended Data Fig. 5d-e). The coupling between sweeps and direction signals persisted when conjunctive grid cells were excluded, and position and direction were decoded separately from non-conjunctive grid cells and ‘internal direction’ cells (Extended Data Fig. 5f).

### A microcircuit for directing sweeps

The correlations between internal direction and sweep signals, along with the strong projections from parasubiculum to layer II of MEC^42^, point to internal direction cells as a possible determinant of sweep direction in grid cells. To examine if this is reflected in the functional connectivity of the circuit, we temporally cross-correlated the spikes of all recorded cell pairs (1,593,297 cell pairs, 16 animals; Fig. 3a). Pairs where one cell consistently fired ahead of the other at short latency were defined as putatively connected^43,44^. Putative connections were found within and between functional cell classes (Fig. 3b; Extended Data Fig. 6a-d). They included connections from internal direction cells to conjunctive grid cells (0.09±0.007% of cell pairs) and from conjunctive grid cells to pure grid cells (0.33±0.024%). Internal direction cells and conjunctive grid cells had more frequent functional connections to bursty pure grid cells (in MEC layer II, Extended Data Fig. 4b) than to non-bursty pure grid cells (in MEC layer III; Extended Data Fig. 4b) (0.26±0.014% vs. 0.006±0.003%, internal direction and conjunctive grid cells combined; p=5.8*10^-65^, Fisher’s exact test; Extended Data Fig. 6a).

**Figure 3.**
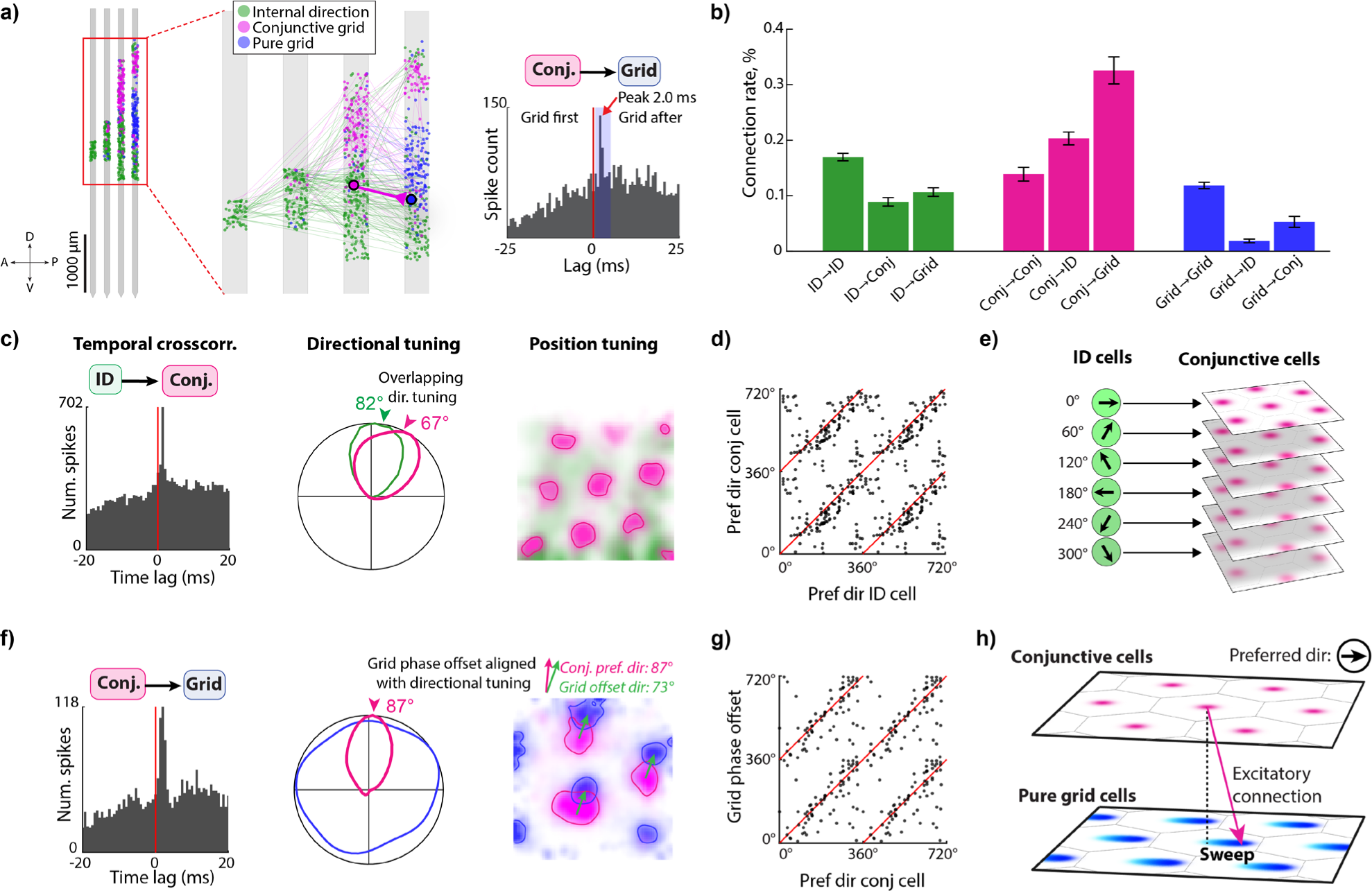
A microcircuit for directing sweeps. **a.** Identification of putative monosynaptic connections. Left: Dots show locations of co-recorded internal direction cells (green), conjunctive grid cells (pink) and pure grid cells (blue) along each of the four recording shanks during an example session. Lines show detected connections between pairs of cells with color indicating the functional identity of the presynaptic cell. One example pair of connected cells (conjunctive grid to pure grid) is highlighted. Right: Cross-correlogram between the firing rates of the two cells highlighted in the left panel. Blue rectangle indicates the time window (0.7-4.7ms) used to detect putative monosynaptic connections. A clear, short-latency positive peak (red arrow) in the cross-correlogram is consistent with the possibility of a monosynaptic excitatory connection from cell 1 to cell 2. **b.** Estimated connection probability (in %, means and standard deviations, data pooled across 16 animals) for projections originating from each of three functional classes (internal direction (‘ID’), conjunctive grid (‘conj’), and pure grid (‘grid’)). Note connections from internal direction cells to conjunctive grid cells and from conjunctive grid cells to pure grid cells, in addition to recurrent connections within each class. **c.** A cell pair consisting of an internal directional cell (green) with a putative projection to a conjunctive grid cell (pink). Left to right: Temporal cross-correlogram of firing rates, directional tuning, position tuning (rate expressed by color intensity, ranging from 10^th^ to 99^th^ percentile of each cell’s firing rate map) for the putative pre- and postsynaptic cells. Note similar preferred direction of the two cells (middle panel). **d.** Preferred directions for all pairs (dots) of putatively connected internal direction to conjunctive grid cells (150 pairs from 13 animals). **e.** Schematic of inferred connectivity between internal direction cells and conjunctive grid cells. Internal direction cells relay the direction signal to conjunctive grid cells with similar directional tuning (preferred directions indicated by short arrows). **f.** A cell pair consisting of a conjunctive grid cell (pink) projecting to a pure grid cell from the same grid module (blue). Left to right: Temporal cross-correlogram of firing rates, directional tuning and position tuning for putative pre- and postsynaptic cell, plotted as in **c**. The conjunctive grid cell projects to a pure grid cell with position receptive fields that are shifted along the preferred direction of the conjunctive cell. The direction and magnitude of this offset (green arrow, right panel) was found by cross-correlating the spatial rate maps of the cells. **g.** Preferred direction of pre-synaptic conjunctive grid cells and grid phase offset direction between pre- and postsynaptic cell for all pairs (dots) of putatively connected conjunctive grid to pure grid cells (86 pairs from 12 animals). **h.** Schematic of inferred connectivity between conjunctive grid cells and pure grid cells. Conjunctive grid cells project asymmetrically to pure grid cells, and excite pure grid cells with a spatial position phase offset that is aligned to the conjunctive cell’s preferred direction. In this connectivity scheme, activation of a set of conjunctive cells with a particular preferred direction leads to a sweep-like trajectory in that direction in postsynaptic grid cells.

In theoretical models, the grid-cell position signal is translated across the attractor manifold by directional input that causes a phase shift of grid-cell activity in the direction of the input signal, via a layer of conjunctive grid cells^11,13,15,25^. Two observations in the cross-correlation analyses were consistent with this prediction. First, putative connections from internal direction cells to conjunctive grid cells were primarily between cells with similar directional tuning (Fig. 3c-e and Extended Data Fig. 6c, f; angle between tuning directions: 14.1 ± 52.1 deg, mean ± s.d.; ciruclar correlation between tuning directions: r=0.41, p=6*10^-7^, n=150 connected cell pairs from 13 animals; see Extended Data Fig. 6b, e for within-class connections and other combinations). Second, putative connections from conjunctive grid cells to pure grid cells targeted cells with a slightly shifted grid phase (Fig. 3f and Extended Data Fig. 6d; magnitude of grid phase offset: 26.0 ± 12.8% of the grid spacing, mean ± s.d.; n=86 connected cell pairs from 12 animals). The direction of the spatial phase offset closely matched the preferred internal direction of the conjunctive grid cells (Fig. 3f and Extended Data Fig. 6f; correlation between grid phase offset direction and preferred direction: r=0.62, p<5.1*10^-8^; angle between directions: 0.4 ± 55.3 deg, mean ± s.d). This directional and positional alignment of connected cell pairs differed substantially from that of randomly selected non-connected cell pairs (Extended Data Fig. 6g). Taken together, the findings support the notion that sweep direction is determined by activation of internal direction cells, via a layer of conjunctive grid cells (Fig. 3h).

### Sweeps extend to never-visited locations

In place cells, forward-projecting sweeps before the choice point in a maze are thought to reflect a deliberation over behavioral options^9,10^. If sweeps are involved in planning, we would expect their direction to correlate with the rat’s subsequent movement. The sweeps recorded during foraging in the present study showed a high degree of stereotypy not consistent with a role in goal-oriented navigation. However, because most of these sweeps corresponded to navigable and previously travelled paths, they do not rule out an exclusive role for sweeps in trajectory planning. This limitation led us to record in environments where the navigational opportunities were constrained to one-dimensional paths. The rats ran either on an elevated 2.0 m linear track with reward delivered at the ends (5 rats; Fig. 4a, left) or on an elevated “wagon wheel” track consisting of a 1.5 m-diameter circle with two diagonal cross-bridges^5^ (2 rats; Fig. 4a, right). Directional signals were decoded from all MEC-parasubiculum cells as before, by correlating instantaneous population vectors with head-direction tuning curves computed over the whole session, long enough for all directions to be sampled (Extended Data Fig. 7a). In both tasks, the decoded signal pointed consistently to the sides of the tracks, towards places never navigated, in an alternating pattern similar to what we observed in the open arena (Fig. 4a; Extended Data Fig. 7a). As before, alternating direction signals were accompanied by sweeps, decoded from grid cells belonging to a single module, based on the phase relationships of these cells in a prior open field session. Sweeps travelled into the unvisited space along the sides of the tracks in parallel with the direction signals (Fig. 4b).

**Figure 4.**
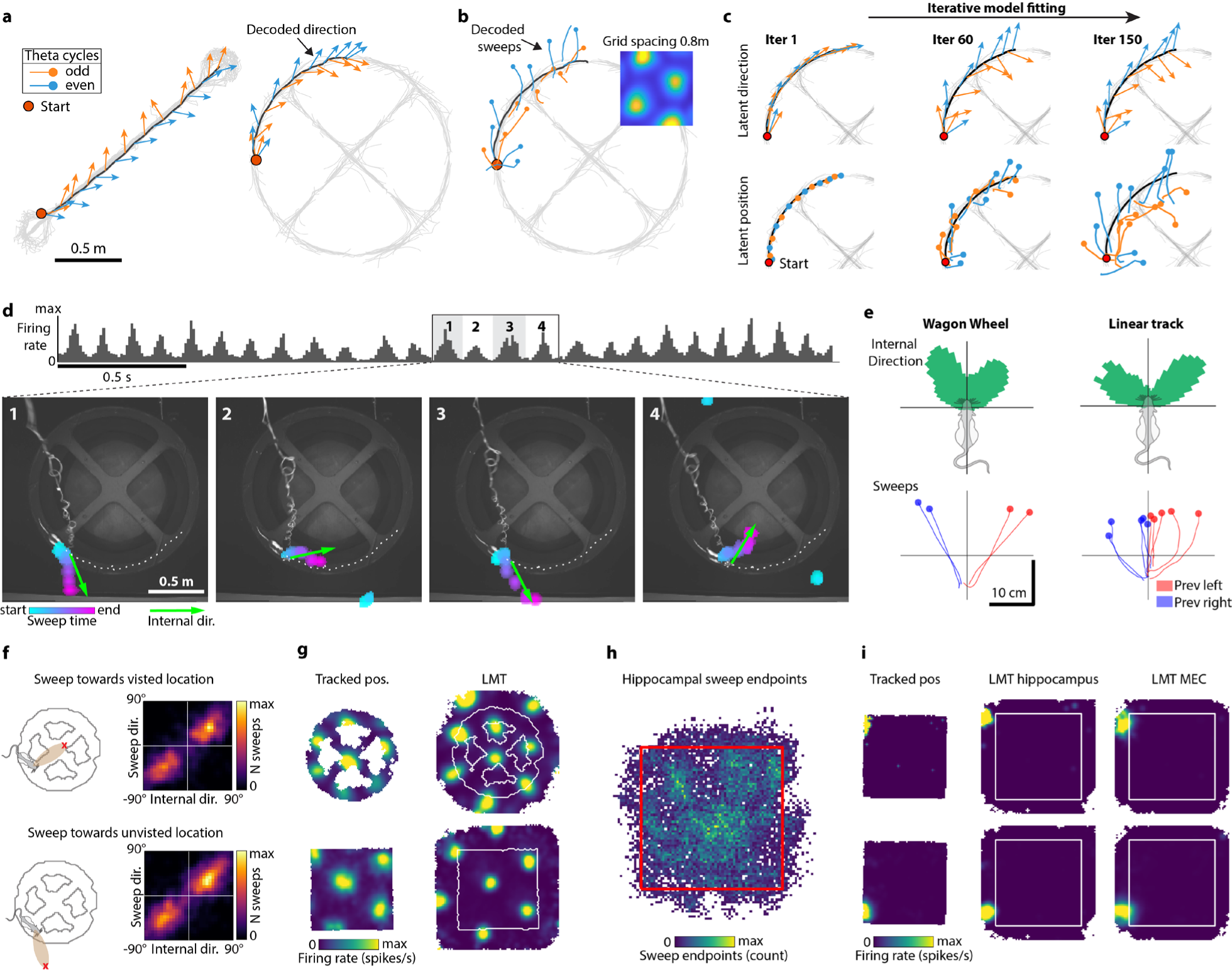
Sweeps extend to never-visited locations. **a.** Decoded (‘internal’) direction signals (population-vector correlation, based on head-direction tuning curves from the same session, all MEC-parasubiculum cells) point towards inaccessible and never-visited locations along an elevated linear track (left) or a wagon-wheel track (right). Panels show decoded direction (arrows of constant length) for successive theta cycles over the course of a 3-s running segment in each task. Alternating theta cycles are plotted with different colors. The running trajectory from the segment is plotted in black, while the full trajectory is plotted in light grey. **b.** Sweeps on the wagon-wheel track decoded from a single grid module, based on rate maps from a preceding open field recording (inset shows example rate map). Decoded position during each theta cycle is plotted on top of the animal’s running trajectory. **c.** A latent manifold tuning (LMT) model allows decoding to include never-visited locations. The LMT model is fitted to the neural data by iteratively updating a latent 1-D and 2-D trajectory. Orange and blue arrows/lines show latent direction (top) and position trajectory (bottom) during 17 theta cycles from a wagon-wheel session at different stages of the model-fitting (left to right). The full running trajectory is plotted in light grey. The latent direction and position signals are initialized with the rat’s actual head direction and running trajectory (iteration 1) but evolve into sweep-like trajectories that cover the 2-D space surrounding the maze (iteration 150). **d.** Sequence of four successive sweeps and internal direction signals during navigation on an elevated wagon-wheel maze (Bayesian decoder, based on fitted LMT tuning curves for all MEC-parasubiculum cells). Each video frame shows the maximum-likelihood direction (green arrow; length is fixed) and position probability (colored bumps, corresponding to positions where p > 99.9^th^ percentile of each frame) during one sweep. Note that sweeps travel into the inaccessible space inside and outside the navigable track. **e.** The stereotypical left-right alternating pattern of internal direction and sweep signals is preserved in 1-D environments. Top row: Circular histogram showing head-centered distribution of fitted LMT internal direction values from one example session on the wagon-wheel track (left) and on the linear track (right). Note that internal direction is bimodally distributed around the animal’s head direction (wagon wheel: 23.7 ± 0.7 deg to either side, left-right alternation in 76.1 ± 5.7% of theta cycles, 2 rats; linear track: 36.4 ± 0.1 deg to either side, left-right alternation in 80.7 ± 0.3% of theta cycles, 5 rats), like in the open field (Fig. 2). Bottom row: Lines show sweeps averaged across each recording session (plotted as in Extended Data Fig. 2e), during theta cycles that followed a left (red) or right (blue) sweep. Each pair of red and blue lines corresponds to one animal (wagon-wheel: 2 rats; linear track: 5 rats). **f.** Out-of-bounds sweeps reliably coincide with internal direction pointing towards the same location. Sweeps terminating inside and outside the environment boundaries are similarly coupled to the internal direction signal. Each plot shows a color-coded 2D histogram of conditional occurrences of the two LMT latent variables – internal direction and sweeps (in head-centered coordinates) – for sweeps that terminate inside (top) or outside (bottom) visited portions of the environment in one animal (rat 25843, same session as **a**-**c**). The circular correlation coefficient between head-centered internal direction and sweep directions was similar when analysis was confined to theta cycles where sweeps terminated inside vs. outside the wagon wheel track: r=0.83 vs. 0.82, 1.2% difference. Similar results were obtained in a second rat with fewer cells (not shown): r=0.34 vs. 0.35, 2.9% difference. **g.** Grid-cell maps extend across never-visited space. Plots show firing-rate maps of the same grid cell in (top row) the wagon-wheel session shown in **a**-**c**, or (bottom row) an open-field session recorded on the same day. Left column based on the original position data; right column based on latent position fitted by the LMT model. The LMT model infers the continuation of grid-like periodic tuning to locations beyond the environmental boundaries. **h.** Hippocampal sweeps extend into never-visited locations. Heatmap of sweep end positions as inferred by the LMT model when fitted to hippocampal data from one example session. Note that many sweeps terminate outside the confines of the open field arena (red box). A total of 17.6 ± 0.6% (mean ± s.e.m.) of hippocampal sweeps terminated outside the open field arena (5 rats). **i.** Place-cell maps include never-visited space. Plots show firing-rate maps for two hippocampal place cells during an open-field session based on (left) original tracked position, (middle) latent position extracted from hippocampal activity, and (right) latent position extracted from co-recorded MEC-parasubiculum cells. Note that the two LMT models agree, both inferring place fields outside the walls of the arena.

Representations of unvisited locations were similarly obtained when a “latent manifold tuning” (LMT) framework^45^ was used to decode direction and position directly from the track data, instead of referring to data from a different session (Fig. 4c, Extended Data Fig. 8). The remaining analysis of unvisited space was therefore performed with LMT analyses. Two latent variables (one in 1D, one in 2D) were respectively initialized with tracked head direction and position, and then iteratively fitted to capture the fast dynamics of internal direction cells and grid cells. After some iterations, the latent position signal started to travel, in alternating directions on successive theta cycles, into the inaccessible open space surrounding the animal’s path, either beside the edges of the elevated tracks (Fig. 4c-f, Extended Data Fig. 7c,e) or beyond the opaque, high walls of the open field (Extended Data Fig. 7d). Sweep directions were strongly correlated with decoded internal direction, regardless of whether the sweeps terminated within or outside the environmental boundaries (Fig. 4f). LMT analysis further showed that individual grid cells were tuned to unvisited locations within a sweep’s length outside the open field box or within the interior holes of the wagon wheel track, in agreement with a continuation of the periodic grid pattern (Fig. 4g, Extended Data Fig. 7g). The same pattern of results was obtained when using a single-cell GLM-based model to infer out-of-bounds tuning for each cell independently (Extended Data Fig. 7h,i).

LMT analyses showed that sweeps traveled beyond the walls of the open field arena in hippocampal place cells too (Fig. 4h). These sweeps were expressed in coordination with out-of-bounds sweeps in MEC (Fig. 4h, Extended Data Fig. 7j). Individual place cells similarly showed tuning to locations beyond the walls of the arena (Fig. 4i, Extended Data Fig. 7k). Taken together, these findings show that a seamless map of ambient space – including grid cells, internal direction cells and place cells – is unfolded independently of whether the animal ever visits the locations covered by the sweep signals.

### Sweeps and internal direction signals persist during sleep

If sweeps are a fundamental feature of the grid cell system generated entirely by local circuit properties, they might be present regardless of sensory input, past experience, or behavioral state. In agreement with this idea, sweeps maintained their stereotypic alternation profile during foraging in darkness and in novel environments (Extended Data Fig. 7f), as well as during sleep in a resting chamber (Fig. 5). Data from the sleep sessions were analyzed further. Considering that head direction cells and grid cells traverse the same low-dimensional manifolds during sleep as in the awake state^5,18,19,46,47^, we decoded position and direction during sleep using fitted LMT tuning curves from the preceding or succeeding wake session in the open field. Because the persistence of phase relationships across environments and brain states is stronger within than between grid modules^5,18,19^, we restricted our analysis of sweeps during sleep to individual grid modules.

**Figure 5.**
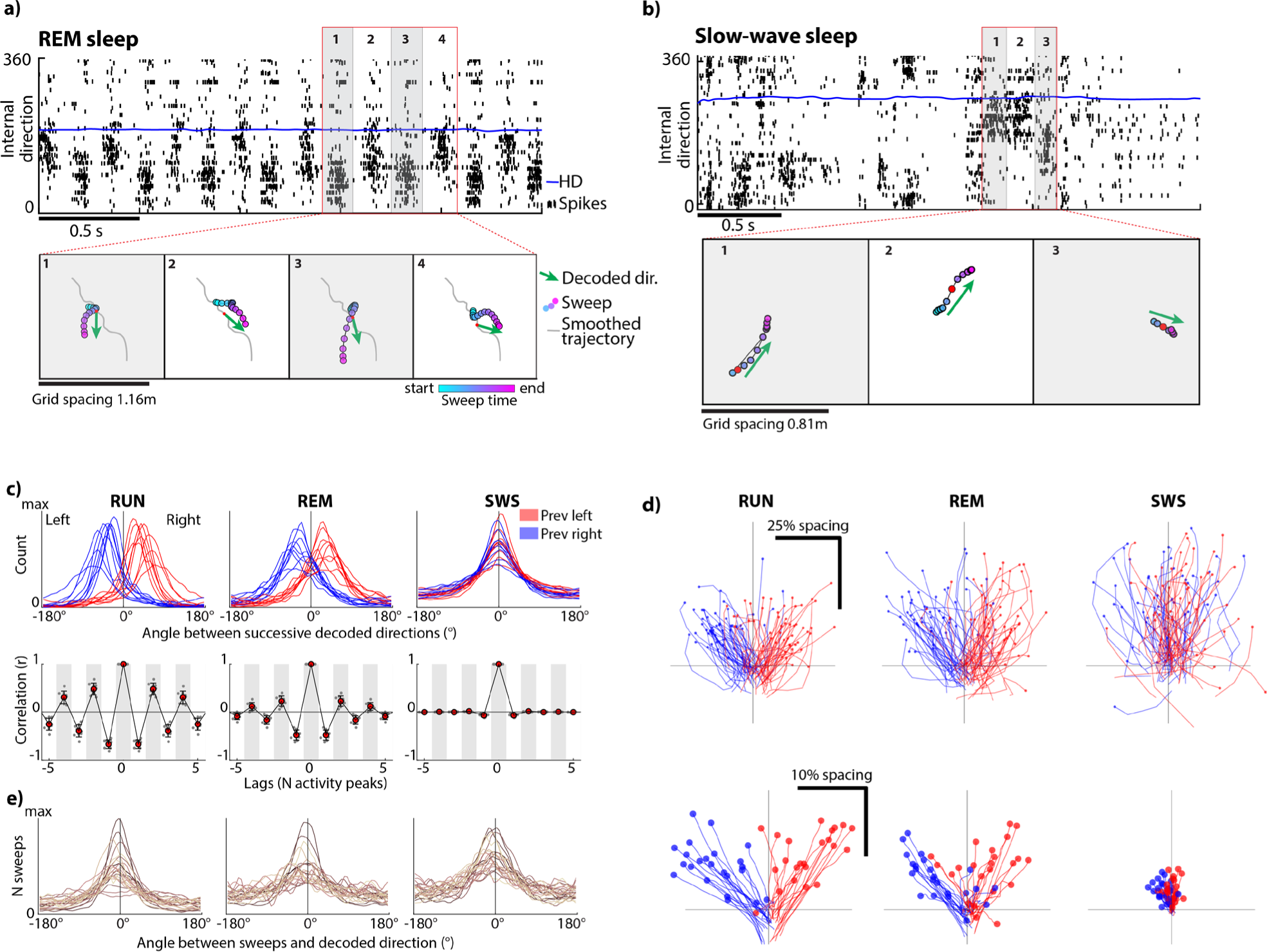
Sweeps and internal direction signals persist during sleep. **a.** Sweeps and alternating internal direction signals are preserved during REM sleep. Top: Raster plot with spike times (black ticks) of internal direction cells sorted by preferred firing direction, and tracked head direction (solid blue line), during a 2.5 s extract from an epoch of REM sleep. Note theta-paced, left-right alternating packets of direction-correlated activity. Bottom: Decoded sweeps (filled circles, color-coded by time) from grid cells of one module and decoded internal direction from direction-tuned cells (green arrows) during 4 successive theta cycles (labelled 1-4 in the top panel). Grey line shows the reconstructed trajectory after smoothing with a wide gaussian kernel (σ = 100 ms) over the entire 2.5 s period. For both position and internal direction, a Bayesian decoder was used, using tuning curves estimated by the LMT model during an open field foraging session from the same recording day. **b.** Internal direction-aligned, non-rhythmic trajectories during SWS. Top: Raster plot (as in **a**) showing activity of internal direction cells during a 3 s extract from SWS. Note sharp transitions between up and down states (with and without spiking activity, respectively). Bottom: Decoded position from a single grid module (as in **a**) during each of three highlighted segments in the top panel. Note sweep-like position trajectories in the same direction as the decoded internal direction (green arrows) in each segment of the up-state. **c.** Decoded direction alternates from side to side during wake and REM, but not during SWS. Top: Distribution of angles between decoded direction (same decoding method as **a**) at successive peaks of activity, when the previous decoded direction was directed to the right (red) or left (blue). Bottom: Autocorrelogram of decoded direction across brain states. Note rhythmic alternation during wake and REM. **d.** Top: Decoded trajectories for 100 example sweeps from one example grid module (same decoding method as **a**), sorted based on whether the previous decoded direction pointed to the right (blue) or left (red). Since spatial representations are decoupled from physical movement during sleep, sweep trajectories are referenced to the low-pass-filtered decoded trajectory (smoothed with a 100 ms gaussian kernel) and aligned to a “virtual head direction” (low-pass-filtered decoded direction). Individual sweeps are plotted as separate lines. Note that during wake and REM most sweeps are directed forward to the opposite side of the previous sweep (few red sweeps on left side, few blue sweeps on right side). Bottom: Averaged sweeps across brain states for all 23 grid modules of all 9 animals. Sweeps were referenced, rotated and sorted as in the top panel, normalized by the grid module’s spacing, and then averaged across all theta cycles. Each pair of red and blue lines corresponds to one module. **e.** Sweeps are aligned with direction signals in all brain states. Panels show distribution of angles between decoded (same method as **a**) direction and position (sweep) signals during wake, REM sleep and SWS (left to right). Note alignment in all states. Individual modules are plotted as separate lines.

Sleep sessions were segmented into epochs of rapid eye movement (REM) sleep and slow wave sleep (SWS), based on electrophysiological and behavioral criteria (Extended Data Fig. 9a). We recorded on average 99.1 ± 34.6 min (mean ± s.d.) of sleep per rat in 9 rats, out of which 11.3 ± 4.5% was classified as REM sleep and 88.7 ± 4.5% as SWS (average epoch durations: 85 ± 54 s and 232 ± 223 s; mean ± s.d.). During REM sleep, the population dynamics of internal direction cells and grid cells was similar to wake. Spiking activity was highly theta-rhythmic (Fig. 5a, Extended Data Fig. 9b-c), and internal direction decoded from neighboring peaks of activity alternated from side to side (alternation in 70.1 ± 0.6% of triplets of neighboring peaks, mean alternation after shuffling peaks: 50.0%; angles between decoded direction at neighboring peaks: 30.5 ± 0.9 deg to the opposite side of the previous pair of peaks, mean ± s.e.m., mean shuffle: 0.04 deg, 9 rats; Fig. 5a,c). Individual grid modules expressed rhythmic sweeps that reset at each theta cycle, alternating in direction across successive theta cycles (Fig. 5d-e). Sweep directions were aligned with decoded internal direction (absolute mean angle between sweeps and internal direction: 18.4 ± 2.0 deg, offset in shuffled data: 91.6 ± 1.1 deg; Fig. 5e). Sweep lengths were comparable to those recorded in the open field (25.4 ± 0.13% of grid module spacing, mean ± s.e.m.). The sweeps were nested on top of behavioral time-scale spatial trajectories extending several meters over the course of tens of seconds (Extended Data Fig. 9g), mirroring how sweeps extend outwards from a slower running trajectory during awake exploration (Fig. 1a).

During SWS, the population dynamics was less regular, and spiking activity was confined to brief bursts during up-states, followed by silent down-states (Fig. 5b, Extended Data Fig. 9b-c). There was no periodic left-right alternation of the internal direction signal between successive local maxima in the summed population activity, neither within nor across up-states (alternation on only 52.4 ± 0.1% of triplets of neighboring peaks, mean shuffle: 50.0%; average angle between neighboring activity peaks: only 5.9 ± 0.3 deg, mean ± s.e.m., mean shuffle: 0.01 deg; Fig. 5b-c). During SWS, bursts of direction-tuned population activity were often accompanied, in grid cells, by sweep-like trajectories that were aligned to the decoded internal direction signal (sweeps were observed in 30.0 ± 0.40% of identified peaks in internal-direction cell activity, mean ± s.e.m.; absolute mean angle between sweep-like trajectories and internal direction: 13.7 ± 1.5 deg, corresponding offset in shuffled data: 86.7 ± 1.7 deg; Fig 5b,d-e, Extended Data Fig. 9d-f). Like the internal direction signals, the sweep trajectories were not rhythmic (Extended Data Fig. 9h). Thus, coordinated direction and sweep signals can exist in all states but theta activity is required to maintain the rigidly coupled rhythmic side-to-side pattern.

### Sweeps sample nearby space with optimal efficiency

Having dissociated sweep and direction signals from navigational goals and spatial decision processes, we hypothesized that the alternation of sweep directions instead signifies a strategy for efficiently sampling ambient space. To test this hypothesis, we simulated an ideal sweep-generating agent that chooses sweep directions that tile space with optimal efficiency. We created a simple model of a sweep’s spatial footprint (Extended Data Fig. 10a) based on our prior observation that sweep lengths are proportional to the module’s grid spacing (Fig. 1g). At each time step, the sweep-generating agent was tasked with choosing a sweep direction that minimizes overlap with the area covered cumulatively by previous sweeps, without foresight of upcoming sweep directions. When the agent was moved along on a linear path at a constant speed, it generated sweeps that alternated between two characteristic directions, 33.0 ± 0.008 deg to either side of the movement direction (mean ± s.e.m. across 1000 runs with random initial conditions; Fig. 6a, Extended Data Fig. 10b), resembling empirical sweep directions on a linear track (Fig. 4a). The prevalence of alternation was quantified using a score that measured alternation in a sliding window of three successive sweep directions, with scores ranging from 0 (no alternation) to 1 (perfect alternation). The agent reliably converged on an alternating pattern that exceeded chance level from the third sweep (mean ± s.d. alternation score at third sweep: 0.66 ± 0.34, p=2.1e-56 with respect to chance level of 0.40, one-tailed sign test) and approached perfect alternation towards the end of the run (alternation score: 0.97 ± 0.020). The simulation results point to alternation as a stable regime for minimizing overlap of successive sweeps (Fig. 6b, Extended Data Fig. 10b). Robust alternation was obtained across a range of sweep widths (Extended Data Fig. 10c).

**Figure 6:**
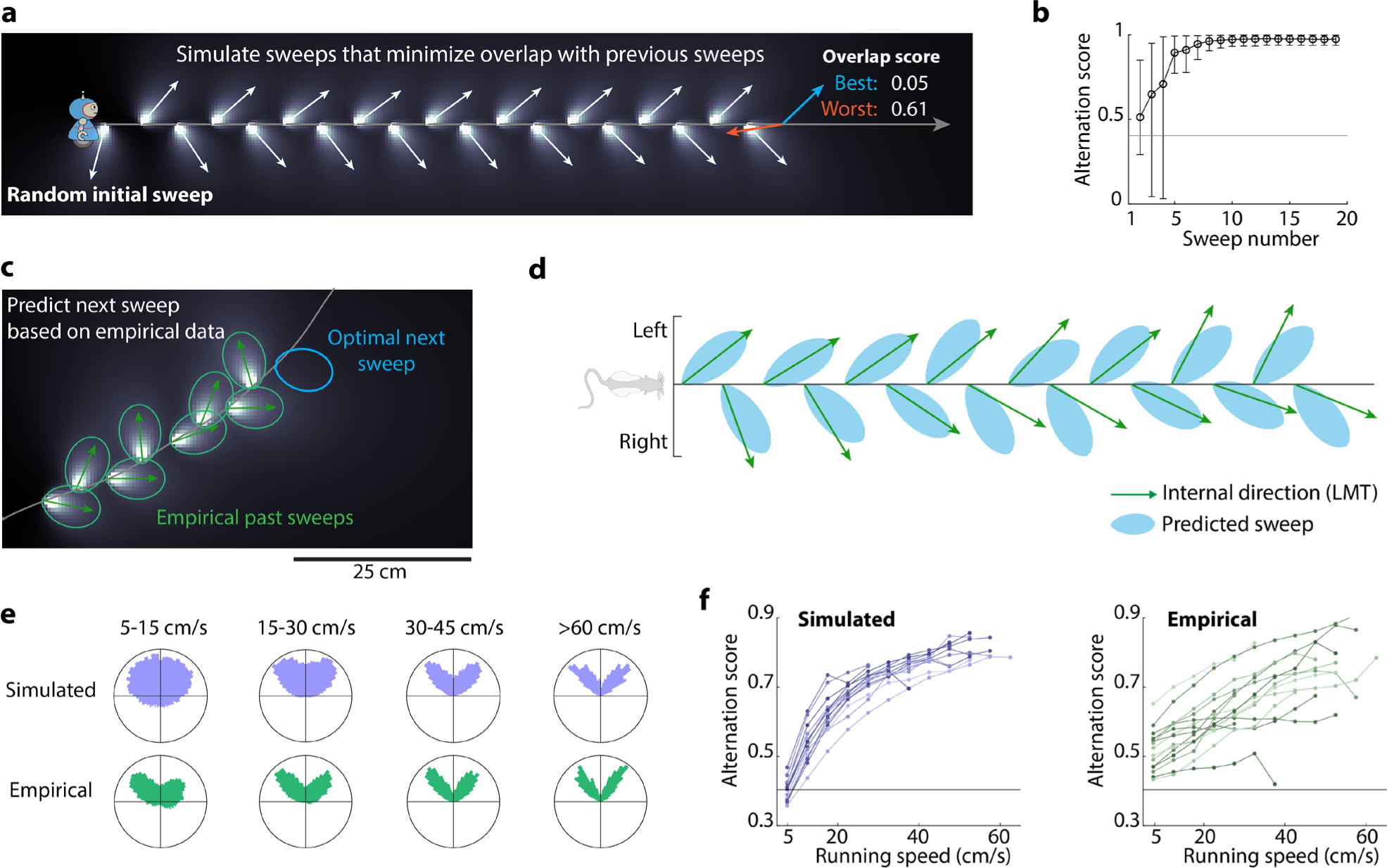
Sweeps sample nearby space with optimal efficiency. **a.** Simulation of a sweep-generating agent deploying an algorithm that minimizes each sweep’s overlap with preceding sweeps. The agent is moved along a scale-free linear path and generates a sweep at each time step by placing a beam-like sweep footprint in a specified direction. The selected sweep directions spontaneously alternate between two directions relative to the direction of movement. **b.** Alternation of sweep directions in 1,000 runs of the simulation in **a**. Circles show mean alternation score (range 0 to 1) for each time step and error bars indicate the 5^th^ and 95^th^ percentiles across runs. Horizontal line indicates expected alternation score for random, uniformly distributed angles. Note reliable and near-perfect alternation within few steps of the simulation. **c.** The agent accurately predicts empirical sweep directions. Unlike the “free” simulations in **a**-**b**, this “hybrid” version of the model predicts the direction of a sweep in a given theta cycle (blue), given the recent history of decoded direction values (green). **d.** Internal direction fitted by the LMT model (arrows) and sweep directions predicted by the agent (colored patches) during 16 theta cycles in an open field session (both plotted in a head-centered reference frame). Theta cycles are evenly spaced along the horizontal axis. Note alignment between decoded and predicted directions. **e.** Running speed modulates the distribution of sweep direction. Top row: distribution of head-centered sweep directions chosen by the agent in an open-field session (same as in **d,** but simulation now run in “free” mode without influence of decoded direction), shown separately for different running-speed bins. Bottom row: distributions of head-centered internal direction decoded from empirical data (same session). Note that in both cases, the distribution’s bimodality increases with running speed. **f.** Alternation of sweep directions increases with speed. Left: Average alternation score of the sweep directions chosen by the agent in “free” mode during an open field session, binned at different running speeds. Each set of connected dots shows data from a different animal. Horizontal line indicates expected alternation score for random, uniformly distributed angles. Right: same, but for decoded (“empirical”) internal direction from the same 12 animals.

To determine whether the agent could predict the angles of individual sweeps in the empirical neural data, we moved the agent along the animal’s recorded locomotor path in the open field arena, and for each sweep detected in the empirical data, we determined the optimal sweep direction at the animal’s position based on previous empirical directions (using the fitted LMT internal direction variable; Fig. 6c). In order to prevent the agent from being influenced by sweeps that occurred at similar locations far in the past, we introduced a temporal decay to the cumulative coverage trace of previous sweeps (Extended Data Fig. 10d). The agent’s chosen directions aligned with empirical directions in all 12 animals with sufficient numbers of internal direction cells to reliably extract internal direction (correlation simulated vs. decoded direction: r=0.48±0.014, p<0.001 in all animals; mean ± s.e.m. offset from the decoded direction: 0.17 ± 0.40 deg; Fig. 6d). Sweep directions of the agent alternated in step with decoded directions in 72.4 ± 0.8% (mean ± s.e.m.) of the theta cycles, more often than expected by chance (p<0.001 for all sessions in all animals with respect to a chance level of 50%, binomial test, Fig. 6d, Extended Data Fig. 10f). This alignment was maintained across a range of temporal decay factors, with the highest accuracy obtained with rapid forgetting (median decay constant τ: 0.01, n=12 animals, corresponding to a halving of sweep trace intensity every 150 ms, Extended Data Fig. 10d). In both the simulated and the empirical data, the egocentric distribution of sweep and direction signals became increasingly bimodal during fast and straight running, with alternation scores increasing with running speed and path straightness (mean ± s.e.m. correlation speed vs. alternation: r=0.92±0.003 for simulated and r=0.96±0.004 for decoded directions; straightness vs. alternation: r=0.917 ± 0.004 and r=0.862±0.013, p<0.05 in 12/12 animals, Fig. 6e-f, Extended Data Fig. 10e).

## Discussion

We show that grid cells encode trajectories that within each theta cycle sweep outwards from the animal’s location, with direction alternating between left and right across successive theta cycles. Sweeps in grid cells were aligned to a similarly alternating directional signal in a separate population of direction-tuned neurons. Sweeps were identified also in hippocampal place cells, but these were delayed compared to sweeps in grid cells, suggesting they were propagated from MEC^24,34^. Collectively, the findings point to a specialized circuit of space-coding neurons that on alternating theta cycles generates paths to locations on the left and the right side of a navigating animal’s trajectory.

The sustained expression of directionally oscillating sweeps and direction signals, and the invariance of the sweep geometry on the grid-cell manifold, point to a fundamental role for sweeps in sub-second mapping of the surrounding environment. Grid cells were shown to sweep with directional offsets that maximize coverage of the ambient space, at lengths traversing a substantial fraction of the periodic grid cell manifold. By way of sweeps, grid cells may link locations in the proximal environment into a continuous two-dimensional map without extensive behavioral sampling^48,49^, which in turn would allow new maps to be formed faster and more effectively than if animals had to physically run each of those trajectories. In familiar environments, alternating sweeps may allow animals to efficiently retrieve representations of the surrounding space, one sector at a time, mirroring the alternating sonar beams of echolocating bats^50^. Our data suggest that such sweep-based encoding and retrieval mechanisms are implemented in the entorhinal-hippocampal circuit.

In our recordings, sweeps rarely reflected the rat’s future trajectory or the location of navigational goals. At first glance, this contrasts with the forward-directed theta sweeps reported previously in hippocampal place-cell ensembles on linear tracks^6–8^ or T-shaped mazes^9,10^. In those studies, sweeps – decoded only with respect to visited positions on the track or maze – travelled down possible future paths, with alternations on the maze stem reflecting upcoming bifurcations. In the present study, we were able to decode representations in the full ambient 2-D space, beyond the animal’s path, either by leveraging the invariance of grid phase relationships from a different condition, or by employing a latent-variable model that characterized position tuning to any location in the nearby environment. The fact that sweeps invariantly oscillated between left and right in the full 2-D space raises the possibility that forward-looking sequences reported previously in linear environments reflect projections of left-right-alternating sweeps onto the animal’s running trajectory, with sweeps towards unvisited lateral locations going undetected since the decoding procedures then used could only match activity with physically visited locations. The findings also provide a framework for understanding corollaries of sweep sequences in individual cells, such as theta phase precession^6,51^, and theta cycle skipping^10,39–41^. However, it should be noted that the presence of a hardwired side-shifting mechanism does not rule out that with training in a structured maze task, sweeps in the hippocampus^9,52^ or downstream^53^ may gradually be directed towards specific goal and reward locations, enabling encoding of preferred routes as neural maps are built up during experience.

Finally, our findings provide some clues to the mechanisms of sweep formation. The functional connectivity analyses raised the possibility that internal direction cells, directly or indirectly through conjunctive grid cells, drive the generation of grid-cell sweeps in the same direction^21–23,25^. The projections from conjunctive grid cells to pure grid cells were asymmetric, in the sense that they preferentially activated cells with grid phases displaced in the direction of the internal direction signal, mirroring a vector integration mechanism proposed for bump movement in continuous attractor network models for grid cells^11,13,15,21^. The scalar component of the vector computation remains to be identified, however. Sweep lengths are not a direct reflection of running speed, since sweeps persist during sleep. Instead or additionally, their length may be influenced by factors such as intensity or duration of the internal direction input. Our observations also leave open the mechanism of directional alternation. Two classes of alternation mechanisms can be envisaged. First, alternation may be hardwired into the connectivity of the circuit. Rhythmic alternations reminiscent of those observed here have been described in many brain systems: in left-right shifting spinal-cord circuits for locomotion^54^, in inspiration-expiration circuits for breathing in the medulla^55^, and in hemisphere-alternating REM sleep circuits of reptiles^56^. Alternations between opposing states in these networks rely on central pattern generator mechanisms in which activity is switched periodically between two internally coupled subcircuits^56–59^. A central pattern generator might underlie also the left-right alternations of the cortical navigation circuit^60^. Second, and alternatively, alternating sweep directions may emerge spontaneously as a result of a spatial overlap-minimizing rule, without explicit implementation of alternation, as shown in the artificial agent simulations. A plausible mechanistic substrate for such a rule exists in single-cell firing-rate adaptation^25,61–63^, which penalizes repeated activation of the same neural activity patterns.

## Acknowledgments

We thank T. Waaga for sharing data and helping out with recording in darkness^64^ (Extended Data Fig. 7f). We thank B.A. Dunn, M. Nau, Y. Burak and M. Witter for discussion and S. Ball, K. Haugen, E. Holmberg, K. Jenssen, E. Kråkvik, and H. Waade for technical assistance. The work was supported by a Synergy Grant to E.I.M. from the European Research Council ERC (‘KILONEURONS’, Grant Agreement N° 951319); FRIPRO grants to E.I.M. (grant number 286225) and M.-B.M. (grant number 300394/H10), two Centre of Excellence grants to M.-B.M. and E.I.M. (Centre of Neural Computation, grant number 223262; Centre for Algorithms in the Cortex, grant number 332640), and a National Infrastructure grant to E.I.M. and M.-B.M. from the Research Council of Norway (NORBRAIN, grant number 295721); as well as a grant from the Kavli Foundation (M.-B.M. and E.I.M.), a grant from the K.G. Jebsen Foundation (grant number SKGJ-MED-022), St. Olav’s Hospital and The Regional Health Authorities of Mid Norway (2020/7569), a gift from Erik Solberg and family, a direct contribution to M.-B.M. and E.I.M. from the Ministry of Education and Research of Norway, and a Ph.D. fellowship grant from NTNU’s Faculty of Medicine and Health Sciences to A.Z.V. Clipart images were adapted with permission from https://scidraw.io (rat image) and https://www.nicepng.com (robot image).

## Author Contributions

All authors planned and designed experiments, conceptualized and planned analyses, and interpreted data; A.Z.V. and R.J.G. performed experiments; A.Z.V. and R.J.G. visualized and analyzed data; A.Z.V., R.J.G. and E.I.M. wrote the paper. All authors discussed and interpreted data. M.-B.M. and E.I.M. supervised and funded the project.

## Supplementary Information

is available for this paper.

## Reprints and Permissions

information is available at www.nature.com/reprints

## Methods

### Subjects

The data were obtained from 18 Long Evans rats (17 males, 1 female; 300–500 g at time of implantation). Data from five of the animals have been used for other purposes in published data^5,64^. The rats were group-housed with three to eight of their littermates before surgery and were thereafter housed singly under enriched circumstances in large two-story metal cages (95 × 63 × 61 cm) or smaller Plexiglas cages (45 × 44 × 30 cm). They were kept on a 12-h light–12-h dark schedule in humidity and temperature-controlled rooms. Experiments were approved by the Norwegian Food Safety Authority (FOTS ID 18011). and performed in accordance with the Norwegian Animal Welfare Act and the European Convention for the Protection of Vertebrate Animals used for Experimental and Other Scientific Purposes.

### Surgery and electrode implantation

Eighteen rats were implanted with Neuropixels silicon probes targeting either the MEC-parasubiculum region (10 rats, of which 3 rats were implanted bilaterally), the hippocampus (2 rats), or both regions (6 rats). Neuropixels prototype phase 3A single-shank probes^26^ were used in 8 of the rats, and prototype 2.0 multi-shank probes^27^ were used in the other 10 rats. Probes targeting MEC-parasubiculum were implanted 4.2-4.7 mm lateral to the midline and 0.0-0.3 mm anterior to the transverse sinus, at an angle of 18-25 deg in the sagittal plane, with the tip of the probe pointing in the anterior direction. Probes were lowered to a depth of 4,100-7,200μm. Hippocampal probes were positioned vertically ML 1.4-3.0 mm from the midline and AP 1.9-4.0 mm posterior bregma. In rats with probes in both MEC-parasubiculum and hippocampus, the two probes were implanted in different hemispheres. The implants were secured with dental cement. A jeweller’s screw in the skull above the cerebellum was connected to the probe ground and external reference pads with an insulated silver wire. The detailed procedure for chronic Neuropixels surgeries has been described elsewhere^27^. Postoperative analgesia (meloxicam and buprenorphine) was administered during the surgical recovery period. Rats were left to recover until they resumed normal foraging behavior, at least 3 hours after surgery.

### Electrophysiological recordings

Instruments and procedures were similar to those described for Neuropixels recordings used in the lab^5,26,27,64^. Briefly, neural signals were amplified (gains of 500 for phase 3A and 80 for 2.0 probes), filtered (0.3-10 kHz for phase 3A and 0.005-10 kHz 2.0 probes) and digitized at 30 kHz by the probe’s on-board circuitry. Signals were multiplexed and transmitted to the recording system along a tether cable. SpikeGLX software (https://billkarsh.github.io/SpikeGLX/) was used to control acquisition and configure the probes. A motion capture system – based on retroreflective markers on the implant, OptiTrack Flex 13 cameras, and Motive recording software – was used to track head position and orientation in 3D. The 3D tracking coordinates were subsequently projected onto the horizontal plane for estimation of 2D position and head direction azimuth. An additional camera (Basler acA2040-90umNIR) was used to capture overhead infra-red video in a subset of the recordings. Overhead video frames were aligned to OptiTrack tracking data with an affine transformation between corresponding points in the video and tracking data. Timestamps from each data stream were synchronized as previously described^5,64^, by generating randomized sequences of digital pulses with an Arduino microcontroller and sending them to the Neuropixels acquisition system as direct TTL input and to the OptiTrack system and video camera via infrared LEDs placed on the edge of the arena.

### Behavioral procedures

Recordings were obtained while animals foraged in an open field, while they navigated for rewards on a linear track or on a wagon wheel-shaped circular track, or during sleep. Due to the previously reported gradual decay in signal quality over the first 7-14 days after probe implantation^65^, most recordings were performed within the first week after surgery (full range: 0-151 days post-operatively). All behavioral tasks for a given animal were performed in the same recording room (except for one recording in a novel room, Extended Data Fig. 7f), often consecutively on the same day. Recording sessions were sometimes interrupted to remove twists from the Neuropixels tether cable. During pre-surgical training, some of the rats were food-restricted, maintaining their weight at a minimum of 90% of their free-feeding body weight. Food restriction was not used in any of the animals at the time of recording.

### Open-field foraging task

18 rats foraged for randomly scattered food crumbs (corn puffs or vanilla cream cookies) in a square open-field (OF) box with a floor size of 150 × 150 cm and a height of 50 cm. The floor was made of black rubber; walls were made of black expanded PVC plastic. The arena was placed on the floor centrally in a large room (16 or 21 m^2^) with full visual access to background cues. A large white cue card was affixed to one of the walls (height same as the wall; width 41 cm; horizontal placement at the middle of the wall). In all illuminated trials, at the time of the surgery, each rat was already highly familiar with the environment and the task (having experienced 10–20 training sessions prior to surgery, lasting at least 20 min each). Recording sessions lasted 23-141 min.

In one exceptional case, a rat foraged during recording in a dark room encountered for the first time (Extended Data Fig. 7f). In this experiment, a 150 cm diameter circular arena was used, as described in a previous study^64^. The arena was encircled by thick, dark blue curtains. All light sources in the recording room were turned off or occluded before the recording started.

### Linear track task

Five rats with MEC-parasubiculum implants, of which two also had hippocampal implants, shuttled back and forth on a 200 cm linear track with liquid rewards delivered at each end (chocolate-flavored oat milk dispensed via a tubing system). When the rat consumed a reward at one end of the track, the reward port at the opposite end was refilled. Before surgery, rats were trained on the track task until they reliably completed ∼40 laps in one training session. Recording sessions lasted 45-66 minutes, with 18-48 minutes of running between reward sites.

### Wagon-wheel task

Two rats with MEC-parasubiculum implants were tested in a “wagon-wheel” track – an elevated 10-cm-wide circular track with two perpendicular cross-linking arms spanning the circle’s diameter^5^. The track was fitted with 8 reward wells, placed halfway between each of the 5 junctions. Each of the wells could be filled with chocolate oat milk via attached tubing. At any given time, a pseudorandom subset of 1-4 of the wells was filled to encourage steady exploration of the entire maze. Before surgery, the rats were trained to asymptotic performance levels, where they obtained at least 30 rewards per 30-minute session.

### Natural sleep

Sleep recordings including both REM and slow-wave sleep (SWS) epochs were obtained from 9 animals with MEC-parasubiculum implants (2 of them with combined MEC-hippocampal implants). Sleep was promoted by putting the rat in a black acrylic box (40 × 40-cm floor, 80 cm height), lined with towel on the floor, before recording. The box walls were transparent to infrared, allowing the rat’s position and orientation to be tracked through the walls. Water was available ad libitum. During recording, room lights were on and pink noise was played through the computer speakers to mask background sounds. Sleep sessions typically lasted 2–3 h (see Extended Data Fig. 9a for an example recording).

### Spike sorting and single-unit selection

Spike sorting was performed using KiloSort 2.5^27^, with customizations as previously described^5^. To exclude low-firing units, fast-firing interneurons and contaminated clusters, units were excluded if they had a mean spike rate of less than 0.1 Hz or greater than 10 Hz (<0.025 Hz or >5 Hz for hippocampal units), or if their waveforms had a large spatial footprint (i.e. similar waveform amplitude across a wide range of channels), as this seemed to be a reliable indicator of poor cluster quality^27^. The waveform footprint was expressed as the anatomical spread of recording channels where the unit was detected (with detection defined as at least 10% of the unit’s maximal amplitude across all channels), weighted by the unit’s waveform amplitude on each channel^27^:

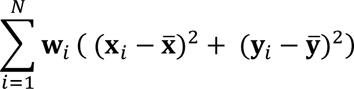

where [**x***_i_*, **y***_i_*] refers to the [*x*, *y*]-location of the *i*^*th*^ recording channel, 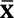 and 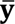 are the centers of mass of the recording channel positions where the unit was detected, **w***_i_* is the amplitude of the waveform at the *i*^*th*^ recording channel, and *N* is the number of channels where the unit was detected. Units were excluded if the spatial footprint of their waveforms exceeded 35 μm (50 μm for hippocampal units). Units that were recorded on sites located outside the regions of interest (MEC-parasubiculum or hippocampus) were excluded from further analysis.

### Preprocessing and temporal binning

During awake sessions, only time epochs in which the rat was moving at a speed above 5 cm/s were used for spatial analyses. Spike times were binned in 10 ms time bins for all population analyses (unless otherwise specified), and tracking data was resampled at the same time intervals to align it with the spike-count data. For computational reasons, wake sessions were truncated in length to the nearest multiple of 100 seconds (given by the chunk size for latent manifold tuning model), by trimming the tail end of the behavior session.

### Rate maps and angular tuning curves

To generate 2D rate maps for the open field arena and wagon wheel track, position estimates were binned into a square grid of 2.5 × 2.5-cm bins. For each bin, we calculated each cell’s firing rate (number of spikes in the bin divided by time spent in the bin). Rate maps were smoothed with a cross-validated smoothing procedure. Briefly, the recording was split into 10 folds of equal duration, and the firing rate *y* during each fold was compared with the expected firing rate 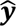 based on the rate map calculated over the remaining 9 folds and smoothed with a gaussian kernel of width σ. The value of σ (1 *cm* < σ < 50 *cm*) that minimized the mean squared error of the firing rate prediction (using the MATLAB function *fminbnd*) was chosen to smooth the rate map. The same procedure was used to compute spatial rate maps with respect to latent position signals from the LMT model (see section *latent manifold tuning model*). For population vector (PV) decoding analyses (see below), a fixed-width gaussian kernel was used to smooth the rate maps (*σ* = 7.5cm). Spatial autocorrelations and grid scores were calculated as described previously^36^, based on the individual cells’ rate maps.

Angular tuning curves with respect to head direction, theta phase or internal direction (see section *Decoding of internal direction based on population vector correlations*) were calculated by binning the angular variable into 60 evenly spaced angular bins. For each 6-degree bin, spike rate was calculated as the number of spikes divided by time spent in the bin. Angular tuning curves were smoothed with the same cross-validated smoothing procedure as the spatial rate maps (0.01 rad < σ < 1 rad) (0.01 *rad* < σ < 1 *rad*), except in PV-decoding analyses, where a fixed-width gaussian kernel of *σ* = 12 degrees was used.

### Identification of grid cells and grid modules

Grid cells were detected as groups of cells corresponding to grid modules by finding clusters of co-recorded cells that expressed similar spatially periodic activity in the open field, based on a similar procedure described previously^5^. Briefly, 2D autocorrelograms were calculated from the coarse-grained spatial rate maps of each cell (10 × 10-cm bins, no smoothing across bins). Autocorrelogram bins within a central radius of 2 bins or beyond an outer radius corresponding to the rate-map size were masked, before vectorizing and concatenating the autocorrelograms in a matrix. Considering the spatial autocorrelograms of all cells as a point cloud, where each point (autocorrelogram) has an *N-* dimensional position representing the value of each of its *N* spatial bins, the Manhattan distances between all points were calculated, and each point’s 30 nearest neighbors were identified. The resulting neighborhood graph was given as input to the Leiden clustering algorithm, which was used to partition the spatial autocorrelograms into clusters, using a resolution parameter of 1.0 (1.5 for sessions with >1000 units). Clusters that contained cells with clear and consistent grid patterns were classified as candidate modules of grid cells. For each cluster, grid periodicity was measured by the grid score of the median autocorrelogram across all cells in the cluster. Grid pattern consistency was measured by computing the Pearson correlation between the average autocorrelogram of the cluster and each individual cell’s autocorrelogram. The median consistency across all cells in the cluster was defined as the cluster’s grid consistency. For a cluster to be classified as a grid module, three criteria had to be fulfilled: (1) cluster grid score greater than 0.3, (2) grid pattern consistency greater than 0.5 and (3) the cluster needed to contain a minimum of 10 cells. In some recording sessions, single grid modules appeared to be split into two clusters with similar spacing and orientation. For this reason, we added a step that merged grid clusters if the correlation between their average autocorrelograms was greater than 0.7.

### Subclasses of grid cells defined by differential bursting

To measure the tendency of cells to fire in bursts, we devised a burst score (BS) based on the firing-rate autocorrelograms of each cell (time range: ±50ms, bin width: 1 ms). The autocorrelogram value of the center bin was set to zero, and the autocorrelogram counts were normalized by the mean. Next, we compared the autocorrelogram values at short time lags (2-10ms) with those found at longer time lags (13-50ms):

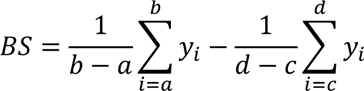

where *y_i_* is the value of the i-th bin of the mean-normalized autocorrelogram, and [a, b, c, d] are the indices of autocorrelogram bins corresponding to 2, 10, 13 and 50ms time lags. Thresholds for classification of bursty vs. non-bursty cells were determined by inspection of the distribution of burst scores across the sample (bimodal with a trough around -0.2; Extended Data Fig. 4a). Cells with a burst score > 0.0 were classified as bursty and cells with burst scores < -0.4 were classified as non-bursty, while cells with burst scores from [-0.4, 0.0] were left unclassified.

While grid cells could be classified into a bursty and non-bursty subclasses based on burst scores, a third subclass of grid cells was revealed by closer inspection of autocorrelogram shape or theta phase modulation (Extended Data Fig. 4a). To reliably identify these three subclasses in an unsupervised manner, we applied an unsupervised clustering algorithm to the temporal autocorrelograms of all grid cells^5^. Temporal autocorrelograms were computed, for each cell, by calculating a histogram of the temporal lags between every spike and all surrounding spikes within a ±100 ms window, using 2 ms bins. The histogram was then divided by the mean value, and concatenated in a matrix, discarding any cells with less than 100 counts in the autocorrelogram. PCA was applied to the matrix of autocorrelograms, treating individual cells as observations and keeping the first 10 components for further analysis. Next, a neighborhood graph was constructed by computing the Manhattan distance between all pairs of points in the 10-D point cloud and finding each point’s 150 nearest neighbors. The graph was used as input to the Leiden clustering algorithm (resolution parameter 0.2). The clustering algorithm detected three distinct clusters with unique autocorrelogram shapes, in agreement with the three subclasses of grid cells identified upon inspection of the theta phase modulation and burst scores across all grid cells (Extended Data Fig. 4a). Most cells in the first two clusters had positive burst scores (referred to as bursty type I and bursty type II), while cells in the third cluster had negative burst score (referred to as non-bursty).

### Classification of direction-tuned cells

Cells were classified as tuned to head direction (HD) if their HD-tuning curves differed significantly from a uniform distribution (p<0.001, Rayleigh test for non-uniformity) and was stable across the first and second half of the recording session (p<0.01, Pearson correlation between tuning curves from first and second half). No further criteria on tuning strength were applied to classify HD tuning, since many internal direction-tuned cells display weak HD-tuning and there was no clear cut-off between HD-tuned cells and non-HD-tuned cells (Extended Data Fig. 5a). Cells were classified as internal direction cells if their tuning curves, with respect to decoded internal direction (see section *Decoding of instantaneous position and direction based on population vector correlations*), passed the above criteria for non-uniformity and stability and had a mean vector length > 0.3. Tuning width was defined as two standard deviations of the tuning curves.

### Theta phase estimation

Theta phase was extracted from the population spiking activity of all units (including fast-firing, putative interneurons) within the MEC-parasubiculum region. Spike times were binned into 10-ms bins and the resulting spike counts were bandpass-filtered with a 2^nd^ order Butterworth filter in the theta range (5-10 Hz). Principal components analysis (PCA) was applied to the matrix of bandpass-filtered spike counts (with units as variables and time bins as observations). The first two principal components typically contained an oscillating circular representation corresponding to the theta rhythm. Theta phase at time *t* was defined as the direction of the projection of the population vector at time *t* onto the plane defined by PC1 and PC2. The phase with minimal firing activity was defined as zero. In the 6 rats with dual entorhinal-hippocampal implants, hippocampal sweeps were referenced to theta phase extracted from MEC-parasubiculum activity. In the two rats with probes only in the hippocampus, theta phase was estimated from hippocampal population activity.

### Theta cycle skipping

Theta skipping, i.e. the tendency for cells to fire on every other theta cycle, was quantified by computing a theta-skipping index (TSI) as in previous descriptions of the phenomenon^10,40,41^. Briefly, firing-rate autocorrelograms were generated for each cell (time range: ±500 ms, bin width: 5 ms, gaussian smoothing: σ=10 ms), and the relative height of the second theta peak *p*_2_compared to the first theta peak *p*_1_ was determined as

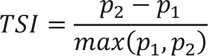

where *p*_1_ is defined as the maximum autocorrelogram value between lags 90 and 170 ms and *p*_2_ as the maximum value between lags 180 and 300 ms. Cells were classified as theta-skipping if the theta-skipping index was positive, i.e. if the second theta peak was higher than the first peak. The theta-skipping index only has a clear interpretation for cells that are theta-rhythmic and is thus reported for cells that were classified as modulated by theta phase. Cells were classified as theta phase modulated if their theta phase tuning curves differed significantly from a uniform distribution (p<0.001, Rayleigh test for non-uniformity) and were stable across the first and second half of the recording session (p<0.01, Pearson correlation between tuning curves from first and second half). Cells were classified as non-rhythmic if they did not meet the above criteria and had theta phase mean vector length <0.2.

### Decoding sweeps based on population vector correlations

This section describes how we visualized and quantified sweeps. We first decoded position from the activity of the entire set of MEC-parasubiculum neurons. For each temporal bin, we correlated the instantaneous population activity (a population vector, PV) with the session-averaged population activity for each location in the environment (reference population vector, rPV). The inputs to the PV correlation decoder were (1) a N×T matrix of temporally smoothed firing rates (gaussian kernel σ=10 ms), with T columns corresponding to time bins and N rows corresponding to neurons, and (2) a N×M matrix of spatial tuning curves, with M columns corresponding to position bins and N rows corresponding to neurons. The spatial tuning curves were normalized by dividing each neuron’s tuning curve by its mean value. To decode position across successive time bins, we computed the Pearson correlation coefficient between each PV (columns of the spike count matrix) and each rPV (columns of the tuning curve matrix). This yielded a vector of correlation values for each time step, with elements corresponding to each [*x*, *y*]-location in the environment. The decoded position was taken as the position bin with the highest correlation value, and the resulting decoded trajectory was smoothed with an σ=8 ms gaussian kernel. To filter out unreliable estimates, the peak correlation value for each time bin was compared to a shuffled distribution of PV correlation values (computed by shuffling rows of the tuning curve matrix). Decoded estimates were discarded if their PV-correlation did not exceed the 99^th^ percentile of the shuffled distribution. Decoding estimates were also discarded for timepoints where fewer than 5 cells were active.

Within each theta cycle, the decoded position swept outwards from a location slightly behind the tracked head position of the animal (Fig. 1). Thus, the starting locations for a series of sweeps formed a slowly evolving trajectory that roughly followed the animal’s running trajectory. This trajectory, referred to as the lowpass-filtered decoded trajectory and estimated by decoding position from spikes emitted in the beginning of each theta cycle, was used as a reference signal for measuring sweeps. Spike counts from the first half of each theta cycle were smoothed with a wide gaussian kernel (σ = 1.7 theta cycles) before the PV-correlation decoding method was used to decode position across all bins. The resulting decoded trajectory was smoothed with a σ=10ms gaussian kernel. Sweeps seemed to be more reliably anchored to the lowpass-filtered decoded trajectory compared to the tracked head position of the animal. This difference was particularly evident when rats navigated in darkness, when sweeps and lowpass-filtered signals could deviate substantially from the rat’s actual trajectory (Extended Data Fig. 7f).

### Extracting sweep trajectories

Individual sweeps, defined as smooth spatial trajectories within each theta cycle, were extracted from the decoded position trajectory by a simple sequence detection algorithm. Within each theta cycle, candidate sweeps were identified as the longest sequence (highest number) of consecutive valid time bins where the decoded position jumped less than 20 cm and changed direction less than 90 deg between consecutive 10 ms bins. Candidate sweeps were truncated to maximize the net Euclidian distance from beginning to end, which effectively removed folds at either end of the sweep. A sweep vector, ***s***, was defined as the vector between the lowpass-filtered decoded position at the beginning of the theta cycle and the distal-most point of the candidate sweep. To identify sweep trajectories that were fairly straight and well described by the sweep vector, we measured, for each candidate sweep, the goodness of fit *r*^2^ between the sweep vector axis and the collection of [*x*, *y*]-points in the sweep, represented by vectors **x**, **y**:

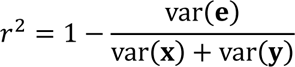

where **e** is a vector of residuals of all points with respect to the sweep vector axis, such that **e***_i_* is the residual of point [**x***_i_*, **y***_i_*] . Candidate sweeps were kept for further analyses if they included at least 4 samples and had a *r*^2^ > 0.5 (meaning that most of the variance is explained by the sweep axis). Sweep direction and sweep length were defined as the direction and magnitude of the sweep vector.

For several analyses and visualizations (e.g. Fig. 1b), sweeps (in allocentric coordinates) were transformed to head-centered coordinates. This was done by first subtracting the tracked position (e.g. Fig. 1b) or lowpass-filtered decoded trajectory (e.g. Extended Data Fig. 2e) and then rotating each [*x*, *y*]-coordinate by the animal’s head direction. Session-averaged sweeps (e.g. Fig. 1b) were computed by first interpolating each head-centered sweep trajectory at 50 time points linearly spaced from the beginning to the end of the trajectory. Interpolated sweeps were grouped into those that followed a right sweep or left sweep, before an average sweep was computed for each group by taking the median position at corresponding time points within the sweep (Fig. 1b, 4e, 5d; Extended Data Fig. 2e-f).

### Decoding of internal direction based on population vector correlations

Internal direction was decoded from the activity of all MEC-parasubiculum neurons using the same PV-correlation procedure described for position decoding but using angular tuning curves for head direction instead of spatial position rate maps. Since we observed that internal direction cells were activated in discrete pulses during each theta cycle (Fig. 2c), the time bin corresponding to the theta phase with maximal activity was used to express internal direction within each theta cycle. The decoded internal direction α at time *t* was taken as the circular mean of all possible decoding angles weighted by the correlation values for each directional bin at time *t*:

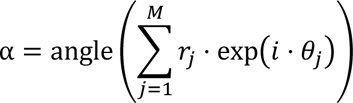

where θ*_j_* denotes the angular value of the *j*^*th*^ bin, *i* is the imaginary unit, *r_j_* is the correlation value for the *j*^*th*^ angular bin and *M* is the total number of angular bins.

For some analyses and visualizations (e.g. Fig 1d), we rotated the decoded internal direction (in allocentric coordinates) to a head-centered reference frame by subtracting the animal’s head direction.

### Left-right alternation of sweeps and internal direction

The extent of directional alternation across successive sweeps and internal direction signals was characterized in a head-centered reference frame (see above) during periods when the animals moved faster than 15 cm/s. The data were generally thresholded at 15 cm/s because alternation was more reliable during running (Fig. 6e-f; see Fig. 6e-f for data at speed thresholds down to 5 cm/s). The prevalence of directional alternation was computed by counting triplets of theta cycles where sweep direction or internal direction alternated in a left-right-left or right-left-right pattern (detected as sign-inversions in the angles between successive directions), divided by the total numbers of theta cycle triplets where sweeps or internal direction were detected. The fraction of theta cycle triplets with directional alternation was compared to a shuffled distribution of scores where head-centered directions were randomly shuffled (1000 iterations). Directional alternation was visualized in temporal autocorrelograms of angles between successive head-centered sweep direction or internal direction. Autocorrelograms were computed as the circular correlation between the original trace of head-centered directions and a series of lagged versions of the signal (lags from -7 to 7 theta cycles). To find and visualize directional modes of sweep and internal direction angles, we computed histograms of head-centered directions that were conditioned on the decoded direction in the previous theta cycle. First, decoded directions were classified as left or right-directed based on the sign of their angular offset from the previous cycle. Next, two histograms were computed, one for decoded directions that followed a left-directed angle and one for decoded angles that followed a right-directed angle. Since sweeps and internal direction angles alternate reliably, this procedure resulted in two unimodal distributions, one on either side of the reference head direction. The directional modes of sweep or internal direction were taken as the peak of each of the two conditional distributions.

In the artificial agent simulation, a time-resolved measure of directional alternation was used to quantify instantaneous alternation of the agent’s chosen sweep direction. A three-sweep sliding window was used, such that at the *i*^th^ timestep a triplet of sweep directions α*_i_*_−1:*i*+1_ was selected. Within the three-sweep sliding window, the two angles between consecutive sweep pairs were calculated: *a* = α*_i_* − α*_i_*_−1_ and *b* = α*_i_*_+1_ − α*_i_* . The alternation score *s* was hence computed as:

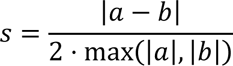

where |·| denotes the absolute value. The alternation score ranged from 0-1, where 1 indicates perfect

alternation.

### Single-module decoding

To decode position from individual grid modules, the PV-correlation decoding analysis (first applied on all cells) was next applied to subsets of cells that belonged to individual grid modules. The decoded position from each grid module was mapped onto the grid module’s hexagonal unit tile using a crosscorrelation procedure. First, we computed a template grid pattern for each grid module by averaging the spatial rate-map autocorrelograms across all cells in the module. The template grid pattern was crosscorrelated with the 2D distribution of PV-correlation values at each time step. Peaks in the resulting spatial crosscorrelogram were detected, and for each crosscorrelogram the peak nearest to the origin was taken as the decoded position. To ensure that the decoded position was within the bounds of the central unit tile, the decoded trajectory was wrapped around the three grid axes of the template grid pattern. Single-module sweeps were detected as described for whole-population decoding (i.e. by finding consecutive time bins within each theta cycle where the decoded trajectory formed a smooth trajectory), except that spatial and directional offsets between consecutive decoded positions were computed with periodic boundary conditions derived from the template grid pattern. For some visualizations (Fig. 1e, 4b) the single-module sweeps were aligned to the behavioral trajectory of the animal by subtracting the offset between the animal’s tracked position and the lowpass-filtered decoded trajectory at the beginning of each theta cycle.

### Identification of putative excitatory connections between cells

Putative excitatory monosynaptic connections between pairs of co-recorded neurons were identified by detecting short-latency, short-duration peaks in the firing rate cross-correlograms (CCGs), following previous procedures^66–68^. CCGs were computed with 1 ms bins over a ± 50 ms window. A baseline CCG was computed by convolving raw CCGs with a hollowed gaussian kernel (σ=5 ms, hollow fraction 60%), which has been shown to approximate a jittered CCG^66^. Neuron pairs recorded on the same probe, where each of the neurons emitted at least 2000 spikes and the raw CCG contained at least 1000 counts, were tested for functional connections. The baseline CCG was subtracted from the raw CCG and subsequent peak detection was performed on the baseline-corrected CCG. Neuron pairs were classified as putatively connected if the highest positive CCG peak satisfied the following five conditions: (1) The peak occurred within an asymmetric time range consistent with monosynaptic excitation (0.7 – 4.7 ms); (2) The peak height exceeded 5 standard deviations of the baseline-corrected CCG; (3) The peak’s p-value was smaller than 0.001 (estimated from a Poisson distribution with continuity correction, as in ref.^66^); (4) the peak’s width (defined as the set of bins contiguous with the peak whose values exceeded half of the peak height or 2 s.d. of baseline and had a p-value<0.01) was less than 3 ms (consistent with the precise spike-timing expected from a monosynaptic connection); (5) The peak’s width did not overlap with the zero-lag bin (suggestive of common input). Candidate connections were also discarded if any of the non-peak bins exceeded 2.5 standard deviations of the baseline-corrected CCG or if any of the bins in the anticausal direction had p-values <0.01.

Overall connection probability was computed by dividing the total number of putatively connected neurons by the total number of pairs that were checked for connections. Similarly, target-specific connectivity rates were computed by dividing the number of connections from one functional cell class to another by the number total number of pairs.

### Decoding internal direction with PCA or UMAP

The internal direction signal was also decoded in an unsupervised manner, with principal components analysis (PCA) or uniform manifold approximation and projection (UMAP^69^). In this approach, the high-dimensional neural activity was projected down to a 2D subspace to characterize the trajectory of population activity on a low-dimensional manifold. Only rhythmic direction-tuned cells (with head direction mean vector length MVL > 0.3 and theta phase MVL > 0.3) were included in the analysis, to avoid interference between the ring-like manifold of interest and other ensemble representations (e.g. spatial signals from pure grid cells). Spike counts from *n* neurons were binned into *t* time bins corresponding to individual theta cycles. Theta cycle time bins were used instead of 10ms time bins to prevent global within-cycle firing rate fluctuations from driving the results. Principal components analysis was performed on the resulting *t*-by-*n* matrix. Next, internal direction was either read out (1) directly from the PCA output, or (2) by applying a second dimensionality reduction step on the data with UMAP (Extended Data Fig. 3c-d).

In the PCA decoder, internal direction was read out by projecting the neural data onto the first two eigenvectors and taking the arctangent of each [*x*, *y*]-coordinate in the resultant 2D projection. Because these angles were arbitrarily rotated with respect to the environment, they were aligned to the environment before further analysis. We assumed that internal direction and head direction had equal mean directions, and subtracted the average difference between the two signals from the decoded signal to align it with respect to the environment.

In the UMAP decoder, scores of the 20 principal components with the largest explained variance was used as input to the nonlinear dimensionality reduction algorithm UMAP with hyperparameters: n_components=3, metric=correlation, n_neighbors=199, min_dist=0.3, init=spectral. This yielded a 3-D embedding of the high-dimensional population activity. The 3-D UMAP point cloud typically showed a clear circular shape. The 3-D point cloud was then collapsed to 2-D by projecting the points onto the best-fit 2-D plane. Hence, the 2-D points were converted into angles by taking the arctangent of each x- and y-coordinate. The resulting angles were aligned to the environment by subtracting the average offset from head direction, as described above.

### Bayesian decoding of position

Sweeps could also be decoded using Bayesian reconstruction (Extended Data Fig. 3a). Position was decoded at 10ms time steps from the matrix of firing rates and tuning curves from *N* neurons, with an assumption of Poisson firing and a flat position prior^70^:

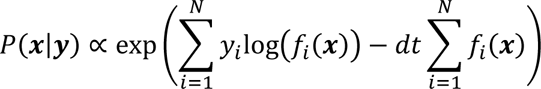

where *P*(***x*|*y***) is the conditional probability for the rat’s 2D location ***x***, given the observed spike count *y* and position tuning curves *f*(***x***). The decoded position was taken as the position bin that maximized *P*(***x*|*y***).

### Latent manifold tuning model

The PV-correlation method has two limitations. First, it can only be used to decode positions and directions that the animal has physically sampled, because it relies on tuning curves with respect to the animal’s tracked position and head direction. This is particularly a problem for hippocampal data, where the neural correlation structure varies between environments and brain states^71–73^, making it impossible to decode position in one environment based on reference tuning curves from a different context. A second challenge is that the sharp spatial and directional tuning of grid cells and internal direction cells is obscured in standard time-averaged reference tuning curves, since sweeps and internal direction deviates from tracked position and head direction (Extended Data Figs. 3f, 4d). To simultaneously extract sweeps through unvisited space and characterize spatial tuning directly from neural activity, we therefore adapted the latent manifold tuning (LMT) model introduced by Wu et al. 2017^45^. In this framework, sweeps may be considered as hidden or ‘latent’ trajectories on a neural manifold that is not directly observable. The goal of the LMT model is to infer (1) the latent trajectory and (2) each cell’s tuning to locations on the manifold, based on neural population activity. The model assumes that the latent variable evolves smoothly with time, that individual neurons are smoothly tuned to locations on the manifold, and that neurons fire according to a Poisson process (see Extended Data Fig. 8 for schematic and ref^45^ for details). For a multidimensional (vector-valued) latent variable x(t), the temporal evolution of component j is modelled as a Gaussian process:

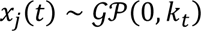

where *k*_*t*_ is a temporal covariance function *k*(*t*, *t*^′^) ≜ cov(*x_j_*(*t*), *x_j_* (*t*′)). In this case, the exponential kernel (*t*, *t*^′^) = *r* exp(−|*t* − *t*^′^|/*l*) is used, with variance *r* and length-scale *l* respectively controlling the amplitude and smoothness of the latent variable. The log tuning curves *f*(**x**) are also modelled as Gaussian processes, with the log tuning of the *i*^*th*^ neuron expressed as:

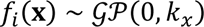

where *k*_*x*_ is a spatial covariance function, in this case a Gaussian kernel 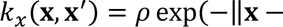 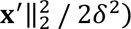 with variance ρ and length-scale δ. The value of the latent variable **x**(*t*), in conjunction with each cell’s tuning curve *f_i_*(**x**), predicts the cell’s log-firing rate, which is then transformed with an exponential nonlinearity into a Poisson-distributed spike count *y_i_*_,*t*_ as below:

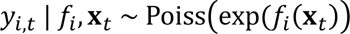

The predicted and observed spike trains are compared, yielding a log-likelihood value. The model was fitted using an expectation-maximization algorithm that maximizes the log-likelihood of the spike trains by separately optimizing the latent variable and tuning curves in alternation. Over multiple iterations of this two-step optimization procedure, both evolve to capture the latent dynamics in the neural population activity, thus improving prediction of the observed spikes.

The original LMT framework uses a single latent variable to predict the neural activity; however, as a form of Poisson regression, the LMT model can be trivially combined with other Poisson regression models, as a means of extracting other factors of interest, or to regress out noise. In the present work, we formulated the activity of each neuron as a sum of log firing-rate contributions from five input variables:

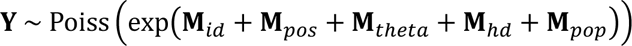

Where Y is the time-by-neurons matrix of predicted spike counts and **M**_*varname*_ is a time-by-neurons matrix of log firing-rate contributions from model *varname*. The first two contributions correspond to the two latent variables of interest: *internal direction* (**M**_*id*_) and *position* (**M**_*pos*_). *Internal direction* was modelled as a 1D latent circular variable (hyperparameters: ρ=0.1, *r*=100, δ=0.5, *l*=0.1), that was initialized with the animal’s tracked head direction. *Position* was modelled as a 2D latent variable (hyperparameters: ρ=0.1, *r*=10, δ=6, *l*=0.015) that was initialized with the animal’s tracked position.

The final three covariates (*theta phase* **M**_*theta*_, *head direction* **M**_*hd*_, and *population firing rate* **M**_*pop*_) are known to modulate MEC-parasubiculum activity^4,19,36,74^, and are here included as ‘noise’ covariates to regress out their substantial contributions to the neural activity, hence reducing the likelihood of them influencing the extracted latent variables. *Theta phase* and *head direction* were modeled as circular 1D variables in the LMT framework; however, the variables were respectively fixed at the values of measured theta phase and head direction, and only tuning curves were optimized (hyperparameters: ρ=1000, δ=2 for theta phase and ρ=0.1, δ=0.5 for head direction). The *population firing rate* model was implemented as a generalized linear model **M**_*pop*_ = **x**_*pop*_β_*pop*_, where **x**_*pop*_ is a column vector of log population firing rates, and **β**_*pop*_ is a row vector of learned coefficients for all neurons. Population firing rate was computed as the average instantaneous firing rate across neurons, smoothed with a σ=20ms gaussian kernel.

Each step of fitting in the composite model consisted of serially updating the parameters for each of the five sub-models by maximizing the log likelihood of the model. Since the latent trajectories may be arbitrarily rotated and distorted with respect to the physical environment, an alignment procedure was performed after model fitting^45^. The latent internal direction trajectory was aligned to the tracked head direction of the animal by subtracting the average angle between the two signals. The latent position trajectory was aligned to the tracked position of the animal with an affine transformation. The fitted latent variables were used for most LMT-based analyses of sweeps and internal direction. For some visualizations (e.g. Fig. 4d) and analyses during sleep, position and direction was decoded from neural activity and fitted LMT tuning curves using the Bayesian framework described in previous section. For single-module analyses during sleep (Fig. 4, Extended Data Fig. 9) and in Extended Data Fig. 3b, the LMT model was fit separately on the activity of grid cells in individual grid modules.

### GLM-based single-cell tuning model

Theta sweeps introduce offsets between the animal’s current location and the location represented by place and grid cells, resulting in smeared receptive fields when a cell’s spikes are plotted as a function of animal location (Extended Data Fig. 4e-f). To characterize the spatial tuning of individual cells independently, in a manner that accounts for sweeps, we used a Poisson Generalized Linear Model (GLM) to model the spike train of a cell, *y*, as a function of a set of explanatory variables, **X**, parametrized by the learned parameters ***β***:

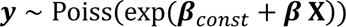

#### Construction of the data matrix X

The model included four explanatory variables: position, head direction, internal direction (from LMT) and theta phase. The variables were expressed by using a ‘basis-expansion’ procedure, by which a set of smooth basis functions was used to decompose each single variable into multiple variables. Each basis function’s weighting was given by a corresponding ***β*** parameter, giving the GLM flexibility to fit any smooth function of the input variable in question. The 2D position variable was expressed by a set of 2D Gaussian basis functions (σ = 2 cm) arranged in a 10-cm-spaced triangular grid which tiled the open field arena and a surrounding ‘buffer zone’. The angular variables head direction, internal direction and theta phase were expressed by a set of 50 Von-Mises functions (κ = 10) with equally spaced mean values from 0 to 2π. Basis expansion was performed by evaluating each of the basis functions for a given value of the input variable in question.

The basis-expanded representations are high-dimensional and multicollinear (i.e. the basis function values are correlated). In a regression model, these attributes tend to cause overfitting; hence we used PCA to produce a low-dimensional, orthogonal representation of the basis-expanded data matrix. PCA, which acted as a form of regularization. For position, the principal components explaining 99% of the variance were retained, reducing the dimensionality from 527 to 92. For head direction and theta phase, the principal components explaining 80% of the variance were retained, reducing the dimensionality from 50 to 6. After performing basis expansion and dimensionality reduction for the input variables, the resultant matrices for all input variables were concatenated into the GLM design matrix X:

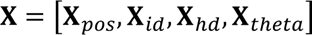

#### Theta-phase-dependent shifting of the position covariate

To model the effect of sweeps on position-modulated firing, we added a pre-processing step which applied a theta phase-dependent shift to the animal’s tracked position coordinates. Specifically, the 2D position coordinates (***x***) were parametrically shifted by a distance δ along the internal direction axis (α):

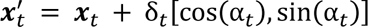

where ^′^ denotes the shifted position coordinates. The shift quantity, δ_*t*_, at each time point was modelled as a function of the current theta phase Specifically, the shift quantity δ was modelled as a function shift parameters ***γ***, fitted by the model, and the basis-expanded theta phase **X**_*theta*_:

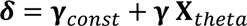

The shifting parameters ***γ***_*const*_ and ***γ*** were fitted together with the GLM **β** parameters using a gradient-based solver with the finite-difference method (MATLAB function ‘*fminunc’*).

Across cells and recordings, the GLM shift model yielded sharper receptive fields than standard rate maps with respect to tracked position (Extended Data Fig. 7i). The model’s estimates of position tuning were more robust than those from the LMT model, because the latter depended on large numbers of co-recorded spatially modulated cells. Therefore, rate maps based on the GLM-shifted position were deemed most appropriate to use for identifying grid cells and grid modules (see section *Identification of grid cells and grid modules*).

### Sweeps through unvisited space

For analyses of sweeps and spatial tuning to never-visited locations outside the bounds of the wagon-wheel track, we first defined the area of space that the animal had visited. This was achieved by binning the 2D environment in 2.5 cm bins and finding all bins that the rat had visited (resulting in a binary 2D map with values of 1 for visited bins and 0 otherwise). The bounds of animal’s coverage was found by applying a binary dilation operation of the occupancy map (Matlab function *imdilate* with a disk-shaped structuring element, radius=1) followed by a morphological closing operation (Matlab function *imclose* with a disk-shaped structuring element, radius=1). The zero-valued bins in the resulting occupancy map were defined as never-visited.

### Sleep stage classification

Sleep stages were identified as described in previous studies^5,19^. First, we identified periods of sustained immobility (longer than 120 s, locomotion speed below 1 cm/s, head angular speed below 6 deg/sec). These periods were subclassified into SWS and REM based on delta- and theta-rhythmic population activity in the recorded cells. Population firing rate was computed by summing the binarized 10ms spike counts from each cell. The rhythmicity of this aggregated firing rate with respect to delta (1–4 Hz) and theta (5–10 Hz) frequency bands was quantified by applying a zero-phase, fourth-order Butterworth band-pass filter and then calculating the amplitude from the absolute value of the Hilbert transform of the filtered signal, followed by smoothing (Gaussian kernel with σ = 5 s) and standardization (z-scoring). Periods for which the ratio of the amplitudes of theta and delta activity (theta/delta ratio) remained above 5.0 for at least 20 s were classified as REM. Periods during which theta/delta ratio remained below 2.0 for at least 20 s were classified as SWS (Extended Data Fig. 9B).

### Detection of sweep and internal direction signals during sleep

To decode sweeps and internal direction from neural activity during sleep, we used tuning curves (LMT) from open-field sessions from the same recording day as the sleep session. Position was decoded separately for individual grid modules (see section *single-module decoding*), since the correlation structure of grid cells across brain states may be preserved within but not across modules^5,19^. Since the theta rhythm is absent during slow-wave sleep (SWS), we used local maxima in population activity as reference points for analysis of sweeps and direction signals in all brain states. To detect local maxima in the population activity, regardless of brain state, the spike counts of all internal direction cells was summed and smoothed with a Gaussian kernel (σ = 20ms), before applying the Matlab function *findpeaks* with default parameters to detect peaks in the summed activity. Local maxima occurred at theta-rhythmic intervals during wake and REM and irregularly during SWS (Extended Data Fig. 9). Pairs of local maxima 2–250 ms apart were used to quantify directional alternation in all brain states, while all detected maxima were used to measure alignment between sweep and direction signals. Internal direction was taken as the decoded direction at the time of local maxima. To extract sweeps, we identified smooth sequences of decoded positions (sequences where decoded position jumped less than 15% of grid spacing and changed direction less than 2 radians between successive time bins) that occurred in windows centered around each of the local maxima in population activity. The windows extended 50 ms to either side of local maxima or to the edge of the neighboring window. Since spatial representations were decoupled from physical movement during sleep (Extended Data Fig. 9g), sweep trajectories were referenced to the low-pass-filtered decoded trajectory (smoothed with a 100 ms gaussian kernel) and aligned to a “virtual head direction” (low-pass-filtered decoded direction, σ = 1 theta cycle gaussian smoothing).

### Simulation of an ideal sweep-generating agent

To test the hypothesis that alternating sweeps are controlled by an algorithm that maximizes the sampling of surrounding space, we simulated a sweep-generating agent that maximized environmental sampling by choosing sweep directions that minimized overlap with previous sweeps.

First, we modeled the spatial coverage of a single sweep. Since grid modules express sweeps at multiple spatial scales, we reasoned that the total spatial coverage of a sweep may be considered as a sum of sweeps across individual grid modules (Extended Data Fig. 10a). A model sweep footprint was formulated, based on previous empirical observations of geometric relationships between single-module grid patterns^16^, and the geometric properties of sweeps in the present results (Fig. 1e-g). Briefly, we summed the grid patterns of five idealized grid modules, with an inter-module scale ratio of √2 and gaussian-shaped grid fields with σ = 1⁄6 of each module’s spacing, at offsets from the origin corresponding to typical single-module sweep lengths of 1⁄3 of module spacing. The sum of sweeps across modules resembled a torch beam radiating outwards: as distance from the origin increases, the footprint broadens and decays in intensity (Extended Data Fig. 10a). We approximated this shape by multiplying two simple spatial functions: (1) an inverse distance function and (2) an angular weighting function taken from a Von Mises distribution.

If we let *d* and θ denote the distance and direction from the agent’s [*x*, *y*]-position **x_*agent*_** to a location **x** in the environment (*d* = ǁ **x** − **x_*agent*_** ǁ and θ = *arctan*(**x** − **x_*agent*_**)), the intensity of the sweep footprint at location **x** for a chosen sweep direction α becomes:

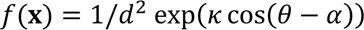

where κ is the angular concentration parameter of a Von Mises distribution, which determines the angular width of the sweep footprint. A value of κ = 5 was used initially (Fig. 6a) to reflect the empirically derived sweep-shape (Extended Data Fig. 10a), but a parameter search revealed that stable alternation emerged across a range of κ-values (Extended Data Fig. 10c).

To run the simulation of sweeps on a linear track, we created an artificial scale-free 2D environment, binned into a 401-by-401 square grid. The agent was moved along a linear path at constant speed and asked to generate a sweep every time step, by placing a sweep footprint in a specified direction. The cumulative trace of sweeps ℎ at time *t* was computed by summing the footprints of previous sweeps:

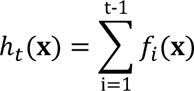

The optimal sweep direction α_optimal_ at time t was chosen by finding the angle α that minimized the spatial overlap between the current sweep *f* and the cumulative trace of previous sweeps ℎ:

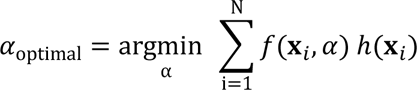

where the spatial overlap was computed by multiplying the current sweep footprint and the cumulative sweep trace and summing across all *N* spatial bins.

Next, the simulation was run using the recorded behavioral trajectory of rats running in the open field. Real-world positions were mapped onto the agent’s simulated environment by setting the agent’s bin size to 1 cm and placing the open field at the center of the bin grid. The agent’s time steps were yoked to the times of theta cycles in the experimental data, and for each theta cycle, the agent deployed the above algorithm to select the optimal sweep direction. To prevent the agent from being influenced by sweeps that occurred at similar locations far in the past, we introduced a temporal decay factor *r* (range: 0-1), that exponentially discounted the intensity of the cumulative coverage trace at each time step:

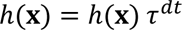

Since the agent’s sweep-direction choices are determined solely by its previous decisions, we call this the “free” version of the model. We also formulated a “hybrid” version, where the agent was tasked with predicting the optimal sweep at each time step, given the directions of previous sweeps decoded from neural data. This was implemented by using the latent internal direction (LMT), instead of the agent’s past sweep directions, to compute the cumulative sweep trace. Four animals with were excluded from these analyses, since internal direction could not be reliably estimated in these animals.

### Histology and recording locations

The rats received an overdose of pentobarbital, after which they were perfused intracardially with saline followed by 4% formaldehyde. The brains were extracted and stored in 4% formaldehyde, and later cut in 30-µm sagittal or coronal sections with a cryostat. The sections were Nissl-stained with cresyl violet and probe shank traces were identified in photomicrographs. In 14 animals, recording sites on the probes targeting MEC-parasubiculum were aligned to the histological sections as done previously^5^ by using as reference points (1) the tip of the probe shank and (2) the intersection of the shank with the brain surface. The aligned shank map was then used to calculate the anatomical locations of individual recording sites (Extended Data Fig. 1). Estimates of anatomical locations are subject to some degree of measurement error, due to the limited accuracy of the alignment process and the fact that units may be detected some distance away from the recording site.

### Data analysis and statistics

Data analyses were performed with custom-written scripts in Matlab and Python. Clustering analyses of grid cell modules and bursting subtypes of grid cells were conducted using the python package Scanpy^75^ and its dependencies. The latent manifold tuning model and the functional connectivity analyses were implemented by adapting publicly available code from Wu et al.^45^ and Spivak et al.^68^, respectively. Statistical analysis was performed in Matlab. Circular statistics were computed using the Circular Statistics Toolbox^76^. Results are reported with means ± s.e.m. unless otherwise indicated. Statistical tests were nonparametric and two-tailed, unless otherwise indicated. The Mann-Whitney U test was used for unpaired comparisons, and the Wilcoxon signed-rank test was used for paired comparisons. Pearson correlations were used unless otherwise indicated. Power analysis was not used to determine sample sizes. The study did not involve any experimental subject groups; therefore, random allocation and experimenter blinding did not apply and were not performed.

### Data availability

The datasets generated during the current study will be available at… (to be provided before publication).

### Code availability

Code for reproducing the analyses in this article are available at… (to be provided before publication).

## Extended Data Figures

**Extended Data Fig. 1:**
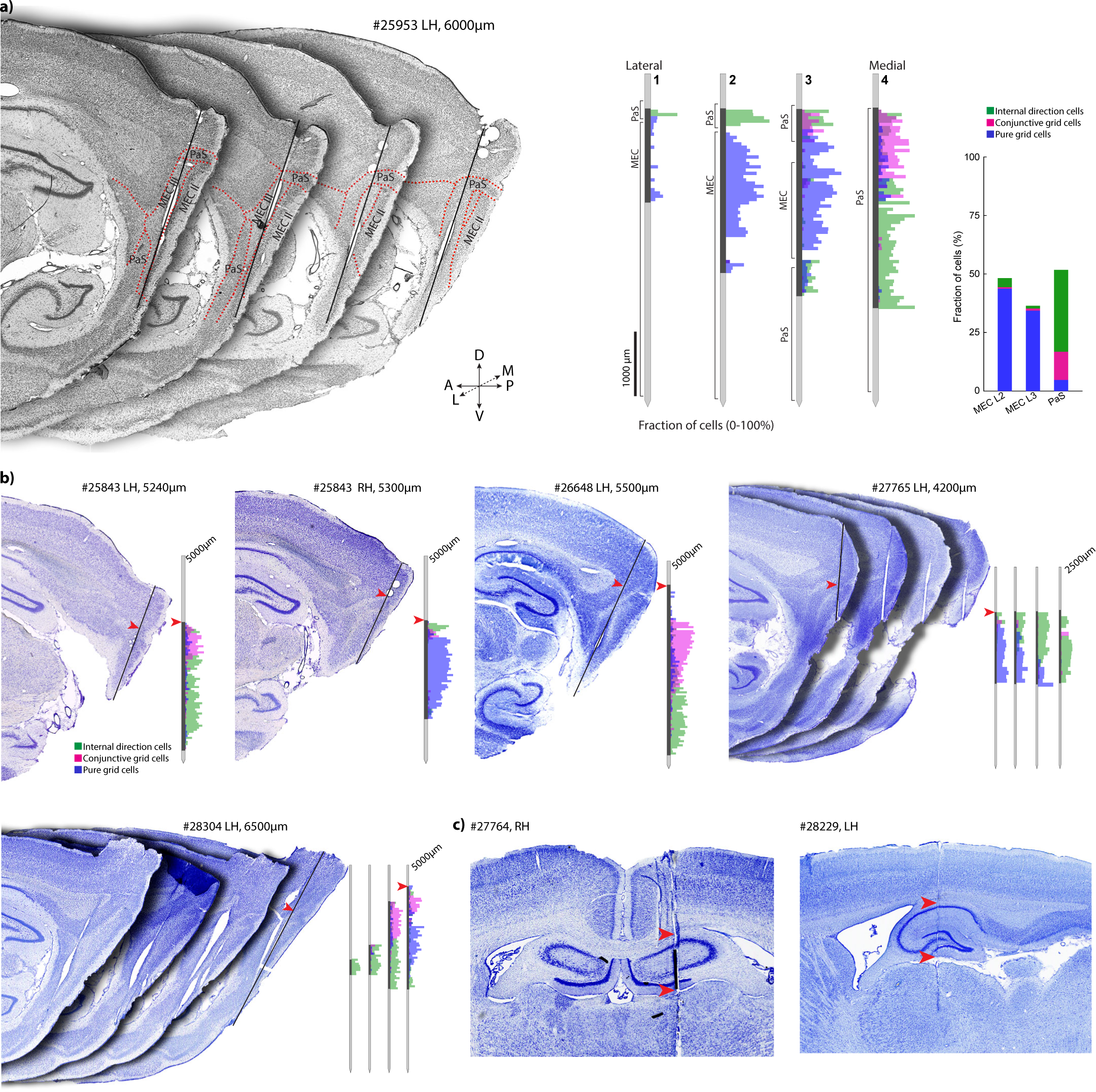
Recording locations and anatomical segregation of functional cell types. **a.** Anatomical separation of rhythmic direction-tuned cells (‘internal direction cells’), conjunctive grid cells and pure grid cells. Left: Serial sagittal sections from rat 25953 (also shown in 2b) showing tracks from a 4-shank Neuropixels 2.0 probe. Sections are organized from lateral to medial and each section contains the track from one of four recording shanks (∼250 µm apart; section with clearest track is shown). Borders between brain regions are indicated with dashed lines (MEC, medial entorhinal cortex; PaS, parasubiculum). Middle: Density of internal direction cells (green), conjunctive grid cells (pink) and pure grid cells (blue) along each of the probe shanks, estimated by counting the number of functional cells recorded in 50 μm bins along the probe divided by the total number of recorded cells in each bin. To maximize anatomical coverage of the distribution plot, cell counts are pooled across 7 recording sessions with different configurations of active recording sites (only 384 out of 5,120 sites can be recorded at any given time). Black portions of probe shanks show sites that were recorded from. Right: Percentage of recorded cells in MEC and parasubiculum belonging to each of the three functional cell classes (MEC layer II: 43.7% pure grid cells, 0.7% conjunctive grid cells, 3.8% direction-tuned cells; MEC layer III: 34.4% pure grid cells, 0.9% conjunctive grid cells, 1.1% internal direction cells; PaS: 4.8% pure grid cells, 12% conjunctive grid cells, 34.9% internal direction cells; 27,413 cells from 14 rats, 2-7 recording sessions per rat). Note that most pure grid cells in the sample were located in MEC layers II and III (3,377/3,861 or 87.4%), while most of the conjunctive grid cells (1,224/1,285 or 95.3%) and internal direction cells were located in parasubiculum (3,547/3,796 or 93.4%). **b.** Sagittal histological sections with probe tracks for four additional representative animals with probes in MEC-parasubiculum (reconstructions were performed for 14 of the 16 animals with MEC-parasubiculum implants). Animal identity and hemisphere are indicated above each section for the four animals. For each implanted probe shank, the section with the clearest track in MEC-parasubiculum is shown. Insets show, for each animal, the density of internal direction cells, conjunctive grid cells and pure grid cells along the probe shanks (as in **a**) across multiple recordings (range: 2-4). Red arrowheads indicate the most dorsal enabled recording site for each probe. Note that functional cell types are anatomically segregated in most animals. **c.** Histological sections (coronal and sagittal) showing two representative examples of recording locations in hippocampus. Arrowheads indicate the dorsoventral range of recording sites that were included in the study.

**Extended Data Fig. 2:**
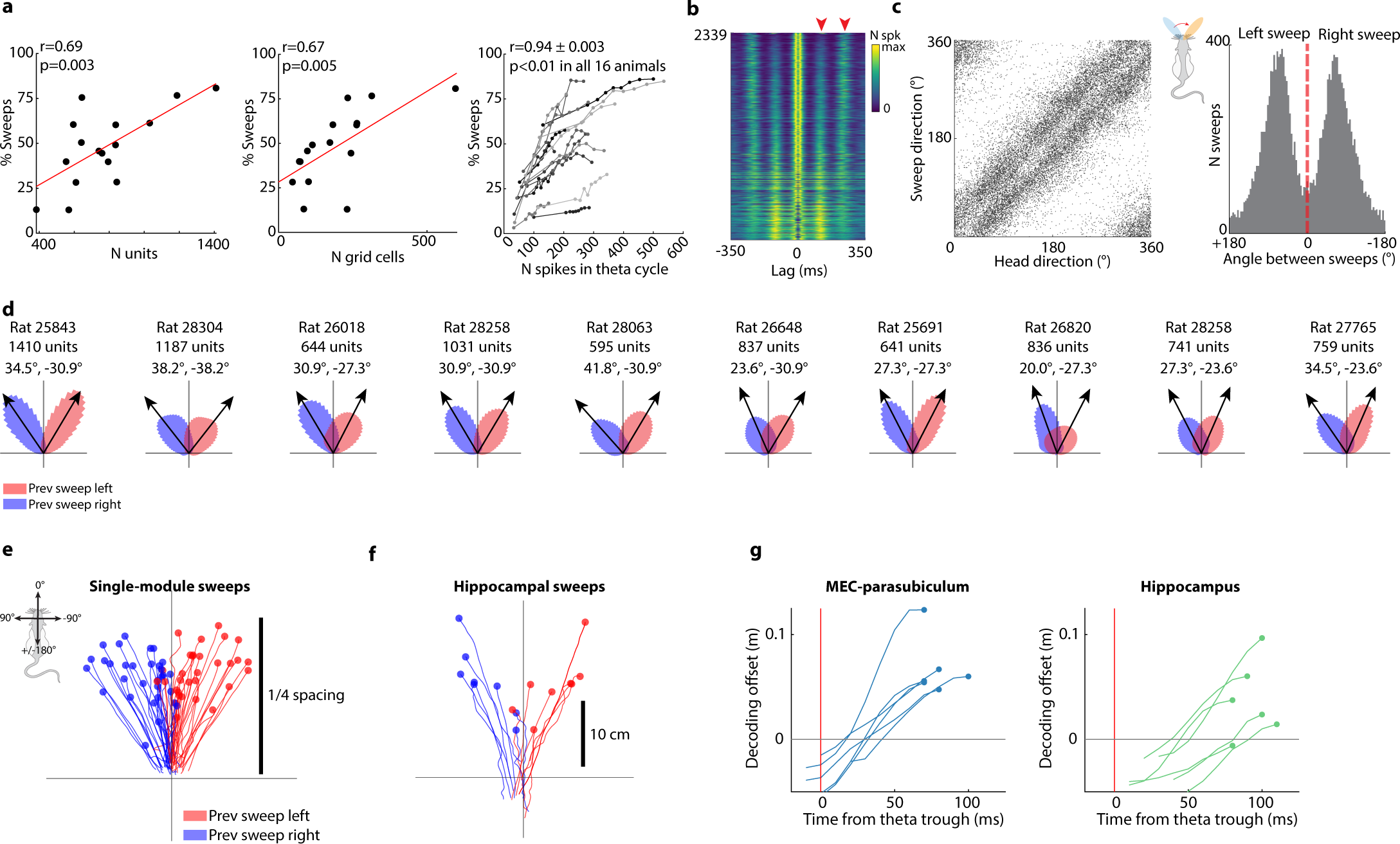
Left-right alternating sweeps in grid cells and place cells. **a.** Number of detected sweeps scales with number of recorded MEC-parasubiculum cells (left), number of grid cells (middle), and number of spikes per theta cycle (right), suggesting that the reported fraction of theta cycles with sweeps is underestimated. Data from individual animals are plotted as dots (left and middle) or lines (right). **b.** Stacked firing-rate autocorrelograms (±350 ms) for all theta-modulated grid cells (2,399/3,194 or 75.1% of grid cells were theta-modulated), sorted by tendency to fire on alternating theta cycles. Firing rates are color-coded. Lags corresponding to repetitions of the ∼8 Hz theta rhythm is indicated with red arrowheads. Note that a large fraction of theta-modulated grid cells (1,289/2,399 or 53.7%) display theta skipping, with prominent peaks at lags corresponding to every other theta cycle. **c.** Left: Scatter of directions of all sweeps and tracked head direction at the corresponding times during one example recording session (same session as Fig. 1a). Sweep directions are distributed bimodally around the animal’s head direction (circular correlation between sweep direction and head direction r=0.55 ± 0.007, p<0.001 in all 16 animals, absolute mean offset 3.1 ± 0.7 deg, mean ± s.e.m. across 16 animals). Right: Histogram of angles between successive sweeps. Note that these angles are clustered around ±60 deg. Sweeps directed to the right or left of the previous sweep are defined as ‘right’ and ‘left’ sweeps, respectively. **d.** Circular histograms of head-centered sweep directions for the 10 animals with the highest fraction of theta cycles with detected sweeps. Sweeps are rotated to head-centered coordinates (head orientation is vertical) and sorted based on whether the previous sweep was directed to the right (blue) or left (red). Note clustering of sweeps around two principal head-centered directions offset ∼30 deg to the left and right of the animal’s head direction. **e.** Sweeps averaged across theta cycles for all cells of a grid module, as in Fig. 1b, but now showing all modules. Sweeps are plotted with reference to the lowpass-filtered decoded trajectory (origin) and rotated to head-centered coordinates (head orientation is vertical). Sweep lengths are normalized by the grid spacing of each module. Note that sweeps have similar lengths relative to the scale of the grid cells. **f.** Averaged sweeps decoded from hippocampal ensemble activity (8 rats), plotted as in Fig. 1b. Hippocampal sweeps alternated from side to side in 78.5 ± 1.1% (mean ± s.e.m.) of successive triplets of theta cycles with detected sweeps, significantly more than when sweep directions were shuffled (>99.9^th^ percentile for all animals). **g.** Forward progression of decoded sweeps (projected onto the rats’ head axis) as a function of time from the beginning of each theta cycle for MEC-parasubiculum (left) and hippocampus (right) for all 6 rats with dual HC/MEC implants, plotted as in Fig. 1i. Data from each of the rats is plotted as separate lines (1 session per rat). Note that hippocampal sweeps are delayed relative to MEC sweeps.

**Extended Data Fig. 3:**
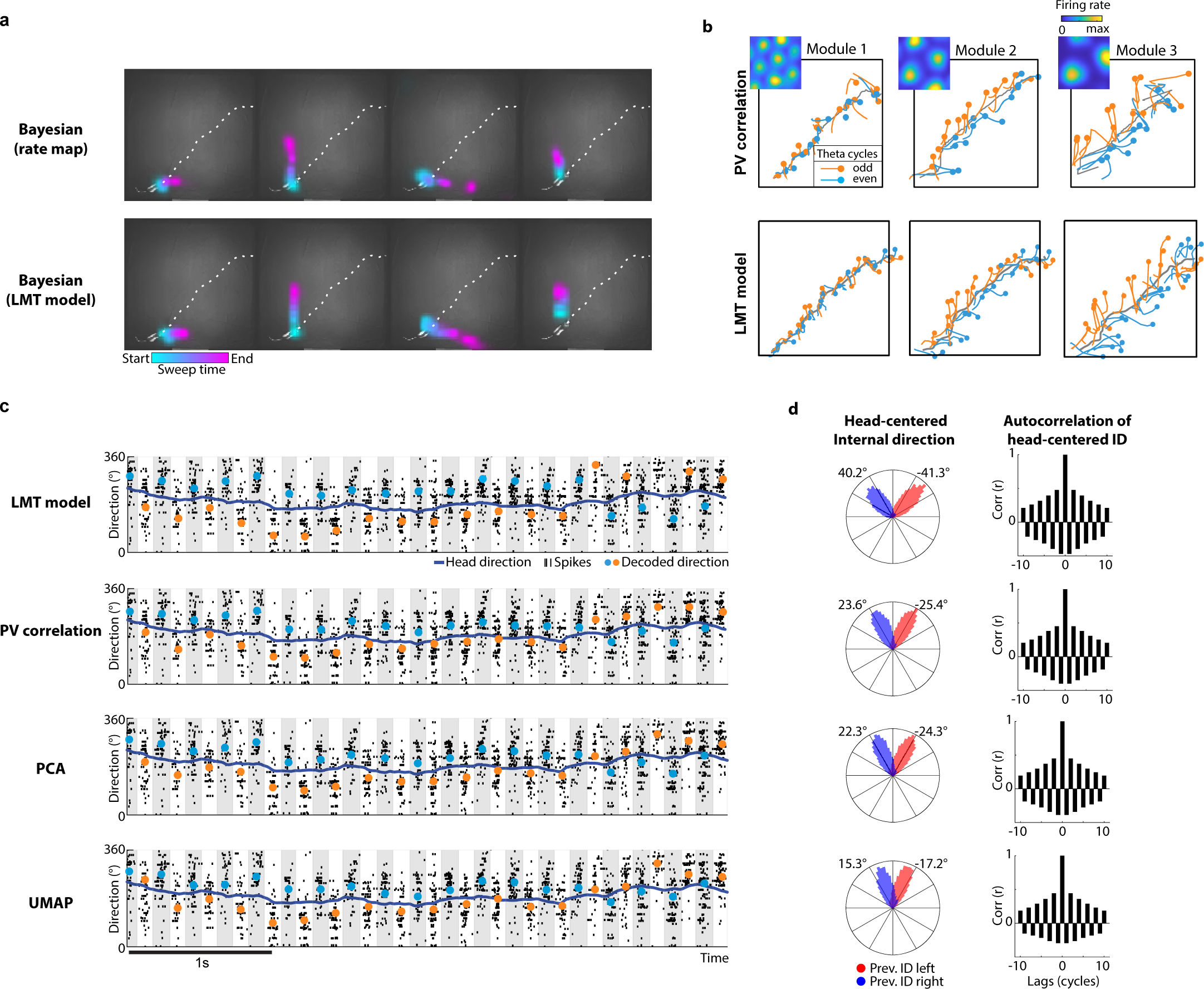
Alternating sweeps and directional signals can be decoded with several methods. **a.** Example sweeps decoded from all co-recorded MEC/PaS cells (same time period as Fig 1a) using different decoding methods. The Bayesian decoder was trained with either standard rate maps (top row) or the fitted tuning curves from the latent manifold tuning model (bottom row; Fig 4, Extended Data Fig. 8). Sweeps are similar with all methods, but can be extracted with higher fidelity by the LMT model. **b.** Single-module sweeps extracted by PV-correlation decoding (top; as in Fig. 1e) and by the LMT latent variables (bottom). Note that the two methods yield similar results. **c.** Left: Alternating sequences in decoded internal direction are similar regardless of the decoding method used. In each plot, rows of black ticks indicate spike times from 533 rhythmic directional cells, with cells vertically sorted by preferred direction (as in Fig. 2c). Colored dots show the extracted population signal using 4 different decoding methods (colors indicate odd and even-numbered theta cycles). Note that all methods extract a left-right alternating directional signal that follows the angles of the packets of spiking activity. **d.** Left: Circular distribution of head-centered internal direction (as in Fig 2d). Right: auto-correlograms of internal direction (as in Fig 2e) for each of the decoding methods.

**Extended Data Fig. 4:**
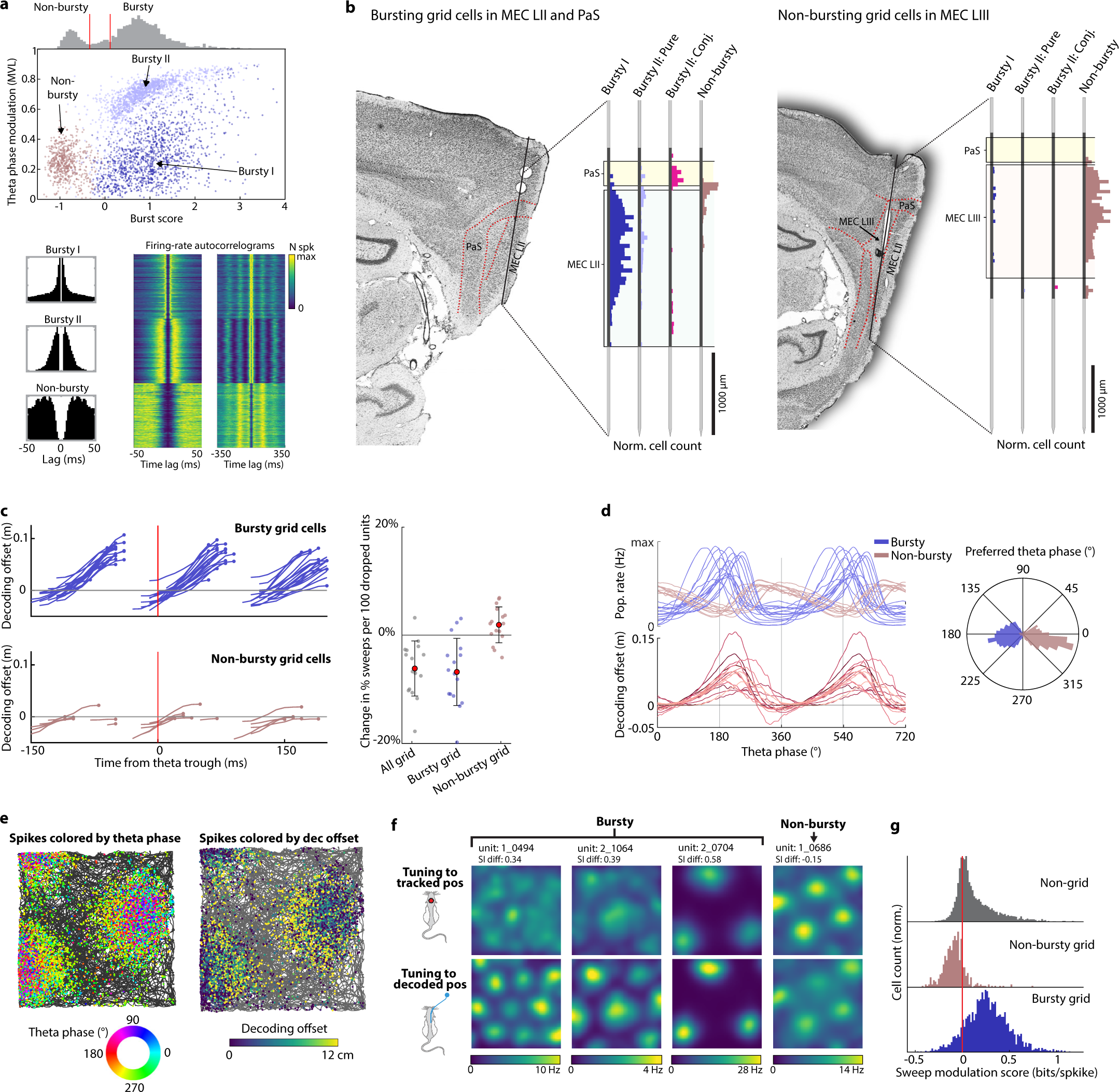
Bursty MEC layer II grid cells are strong carriers of the sweep signal. **a.** Three subclasses of grid cells with distinct temporal dynamics are revealed by clustering of firing-rate temporal autocorrelograms. Top: Scatter plot showing burst scores (Methods) and theta phase modulation for all recorded grid cells (dots). Note three separable clusters of grid cells. Two of the subclasses fire in bursts (“bursty I” and “bursty II”) and can be dissociated from each other based on autocorrelogram shape and theta modulation. ‘Bursty I’ grid cells include non-directional ‘pure’ grid cells in MEC layer II. ‘Bursty II’ grid cells include most conjunctive grid cells in parasubiculum and some non-directional ‘pure’ grid cells in MEC layer II. The third subclass (‘non-bursty’) does not fire in theta-rhythmic bursts. Histogram above scatter shows bimodal distribution of burst scores in the cell sample, with thresholds for binary classification (bursty or non-bursty) indicated by red vertical lines (cells with burst scores between the two lines were unclassified). Bottom: One example autocorrelogram (±50 ms) from each grid cell subclass is shown to the left. Middle and right: Stacked autocorrelograms (±50 ms and ±350 ms) from 500 example grid cells in each subclass. Firing rate is color-coded. Cells are sorted by cluster identity (top to bottom). Note prominent theta-rhythmic activity in bursting grid cells. **b.** Anatomical segregation of grid-cell subclasses (see **a** for reference). Left: Histological section from rat 25843 with electrodes in MEC layer II and parasubiculum (labelled ‘PaS’). The proportions of recorded cells belonging to each grid-cell subclass at each location along the probe are shown as vertical histograms. Bursty II grid cells with conjunctive direction tuning (labelled ‘Conj.’) are plotted separately from pure bursty II grid cells. Note predominance of bursty grid cells in MEC layer II and parasubiculum. Right: Histological section from rat 25953 with electrodes in MEC layer III and PaS. Note predominance of non-bursty grid cells in MEC layer III. Based on inspection of probe tracks in 14 animals, 84.3% or 2,316/2,747 of bursty I-II grid cells (excluding conjunctive grid cells) were found to be in MEC layer II, 76.5% or 724/946 of non-bursty cells were in MEC layer III, and 92.7% or 1,240/1,337 of conjunctive grid cells were in parasubiculum (data from 2-7 sessions per animal). **c.** Left: Progression of sweeps decoded separately from bursty grid cells (top) and non-bursty grid cells (bottom) visualized by plotting the offset between decoded position and lowpass-filtered trajectory projected onto the rats’ head axis and averaged across all theta cycles in each session. Position was decoded from a random subsample of 100 bursty or non-bursty grid cells in all sessions with more than 100 co-recorded bursty grid cells (9 sessions from 10 animals, 1-3 sessions per animal) or non-bursty grid cells (5 sessions from 3 animals, 1-3 sessions per animal). Each line corresponds to one session. Sweeps are more prominent when decoding from bursty grid cells. Right: Decrease in number of detected sweeps when omitting all grid cells or individual grid-cell subclasses from the decoder input data (all 16 animals, 1 session per animal, plotted as individual dots). Values are normalized by comparing the number of sweeps obtained when omitting a size-matched random selection of non-grid cells (repeated 100 times per session) and dividing by the number of dropped cells. Means and s.d. are shown as red dots and whiskers. Excluding all grid cells (47-650 cells) or bursty grid cells (27-567 cells) results in a significant drop in the number of detected sweeps (all grid cells: -6.2 ± 0.3% per 100 dropped units, p=0.0008; bursty grid cells: -6.8 ± 0.4%, p=0.002, Mann-Whitney U test). Excluding non-bursty grid cells (3-244 cells) did not result in a significant change in the number of detected sweeps (+1.9 ± 0.2%, p=0.12). **d.** Bursty grid cells are maximally active when decoded position sweeps outwards. Top: Firing rate for bursty and non-bursty grid cells during the course of the theta cycle (summed across all co-recorded cells, one line per session). Bottom: Offset between decoded position and lowpass-filtered decoded position as a function of theta phase (each line corresponds to one animal). Right: Polar histogram of preferred firing phase for all bursty and non-bursty grid cells. Note that bursty and non-bursty grid cells are active at opposing phases of the theta cycle^32,77^. **e.** Out-of-field spikes in bursting grid cells coincide with outgoing sweeps. Panels show firing locations for an example grid cell with dots corresponding to the animal’s location at the time of individual spikes and color corresponding to theta phase (left) or offset between decoded position and tracked position (right). Note that out-of-field spikes occur at phases where the decoded position sweeps ahead of the animal (180-270 deg with 0 deg as the phase of minimum activity). **f.** Spatial receptive fields of bursty grid cells can be sharpened by accounting for sweeps. Example rate maps plotted with respect to tracked position (top row) and rate maps with respect to decoded position (bottom row) for three bursty grid cells and one non-bursty grid cell (columns). Note that for bursty cells, grid patterns are clearer and sometimes only visible when activity is plotted as a function of decoded position, which sweeps outwards from tracked position, indicating that the cells are tuned to the position of the sweep rather than the animal’s tracked location. Difference in spatial information content (‘SI diff’) between the original and corrected rate maps is noted above each cell and indicates the strength of sweep-modulation. **g.** Sweep-modulation is primarily associated with grid-cell identity. Sweep modulation scores (difference in spatial information content between the original and corrected rate maps) are higher for bursty grid cells than for non-bursty grid cells (sweep modulation score for bursty grid cells: 0.27 ± 0.23, mean ± s.d. for n=2,766 cells; non-bursty grid cells: -0.08 ± 0.13 (n=460 cells); non-grid cells, i.e. all putative excitatory cells excluding grid cells; 0.14 ± 0.23 (n=9,702 cells); 16 animals, 1 session per animal; Mann-Whitney U=31.5 p<0.001 (2.1e-218) for bursty vs. non-bursty cells and Mann-Whitney U=28.8 p<0.001 (5.5e-182) for bursty vs. non-grid cells).

**Extended Data Fig. 5:**
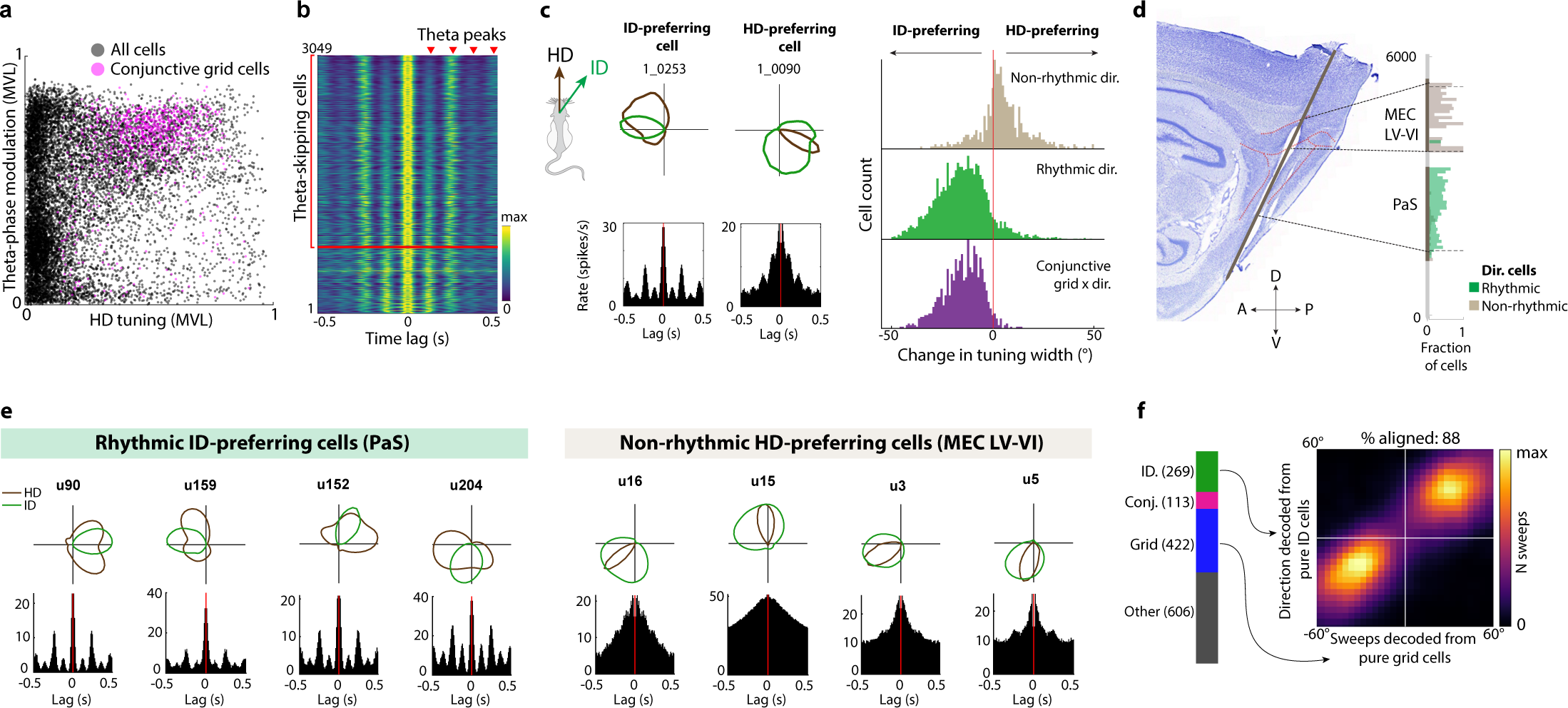
The internal direction signal is expressed by a discrete population of cells in parasubiculum. **a.** Left: Scatter plot of head direction (HD) and theta-phase tuning strength (mean vector length, MVL) for all recorded cells (12,300 cells, 16 animals, 1 session per animal). Each dot corresponds to one cell. Conjunctive grid cells are plotted in pink (23.3% or 848/3,632 of all direction-tuned cells were conjunctive grid cells). Note broad range of directional tuning and high proportion of theta-rhythmic cells (83.9% or 3,049/3,632 of all direction-tuned cells and 93.6% or 794/848 conjunctive grid cells were significantly modulated by theta oscillation phase). **b.** Stacked firing-rate autocorrelograms (±500 ms), color-coded by firing rate, for all theta-modulated direction-tuned cells, sorted by tendency to fire on alternating theta cycles. Lags corresponding to repetitions of the ∼8 Hz theta rhythm are indicated with red arrowheads. Cells above the red horizontal line have positive scores and are thus classified as ‘skipping’ cells. Note that most cells (73.9% or 2,253/3,049) display theta skipping, with prominent peaks at lags corresponding to every other theta cycle. **c.** Left: Directional tuning curves with respect to head direction (HD, brown) and decoded direction (green) for two example cells. Bottom: Temporal autocorrelogram of cells’ firing rates. The theta-rhythmic cell is sharply tuned to the decoded internal direction (ID) signal, while the non-rhythmic cell is more strongly tuned to HD. Right: Histograms showing relative tuning width to HD vs ID from cells with significant tuning to either HD or ID (4,628 cells from 16 animals). Theta-rhythmic directional cells are more sharply tuned to ID than HD (90.5% or 3,527/3,897 of rhythmic directional cells were ID-preferring; difference in tuning width, defined as two standard deviations of the tuning curve, to ID vs HD: -15.4 ± 13.4 deg, mean ± s.d., p<0.01 Wilcoxon signed rank test), while non-rhythmic cells are more strongly tuned to HD (73.4% or 408/556 of non-rhythmic directional cells were HD-preferring; difference in tuning width: 4.8 ± 12.1 deg, p<0.01, Wilcoxon signed rank test). Conjunctive grid cells were mostly ID-preferring (96.5% or 950/985, tuning width difference: -14.9 ± 9.5 deg, mean ± s.d., p<0.01 Wilcoxon signed rank test). This indicates that there are separate populations of directional cells that encode distinct signals. **d.** Sagittal histological section from rat 28304 with recording sites in parasubiculum and MEC layer V-VI, showing the track of a probe shank that went through deep layers of MEC before entering parasubiculum (PaS). Right: Distribution of rhythmic and non-rhythmic directional cells along the probe shank. Note the high proportion of non-rhythmic directional cells in MEC layer V-VI. **e.** Four example rhythmic directional cells recorded from parasubiculum (left) and four non-rhythmic directional cells recorded from the deep layers of MEC (right) in the animal shown in **e**. Tuning curves and autocorrelograms are plotted as in **d**. Note that the rhythmic cells are more strongly tuned to ID, while the non-rhythmic cells are sharply tuned to HD (resembling classical HD cells). **f.** Alternating sweep and direction signals are expressed by separate neural populations. Left: Stacked bar chart showing the number of functional cells recorded in an example animal (25843). Right: Heat map showing alignment between direction of sweeps decoded from pure grid cells (n=422) and internal direction decoded from simultaneously recorded pure internal direction (‘ID’) cells (n=269), both expressed in head-centered coordinates as in Fig. 2g. Conjunctive grid cells were excluded from the decoder input data. Decoded sweep and direction signals remain aligned (both left or both right) in 88.1% of theta cycles where sweeps and direction signals were detected (correlation between sweep and direction signals: r=0.84, p<0.0001), suggesting that the observed dynamics is not caused by using the same cells to decode position and direction.

**Extended Data Fig. 6:**
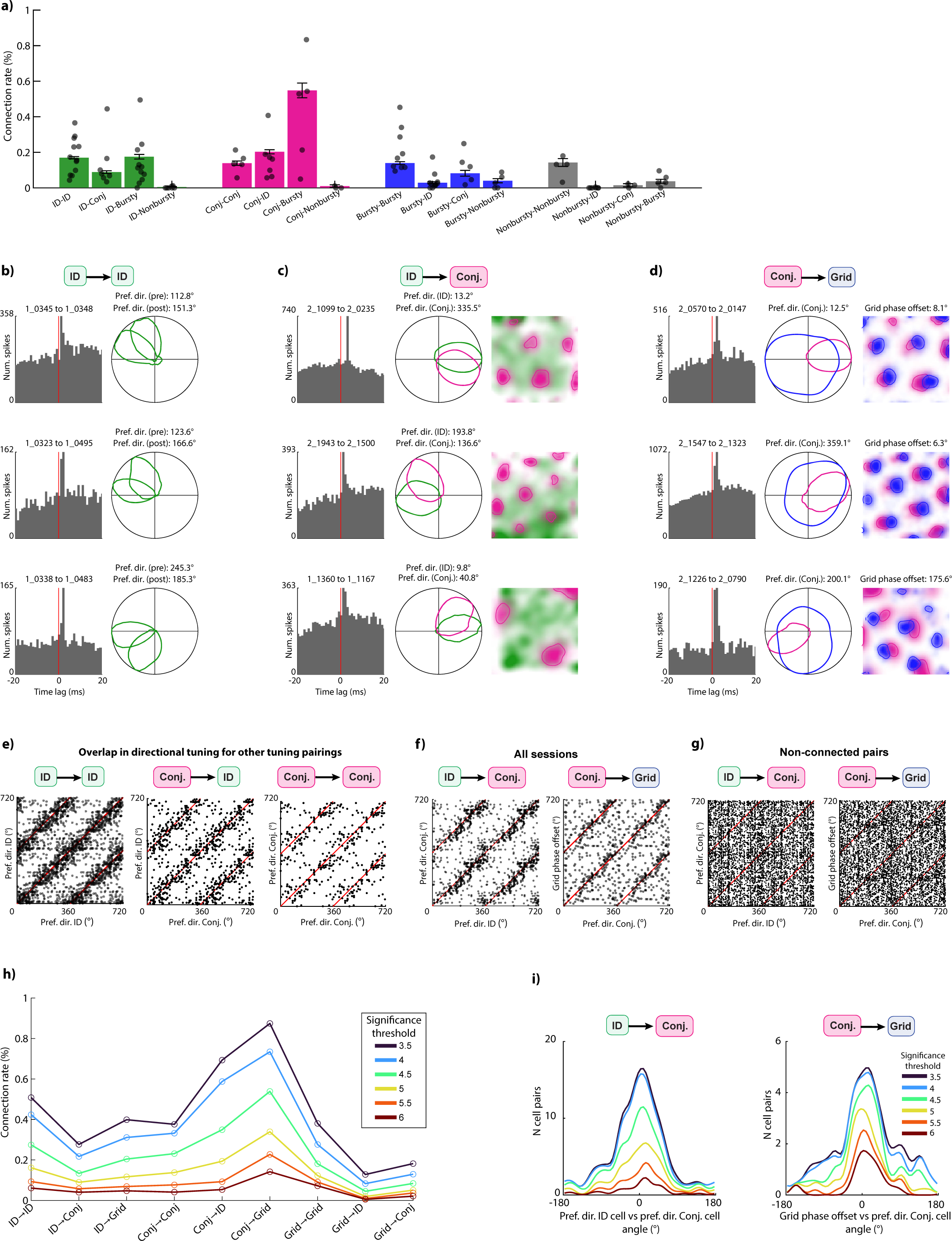
Connectivity between functional cell types. **a.** Estimated connection probabilities for all combinations of cell types. Bars show average connection rate across animals (as in Fig. 3b) and dots show connection rates in individual animals. Note that internal direction (ID) cells and conjunctive grid cells have putative connections to bursty pure grid cells, but not non-bursty pure grid cells (connection rates for direction-tuned to bursty: 239/124,914 or 0.191±0.012%; ID to non-bursty: 4/63,404 or 0.006±0.003%; conjunctive to bursty: 188/32,433 or 0.580±0.042%; conjunctive to non-bursty: 3/19,858 or 0.015±0.009%). Although estimated connection rates varied from animal to animal, the general pattern of connections was preserved. **b.** Examples of putative recurrent connections between internal direction (ID) cells. Each of the three rows shows firing rate cross-correlogram (left column) and directional tuning curves (right column) for an example pair of putatively connected ID → ID cells. Preferred firing direction (PFD) of each cell is indicated above directional tuning curves. Note similar directional tuning between connected cells. **c.** Three example cell pairs illustrating putative connections between ID-tuned cells (green) and conjunctive grid cells (pink). Plotted as in Fig. 3c. **d.** Three example cell pairs showing putative connections between conjunctive grid cells (pink) and pure grid cells (blue). Plotted as in Fig. 3f. **e.** Alignment of directional tuning for other putatively connected cell pairs. Scatter plot shows preferred directions of pairs of pre- and postsynaptic cells, plotted as in Fig. 3d. Note that all combinations of connected direction-tuned cells have similar directional tuning (angle between preferred directions: 12.1 ± 58.0 deg, correlation: r=0.49, p<0.001 for 726 putative ID → ID pairs; angle: 4.6 ± 58.2 deg, correlation: r=0.48 p<0.001 for 323 putative conjunctive grid → ID pairs; and angle: 11.1 ± 58.0 deg, correlation: r=0.52 p<0.001 for 132 putative conjunctive grid → conjunctive grid pairs). **f.** Alignment of directional tuning is present across all recording sessions. Left: Preferred directions for all pairs of connected ID cells to conjunctive grid cells (angle between preferred directions: 8.4 ± 51.6 deg, correlation: r=0.56 p<0.001, n=351 pairs from 40 sessions, 16 animals). Right: Preferred direction of pre-synaptic grid cells and direction of grid phase offset between pre- and postsynaptic cell for all pairs of putatively connected conjunctive grid cells and ‘pure’ grid cells (angle: -0.7 ± 64.1 deg, correlation: r=0.35 p<0.001, n=217 pairs from 40 sessions). Each dot corresponds to one pair of cells. **g.** Left: Tuning directions for randomly selected pairs of non-connected ID cells conjunctive grid cells (1,600 pairs, 100 pairs per animal). Absolute angles between tuning directions were significantly smaller for connected ID → conjunctive grid pairs than for randomly selected cell pairs (mean absolute offset: 42.0 deg vs. 84.6 deg (chance 90 deg), p=7.4e-17, Mann-Whitney U test). Right: Tuning directions for randomly selected pairs of non-connected conjunctive grid cells and pure grid cells (1,600 pairs, 100 pairs per animal). Absolute angles between preferred directions were significantly smaller for connected conjunctive grid → pure grid cell pairs than for randomly selected cell pairs (mean absolute offset: 34.1 deg vs. 91.5 deg, p = 1.5e-11, Mann-Whitney U test). **h.** Connection rates between functional cell types estimated with different significance thresholds for identification of monosynaptic connections. Thresholds are specified in terms of standard deviations from the baseline firing-rate cross-correlograms. While connection probabilities are heavily dependent on detection thresholds (average connection probability ranging from 0.04% to 0.4%), the connection probability ratios are fairly stable. **i.** Alignment of tuning relationships of connected cell pairs is stable across significance thresholds for detecting monosynaptic connections. Left: directional tuning in ID → conjunctive grid cell pairs (mean angles: 12.7–16.3 deg, correlation coefficients 0.35–0.68); right: directional-tuning vs. grid-phase offset angle in conjunctive grid → pure grid cell pairs (mean angles between preferred directions: -2.0 to -0.4 deg, correlation coefficients 0.34–0.79).

**Extended Data Fig. 7:**
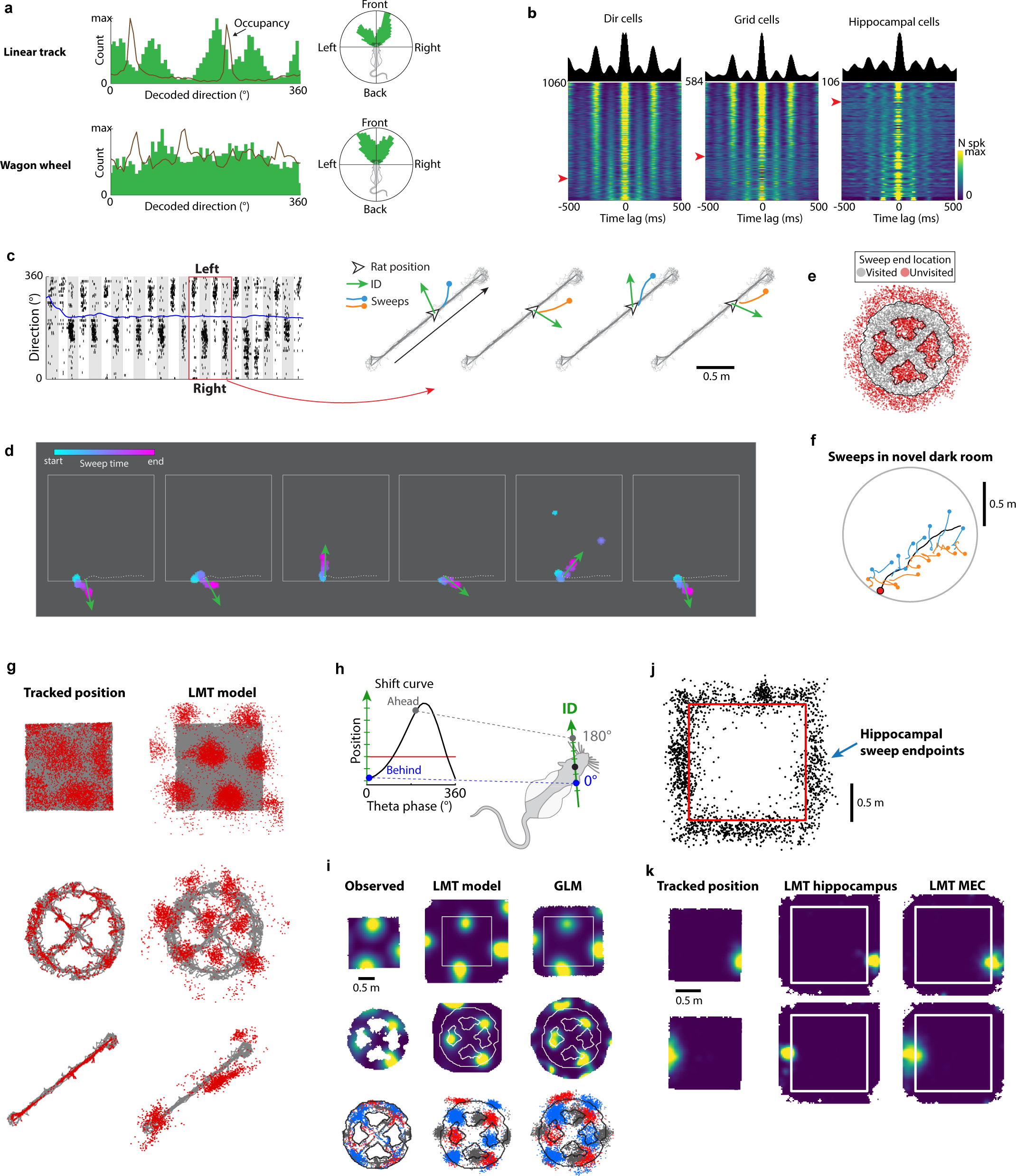
Sweeps extend into never-visited space and persist in novel environments and during darkness. **a.** Left: Histogram of decoded directions (green bars) and head-direction occupancy (solid line) during an example linear-track session (top) and wagon-wheel session (bottom). Although the occupancy on the linear track is biased to the axes of the track (visible as peaks), the sampling of other angles was sufficient for subsequent decoding of the direction to either side of the track. Right: Circular histogram of decoded directions in head-centered coordinates. Note that the internal direction signal is bimodally distributed around the head axis also in 1-D environments. **b.** Theta cycle skipping during linear-track running. Heat maps show firing-rate temporal autocorrelograms (±500 ms) from all theta-modulated direction-tuned cells, grid cells and hippocampal cells (left to right). Each row corresponds to one cell; cells are sorted by theta-skipping index. Cells above the red arrowhead have positive scores and are thus classified as ‘skipping’ cells. One example autocorrelogram from a skipping cell is shown above each plot. The presence of cycle skipping in this task indicates representation of alternating directions and locations, incompatible with coding for the running path, but compatible with representation of unvisited space on either side of the track. **c.** Alternating direction signals and sweeps during linear-track running extracted from fitted latent variables. Left: Raster plot showing spike times of internal direction cells (sorted by preferred internal direction) during a lap on the linear track. Right: Decoded sweeps (lines) and internal direction (arrows) during four theta cycles of the lap (indicated by red square in left panel). Note that sweep and direction signals point to the same side of the track in an alternating pattern. **d.** Decoded direction and sweeps from consecutive theta cycles during a period of running along one of the open-field walls (white square). Sweeps and decoded direction are plotted as in Fig. 4d, with color indicating time within sweep. Note that sweeps travel through the opaque walls of the arena, in alignment with the internal direction signal. **e.** Sweeps span a 2D map even when navigation is confined to 1D paths. Scatter plot of sweep terminal positions during a recording session on the WW maze. Note that density of sweep terminal positions is similar for visited and unvisited locations (density of sweep endpoints in visited vs unvisited portions of the area inside the outer circumference of the maze: 51.2 ± 11.8% vs 50.2 ± 4.1%, mean ± s.e.m. across two rats). **f.** Sweeps extracted from the LMT position latent variable during exploration of a novel circular open field during darkness. Alternating sweeps are visible, although the decoded position deviates substantially from the rat’s actual running trajectory (in grey). **g.** More examples of grid-cell tuning to unvisited locations in the three environments (top to bottom: open field, wagon wheel, linear track). Each column corresponds to one cell and shows the position of the rat at the time of each spike (left) or the latent position from the LMT model at the time of each spike (right). **h.** Because the LMT model, like other dimensionality-reduction methods, finds dense representations of the input data, in principle, grid-like tuning could emerge as a close-packing artifact during fitting. As a control, an alternative single-cell model was used to infer out-of-bounds tuning for each cell independently. The activity of each cell was fitted by a GLM-based model in which the animal’s recorded position (black dot) was shifted along the axis parallel to internal direction (‘ID’) as a function of theta phase, according to a “shift curve” which was fitted separately for each cell (see Methods for details). **i.** Firing-rate maps for a grid cell with respect to tracked 2-D position (left), inferred by the LMT model (middle) and by the GLM-based single-cell model (right) during an open field experiment (top row) and on the wagon wheel track (middle row). Each plot in the bottom row shows the positions of spatial receptive fields (colored patches) of three co-recorded cells on the wagon wheel track. Spikes from each cell is plotted with different colors. Note that the latent fields identified by the LMT and GLM models preserve similar phase offsets between the grid cells, as expected based on extrapolation of their grid patterns. **j.** Scatter of hippocampal sweep endpoints during theta cycles where sweeps from co-recorded MEC-parasubiculum cells terminated outside the open field arena (red box). Out-of-bounds hippocampal sweeps were detected in the majority (65.7 ± 2.6%, mean ± s.e.m.) of theta cycles in which simultaneously decoded MEC sweeps terminated outside the open field arena, significantly more often than during theta cycles that preceded or followed a MEC sweep outside the arena (difference in fraction of outside sweeps: 11.6 ± 1.1%, p=0.031, two-tailed Wilcoxon signed-rank test). **k.** More examples of place-cell tuning to locations outside the open field arena. Each row corresponds to one cell. Columns show firing-rate maps with respect to tracked position (left), and fitted position from the LMT model, either fitted to hippocampal activity (middle) or MEC activity (right).

**Extended Data Fig. 8:**
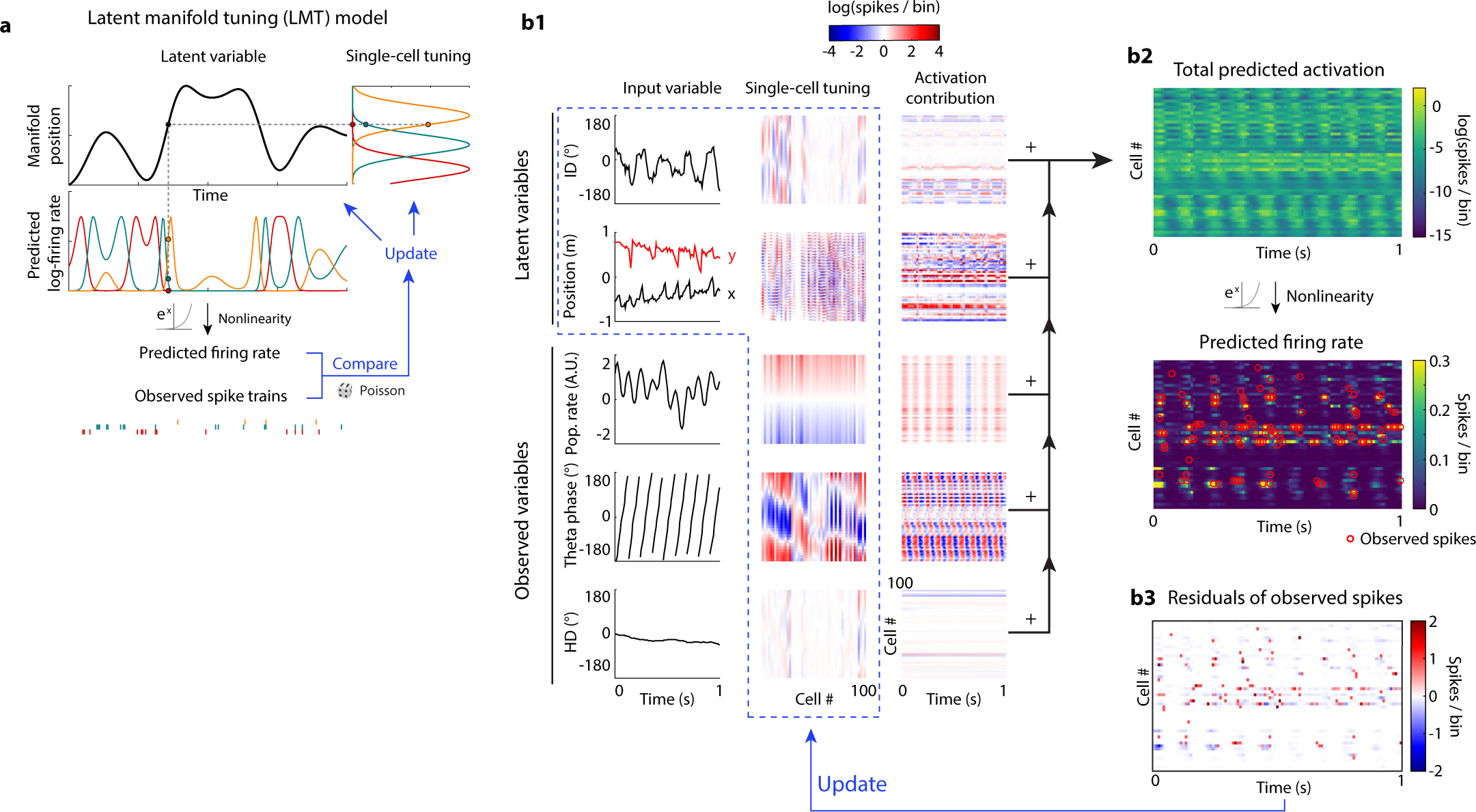
Latent variable model. **a.** Illustration showing the basic principles of using the latent manifold tuning (LMT) model to extract a latent signal from neural population activity. A latent variable (top, black curve) evolves smoothly in time on a one-dimensional manifold. Individual cells are tuned to specific locations on the manifold (top right). At each point in time, the latent variable value predicts each cell’s log-firing rate (second row), which is then transformed with an exponential nonlinearity into a firing-rate prediction. The latter is compared with the observed spikes fired by the cells, treating the spike counts as a Poisson process. The model is learned by iteratively optimizing the latent variable and the tuning curves to improve the prediction of the observed spikes. **b.** Schematic showing the design of the complete “composite” model. **b1**: neural activity is modeled as a function of five input variables (first column). The two latent variables of interest (internal direction (‘ID’) and position; first two rows), are fitted while regressing out contributions of three observed variables known to modulate MEC neural activity (theta phase, HD and population firing rate; three bottom rows). Left column: example traces of the input variables. All variables are assigned with corresponding tuning for each cell (second column; 100 example cells are shown), which, in conjunction with the latent variable’s value, predicts each cell’s log-firing-rate at each time point (third column). The log-firing-rate predictions are linearly summed across all variables (**b2**, top), then the sum is exponentiated, yielding a net prediction of the population firing rates (bottom). For reference, observed spikes for each cell are overlaid (red circles). **b3**: The unaccounted-for (“residual”) neural activity is calculated by subtracting the predicted firing rates from the observed firing. At each iteration of fitting the model, the residuals are used to calculate the next update to the latent variables and the tuning curves (enclosed by blue dashed box), leading to gradual improvement in the match between predicted and observed spiking.

**Extended Data Fig. 9:**
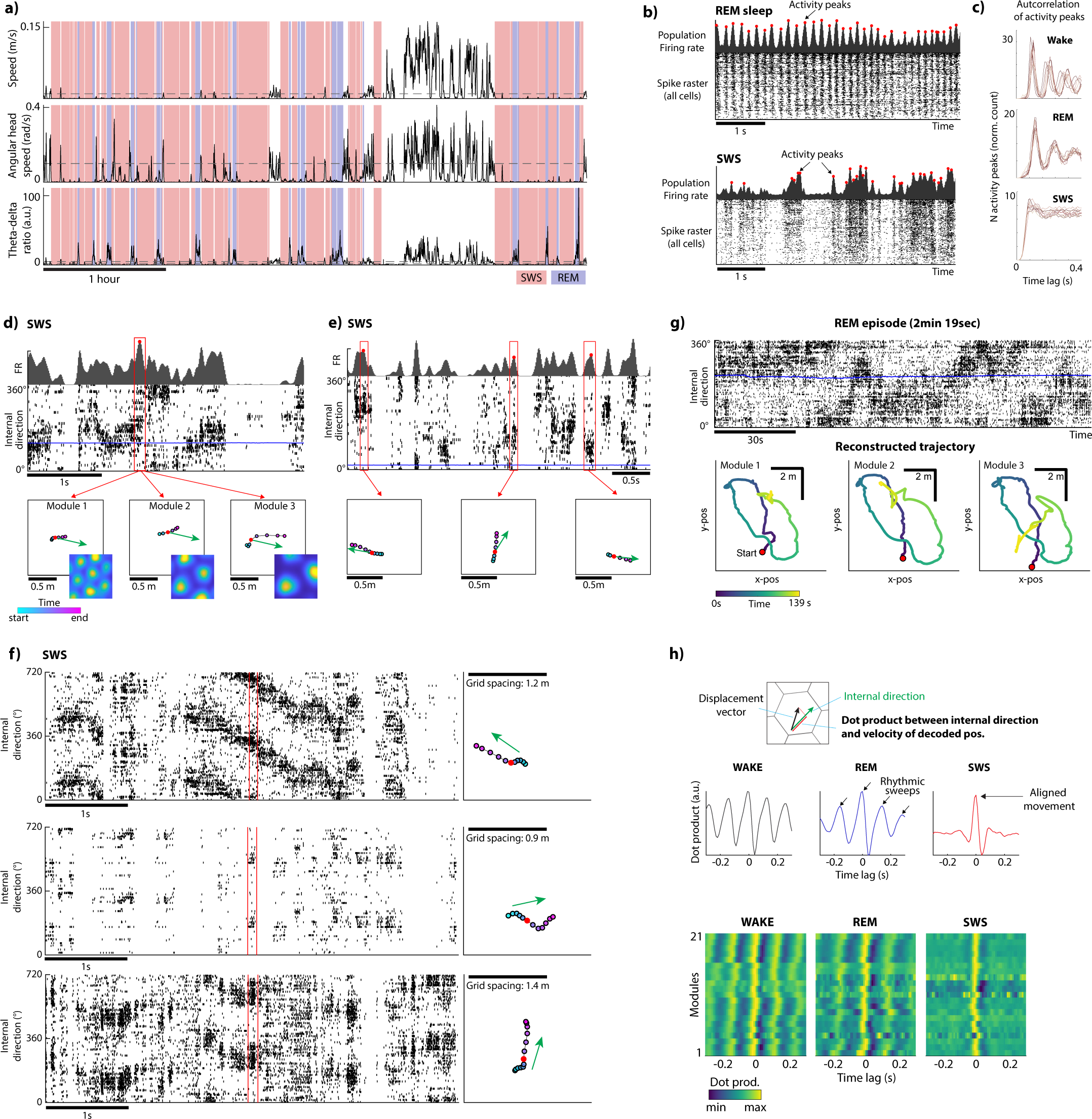
Sleep. **a.** Sleep classification based on movement parameters and electrophysiological signatures, shown here for one example recording. Panels show head speed (top), angular head speed (middle) and theta/delta ratio (bottom) over the course of a recording session (scale bar, 1 h). We used these parameters to identify episodes of slow-wave sleep (SWS; red background shading) and REM sleep (blue background shading). **b.** Spike rasters from all cells recorded simultaneously in MEC-parasubiculum during 5-s epochs of REM sleep (top) and SWS (bottom) in an example animal. Cells are sorted by mean firing rate. Population activity is dominated by theta oscillations during REM and by distinct UP and DOWN-states during SWS. Theta-rhythmic activity peaks can be detected in summed population activity (shown above rasters) during REM, while activity peaks occur at irregular intervals during SWS. **c.** Temporal autocorrelation of activity peaks across brain states. Activity peaks were identified from the summed activity of all direction-tuned cells. Panels show autocorrelation histograms of detected activity peaks during wake, REM and SWS (top to bottom), with counts of activity peaks (y-axis) at different time lags (x-axis). Note that peaks of activity occur at theta-rhythmic intervals during wake and REM, but not during SWS. **d.** Internal direction-aligned, non-rhythmic trajectories during SWS. Top: Summed firing rate of internal direction cells over a ∼3-second SWS epoch. Middle: Sorted population activity of internal direction cells (black ticks) and tracked head direction (solid blue line) during the same period. Note sharp transitions between up and down states (with and without spiking activity, respectively). Bottom: Decoded position (from three simultaneously recorded grid modules; color-coded by time) centered around each of the highlighted peaks of direction-correlated activity (red filled circles in the top panel). Red dots show the decoded position at the time of the activity peaks. Green arrows indicate decoded direction at the time of the activity peaks. Note that the activity burst is accompanied by aligned sweeps in all three modules. **e.** Same as **d** but showing decoded position from a single grid module during three separate peaks of population activity (red filled circles in the top panel). Note that the population activity of direction-tuned cells is discretized in brief bursts and that the internal direction signal often resets between bursts of activity. **f.** More examples of decoded internal direction and sweeps during SWS. Left: Each panel shows spike rasters from internal direction cells (sorted by preferred direction) during a 5 s epoch of SWS activity from three different animals (first examples from same session as in Fig. 5a-b). Right: Each panel shows a decoded sweep and internal direction (same as in **d,e**) during the period highlighted in the corresponding left panel. **g.** Decoded direction and sweep trajectories during a REM sleep episode of 2 min and 19 s. Top: Raster plot showing spike times of internal direction cells during the REM episode, sorted by preferred firing direction during a separate session of open field foraging. Bottom: Decoded position from grid cells during the same REM period based on fitted LMT tuning curves during wake. Position was decoded separately from grid cells belonging to each of three simultaneously recorded grid modules based on tuning curves from the open field session. Trajectories are smoothed in time with a wide gaussian kernel (σ = 100 ms). Each panel shows the decoded trajectory for each of the three grid modules (left to right), color-coded by time. Note that all modules play out similar trajectories (of several meters’ length) over the course of the REM episode (minutes). **h.** Direction-aligned sweeps are rhythmic during wake and REM, but not during SWS. Top: We computed the dot product between two vectors (arrows): internal direction (green) and sweep displacement vectors (black) at a range of time lags relative to directional activity peaks. The dot product is the length of the projection of one vector onto the other (red line). Middle: Dot product across brain states for an example grid module. The dot product is positive at 0-lag during all states, indicating that sweeps and internal direction move synchronously in alignment. Note that the dot product oscillates at theta frequency during wake and REM (arrows), meaning that the grid module expresses rhythmic, direction-aligned trajectories that reset on every theta cycle. Direction-aligned trajectories are also present during SWS (positive dot product at zero lag), but they are not rhythmic. Bottom: Dot product (color-coded) as a function of time lag for all grid modules across animals. Each row shows one grid module (21 grid modules from 9 animals).

**Extended Data Fig. 10:**
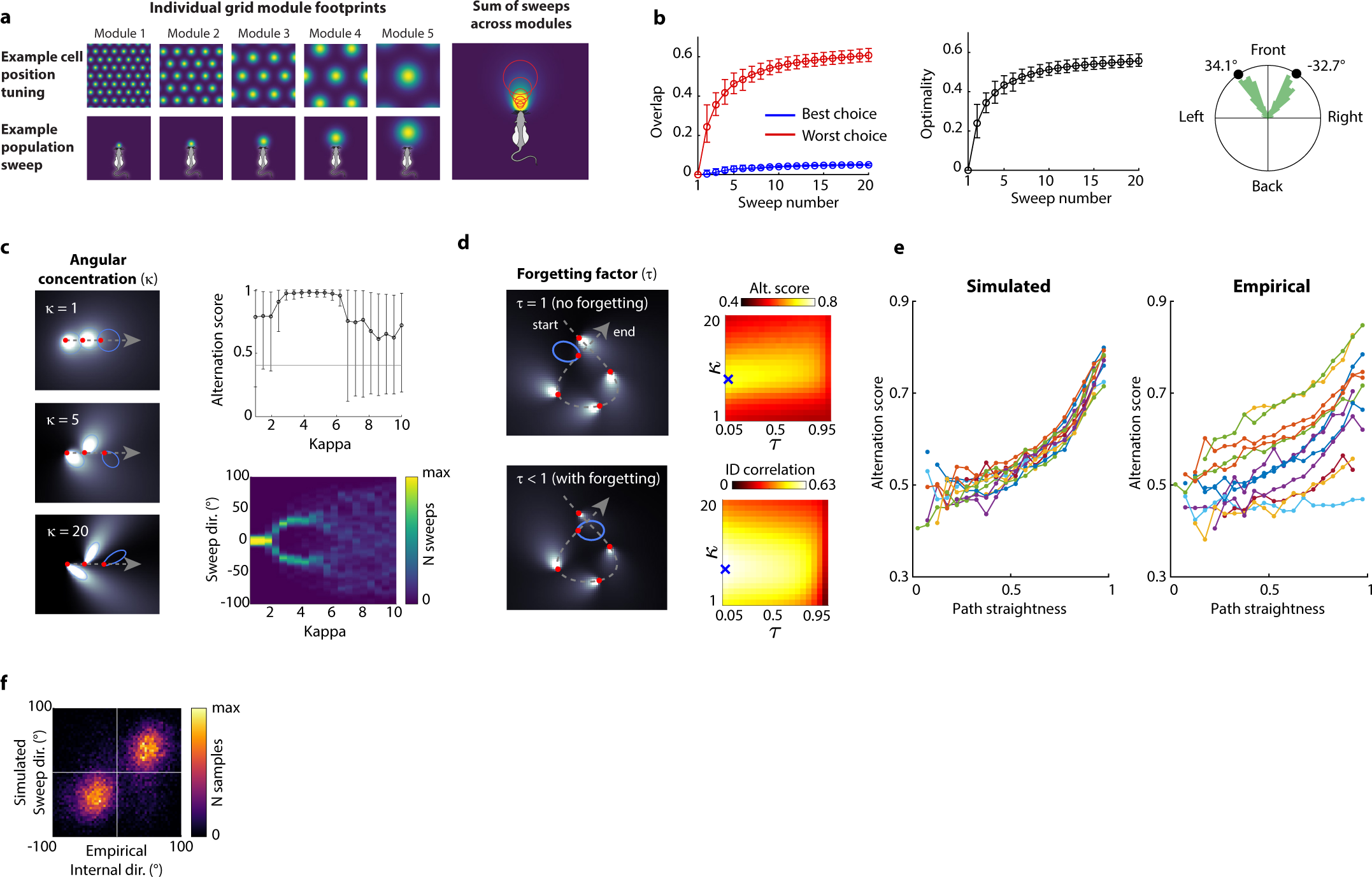
Left-right alternation is a stable regime for efficient spatial sampling. **a.** The sum of sweeps across grid modules has a characteristic beam-like spatial footprint which expands and decays with distance. Left: simulated spatial firing patterns (top row) and sweep footprints (bottom row) for five simulated grid-cell modules. During the simulation, the agent is tasked with placing this footprint in the direction that minimizes overlap with previous footprints, considering all possible directions. Note the proportionality between the grid field size, grid inter-field spacing, sweep footprint size and sweep distance. Right: total sweep footprint yielded by combining the five module footprints, weighted such that each module makes the same total contribution. Note how the intensity decays and broadens with distance, like a torch beam. **b.** Optimality and bimodality of the agent’s choices in 1,000 runs of the simulation in Fig. 6a. Left: average value of the largest (“worst”) and smallest (“best”) sweep overlap values at each time step, considering all possible sweep directions. The overlap value for a sweep at a given position and direction is defined as the product of the preexisting coverage trace with the footprint of the current sweep. Error bars indicate the 5^th^ and 95^th^ percentiles. Middle: optimality at each time step, defined as the difference between the best and the worst choice. Note that all metrics appear to reach convergence before the end of the simulation. Right: Circular histogram of optimal sweep directions (smallest possible overlap with previous sweeps), relative to the direction of movement (“front”). **c.** Alternation is stable within a range of sweep widths. Left: the angular concentration of the sweep is set by the Von Mises distribution parameter κ. Top right: Alternation score (mean and 5^th^-95^th^ percentiles) for repeated simulations across a range of sweep widths. Bottom right: Color-coded histograms (columns) of optimal sweep directions for different sweep widths. Note the bimodality of sweep directions for an intermediate range of sweep widths. **d.** Rapid forgetting yields consistent alternation in the open field. Left: the temporal decay of the cumulative coverage trace is set by the forgetting factor τ. When τ < 1 (bottom) the penalty trace fades over time, meaning that the agent is less influenced by earlier sweeps. The direction of the final sweep (blue ellipse) changes depending on how much the penalty from the first sweep has decayed. Right panels: Exploration of κ and τ parameter space in simulations where the agent was tasked with choosing sweep directions based on the recorded behavioral trajectories of each rat. Top right: average alternation score of the agent’s chosen sweep directions, for each combination of κ and τ, based on the running trajectory of an animal during a recording in the open field (same session as Fig 6c-e). Bottom right: circular correlation coefficient (‘ID correlation’) between decoded internal direction (ID) and the simulated sweep directions when the agent was run in ‘hybrid’ mode (as illustrated in Fig. 6c). Blue crosses indicate the position of the maximum. Note that rapid forgetting and intermediate sweep widths results in robust alternation (τ: 0.039 ± 0.0012, κ: 3.1 ± 0.01, mean ± s.e.m. across 12 animals) and high correlation with empirical sweep directions (τ: 0.01 ± 0.00, κ: 2.3 ± 0.12, mean ± s.e.m. across 12 animals). **e.** Alternation of sweep directions increases with path straightness. Path straightness was computed for each time step of the recording by dividing net travel distance by cumulative traveled distance within a 2 s moving window (high values correspond to straight paths). Left: Average alternation score of the angles chosen by the agent during an open field session, binned at different levels of path straightness. Each set of connected dots (with similar color) shows data from a different animal (n=12). Right: same, but for empirical decoded directions from the same animals. **f.** The agent accurately predicts empirical sweep directions. Heat map of decoded direction and predicted sweep directions (both in egocentric coordinates, relative to head axis) showing a high degree of correspondence between empirical and predicted sweep directions.

## Supplementary Videos

### Supplementary Video 1

Video showing decoded position (Bayesian decoding from LMT tuning curves) from all recorded MEC-parasubiculum cells during an epoch of running in the open field (same example as Fig. 1a; slowed down 15x). Decoded position throughout each theta cycle is plotted as colored blobs with color indicating time within sweep (reset at the beginning of each sweep) and color intensity indicating decoding probability (color range from zero to the maximal posterior probability value of each frame). Note that the decoded position sweeps outwards in a left-right-alternating pattern across theta cycles.

### Supplementary Video 2

Video showing decoded position (Bayesian decoding from LMT tuning curves) from three simultaneously recorded grid modules during an epoch of running in the open field (same example as Fig. 1e; slowed down 15x). Decoded position throughout each theta cycle is plotted as colored blobs (position bins where decoding probability exceeded the 90th percentile of probability values in each frame). Note coordinated left-right alternation and the relationship between the grid spacing and sweep trajectory length.

### Supplementary Video 3

Video showing decoded position (based on LMT position rate maps, as in Video 1) and decoded direction (based on LMT internal direction tuning curves) from all recorded MEC-parasubiculum cells during an epoch of running in the open field (same example as Fig. 1a; slowed down 15x). Decoded ‘internal’ direction is plotted as a green arrow with length scaled to the population firing rate. Note that the decoded direction points in the same direction as sweeps.

## References

1. O’Keefe, J. & Dostrovsky, J. The hippocampus as a spatial map. Preliminary evidence from unit activity in the freely-moving rat. Brain Res. 34, 171–175 (1971).

2. O’Keefe, J. & Nadel, L. *The Hippocampus as a Cognitive Map*. (Oxford University Press, 1978).

3. Wilson, M. A. & McNaughton, B. L. Dynamics of the Hippocampal Ensemble Code for Space. Science 261, 1055–1058 (1993).

4. Hafting, T., Fyhn, M., Molden, S., Moser, M.-B. & Moser, E. I. Microstructure of a spatial map in the entorhinal cortex. Nature 436, 801–806 (2005).

5. Gardner, R. J. et al. Toroidal topology of population activity in grid cells. Nature 602, 123–128 (2022).

6. Skaggs, W. E., McNaughton, B. L., Wilson, M. A. & Barnes, C. A. Theta phase precession in hippocampal neuronal populations and the compression of temporal sequences. Hippocampus 6, 149– 172 (1996).

7. Dragoi, G. & Buzsáki, G. Temporal Encoding of Place Sequences by Hippocampal Cell Assemblies. Neuron 50, 145–157 (2006).

8. Foster, D. J. & Wilson, M. A. Hippocampal theta sequences. Hippocampus 17, 1093–1099 (2007).

9. Johnson, A. & Redish, A. D. Neural Ensembles in CA3 Transiently Encode Paths Forward of the Animal at a Decision Point. J. Neurosci. 27, 12176–12189 (2007).

10. Kay, K. et al. Constant Sub-second Cycling between Representations of Possible Futures in the Hippocampus. Cell 180, 552–567.e25 (2020).

11. McNaughton, B. L., Battaglia, F. P., Jensen, O., Moser, E. I. & Moser, M.-B. Path integration and the neural basis of the ‘cognitive map’. Nat. Rev. Neurosci. 7, 663–678 (2006).

12. Moser, E. I. et al. Grid cells and cortical representation. Nat. Rev. Neurosci. 15, 466–481 (2014).

13. Fuhs, M. C. & Touretzky, D. S. A Spin Glass Model of Path Integration in Rat Medial Entorhinal Cortex. J. Neurosci. 26, 4266–4276 (2006).

14. Guanella, A., Kiper, D. & Verschure, P. A model of grid cells based on a twisted torus topology. Int. J. Neural Syst. 17, 231–240 (2007).

15. Burak, Y. & Fiete, I. R. Accurate Path Integration in Continuous Attractor Network Models of Grid Cells. PLoS Comput. Biol. 5, (2009).

16. Stensola, H. et al. The entorhinal grid map is discretized. Nature 492, 72–78 (2012).

17. Yoon, K. et al. Specific evidence of low-dimensional continuous attractor dynamics in grid cells. Nat. Neurosci. 16, 1077–1084 (2013).

18. Trettel, S. G., Trimper, J. B., Hwaun, E., Fiete, I. R. & Colgin, L. L. Grid cell co-activity patterns during sleep reflect spatial overlap of grid fields during active behaviors. Nat. Neurosci. 22, 609–617 (2019).

19. Gardner, R. J., Lu, L., Wernle, T., Moser, M.-B. & Moser, E. I. Correlation structure of grid cells is preserved during sleep. Nat. Neurosci. 22, 598–608 (2019).

20. Fiete, I. R., Burak, Y. & Brookings, T. What Grid Cells Convey about Rat Location. J. Neurosci. 28, 6858–6871 (2008).

21. Kubie, J. & Fenton, A. Linear Look-Ahead in Conjunctive Cells: An Entorhinal Mechanism for Vector-Based Navigation. Front. Neural Circuits 6, (2012).

22. Erdem, U. M. & Hasselmo, M. A goal-directed spatial navigation model using forward trajectory planning based on grid cells. Eur. J. Neurosci. 35, 916–931 (2012).

23. Erdem, U. M. & Hasselmo, M. E. A Biologically Inspired Hierarchical Goal Directed Navigation Model. J. Physiol. Paris 108, 28–37 (2014).

24. Sanders, H., Rennó-Costa, C., Idiart, M. & Lisman, J. Grid Cells and Place Cells: An Integrated View of their Navigational and Memory Function. Trends Neurosci. 38, 763–775 (2015).

25. Navratilova, Z., Giocomo, L. M., Fellous, J.-M., Hasselmo, M. E. & McNaughton, B. L. Phase precession and variable spatial scaling in a periodic attractor map model of medial entorhinal grid cells with realistic after-spike dynamics. Hippocampus 22, 772–789 (2012).

26. Jun, J. J. et al. Fully integrated silicon probes for high-density recording of neural activity. Nature 551, 232–236 (2017).

27. Steinmetz, N. A. et al. Neuropixels 2.0: A miniaturized high-density probe for stable, long-term brain recordings. Science (2021) doi:10.1126/science.abf4588.

28. Green, J. D. & Arduini, A. A. Hippocampal electrical activity in arousal. J. Neurophysiol. 17, 533– 557 (1954).

29. Mitchell, S. J. & Ranck, J. B. Generation of theta rhythm in medial entorhinal cortex of freely moving rats. Brain Res. 189, 49–66 (1980).

30. Buzsáki, G., Lai-Wo S., L. & Vanderwolf, C. H. Cellular bases of hippocampal EEG in the behaving rat. Brain Res. Rev. 6, 139–171 (1983).

31. Latuske, P., Toader, O. & Allen, K. Interspike Intervals Reveal Functionally Distinct Cell Populations in the Medial Entorhinal Cortex. J. Neurosci. 35, 10963–10976 (2015).

32. Newman, E. L. & Hasselmo, M. E. Grid cell firing properties vary as a function of theta phase locking preferences in the rat medial entorhinal cortex. Front. Syst. Neurosci. 8, (2014).

33. Barry, C., Hayman, R., Burgess, N. & Jeffery, K. J. Experience-dependent rescaling of entorhinal grids. Nat. Neurosci. 10, 682–684 (2007).

34. Hafting, T., Fyhn, M., Bonnevie, T., Moser, M.-B. & Moser, E. I. Hippocampus-independent phase precession in entorhinal grid cells. Nature 453, 1248–1252 (2008).

35. Taube, J. S., Muller, R. U. & Ranck, J. B. Head-direction cells recorded from the postsubiculum in freely moving rats. I. Description and quantitative analysis. J. Neurosci. 10, 420–435 (1990).

36. Sargolini, F. et al. Conjunctive Representation of Position, Direction, and Velocity in Entorhinal Cortex. Science 312, 758–762 (2006).

37. Boccara, C. N. et al. Grid cells in pre- and parasubiculum. Nat. Neurosci. 13, 987–994 (2010).

38. Taube, J. S. Head direction cells recorded in the anterior thalamic nuclei of freely moving rats. J. Neurosci. 15, 70–86 (1995).

39. Jeffery, K. J., Donnett, J. G. & O’Keefe, J. Medial septal control of theta-correlated unit firing in the entorhinal cortex of awake rats: NeuroReport 6, 2166–2170 (1995).

40. Deshmukh, S. S., Yoganarasimha, D., Voicu, H. & Knierim, J. J. Theta Modulation in the Medial and the Lateral Entorhinal Cortices. J. Neurophysiol. 104, 994–1006 (2010).

41. Brandon, M. P., Bogaard, A. R., Schultheiss, N. W. & Hasselmo, M. E. Segregation of cortical head direction cell assemblies on alternating theta cycles. Nat. Neurosci. 16, 739–748 (2013).

42. Köhler, C. Intrinsic projections of the retrohippocampal region in the rat brain. I. The subicular complex. J. Comp. Neurol. 236, 504–522 (1985).

43. Perkel, D. H., Gerstein, G. L. & Moore, G. P. Neuronal Spike Trains and Stochastic Point Processes: II. Simultaneous Spike Trains. Biophys. J. 7, 419–440 (1967).

44. Csicsvari, J., Hirase, H., Czurko, A. & Buzsáki, G. Reliability and State Dependence of Pyramidal Cell–Interneuron Synapses in the Hippocampus: an Ensemble Approach in the Behaving Rat. Neuron 21, 179–189 (1998).

45. 45. Wu, A., Roy, N. A., Keeley, S. & Pillow, J. W. Gaussian process based nonlinear latent structure discovery in multivariate spike train data. in Advances in Neural Information Processing Systems vol. 30 (Curran Associates, Inc., 2017).

46. Peyrache, A., Lacroix, M. M., Petersen, P. C. & Buzsáki, G. Internally organized mechanisms of the head direction sense. Nat. Neurosci. 18, 569–575 (2015).

47. Chaudhuri, R., Gerçek, B., Pandey, B., Peyrache, A. & Fiete, I. The intrinsic attractor manifold and population dynamics of a canonical cognitive circuit across waking and sleep. Nat. Neurosci. 22, 1512– 1520 (2019).

48. Muller, R. U., Kubie, J. L. & Saypoff, R. The hippocampus as a cognitive graph (abridged version). Hippocampus 1, 243–246 (1991).

49. Samsonovich, A. & McNaughton, B. L. Path Integration and Cognitive Mapping in a Continuous Attractor Neural Network Model. J. Neurosci. 17, 5900–5920 (1997).

50. Yovel, Y., Falk, B., Moss, C. F. & Ulanovsky, N. Optimal Localization by Pointing Off Axis. Science 327, 701–704 (2010).

51. O’Keefe, J. & Recce, M. L. Phase relationship between hippocampal place units and the EEG theta rhythm. Hippocampus 3, 317–330 (1993).

52. Wikenheiser, A. M. & Redish, A. D. Hippocampal theta sequences reflect current goals. Nat. Neurosci. 18, 289–294 (2015).

53. Tang, W., Shin, J. D. & Jadhav, S. P. Multiple time-scales of decision-making in the hippocampus and prefrontal cortex. eLife 10, e66227 (2021).

54. Kiehn, O. LOCOMOTOR CIRCUITS IN THE MAMMALIAN SPINAL CORD. Annu. Rev. Neurosci. 29, 279–306 (2006).

55. Feldman, J. L., Negro, C. A. D. & Gray, P. A. Understanding the Rhythm of Breathing: So Near, Yet So Far. Annu. Rev. Physiol. 75, 423–452 (2013).

56. Fenk, L. A., Riquelme, J. L. & Laurent, G. Interhemispheric competition during sleep. Nature 616, 312–318 (2023).

57. Brown, T. G. The Intrinsic Factors in the Act of Progression in the Mammal. Proc. R. Soc. Lond. Ser. B Contain. Pap. Biol. Character 84, 308–319 (1911).

58. Grillner, S. & Zangger, P. On the central generation of locomotion in the low spinal cat. Exp. Brain Res. 34, 241–261 (1979).

59. Smith, J. C., Ellenberger, H. H., Ballanyi, K., Richter, D. W. & Feldman, J. L. Pre-Bötzinger complex: a brainstem region that may generate respiratory rhythm in mammals. Science 254, 726–729 (1991).

60. Yuste, R., MacLean, J. N., Smith, J. & Lansner, A. The cortex as a central pattern generator. Nat. Rev. Neurosci. 6, 477–483 (2005).

61. Adrian, E. D. & Zotterman, Y. The impulses produced by sensory nerve endings. J. Physiol. 61, 465– 483 (1926).

62. Hopfield, J. J. Neurodynamics of mental exploration. Proc. Natl. Acad. Sci. 107, 1648–1653 (2010).

63. Chu, T. et al. Firing rate adaptation affords place cell theta sweeps, phase precession and procession. eLife 12, (2024).

## Methods references

64. Waaga, T. et al. Grid-cell modules remain coordinated when neural activity is dissociated from external sensory cues. Neuron 110, 1843–1856.e6 (2022).

65. Luo, T. Z. et al. An approach for long-term, multi-probe Neuropixels recordings in unrestrained rats. eLife 9, e59716 (2020).

66. Stark, E. & Abeles, M. Unbiased estimation of precise temporal correlations between spike trains. J. Neurosci. Methods 179, 90–100 (2009).

67. English, D. F. et al. Pyramidal Cell-Interneuron Circuit Architecture and Dynamics in Hippocampal Networks. Neuron 96, 505–520.e7 (2017).

68. Spivak, L., Levi, A., Sloin, H. E., Someck, S. & Stark, E. Deconvolution improves the detection and quantification of spike transmission gain from spike trains. *Commun*. Biol. 5, 1–17 (2022).

69. McInnes, L., Healy, J., Saul, N. & Großberger, L. UMAP: Uniform Manifold Approximation and Projection. J. Open Source Softw. 3, 861 (2018).

70. Zhang, K., Ginzburg, I., McNaughton, B. L. & Sejnowski, T. J. Interpreting Neuronal Population Activity by Reconstruction: Unified Framework With Application to Hippocampal Place Cells. J. Neurophysiol. 79, 1017–1044 (1998).

71. Muller, R. U. & Kubie, J. L. The effects of changes in the environment on the spatial firing of hippocampal complex-spike cells. J. Neurosci. 7, 1951–1968 (1987).

72. Fyhn, M., Hafting, T., Treves, A., Moser, M.-B. & Moser, E. I. Hippocampal remapping and grid realignment in entorhinal cortex. Nature 446, 190–194 (2007).

73. Alme, C. B. et al. Place cells in the hippocampus: Eleven maps for eleven rooms. Proc. Natl. Acad. Sci. 111, 18428–18435 (2014).

74. Hardcastle, K., Maheswaranathan, N., Ganguli, S. & Giocomo, L. M. A Multiplexed, Heterogeneous, and Adaptive Code for Navigation in Medial Entorhinal Cortex. Neuron 94, 375–387.e7 (2017).

75. Wolf, F. A., Angerer, P. & Theis, F. J. SCANPY: large-scale single-cell gene expression data analysis. Genome Biol. 19, 15 (2018).

76. Berens, P. CircStat: A MATLAB Toolbox for Circular Statistics. J. Stat. Softw. 31, 1–21 (2009).

77. Mizuseki, K., Sirota, A., Pastalkova, E. & Buzsáki, G. Theta Oscillations Provide Temporal Windows for Local Circuit Computation in the Entorhinal-Hippocampal Loop. Neuron 64, 267–280 (2009).

